# Genome-wide association studies and genomic selection assays made in a large sample of cacao (*Theobroma cacao* L.) germplasm reveal significant marker-trait associations and good predictive value for improving yield potential

**DOI:** 10.1101/2021.11.22.469505

**Authors:** Frances L. Bekele, Gillian G. Bidaisee, Mathilde Allegre, Xavier Argout, Olivier Fouet, Michel Boccara, Duraisamy Saravanakumar, Isaac Bekele, Claire Lanaud

## Abstract

A genome-wide association study was undertaken to unravel marker-trait associations (MTAs) between SNP markers and yield-related traits. It involved a subset of 421 cacao accessions from the large and diverse collection conserved *ex situ* at the International Cocoa Genebank Trinidad. An average linkage disequilibrium (r^2^) of 0.10 at 5.2 Mb was found across several chromosomes. Seventeen significant (*P* ≤ 8.17 × 10^-5^ (–log10 (p) = 4.088)) MTAs of interest, which accounted for 5 to 17% of the explained phenotypic variation, were identified using a Mixed Linear Model in TASSEL version 5.2.50. The most significant MTAs identified were related to seed number and seed length on chromosome 7 and seed number on chromosome 1. Other significant MTAs involved seed length to width ratio on chromosomes 3 and 5 and seed length on chromosomes 4 and 9. It was noteworthy that several yield-related traits, *viz*., seed length, seed length to width ratio and seed number were associated with markers on different chromosomes, indicating their polygenic nature. Approximately 40 candidate genes that encode embryo and seed development, protein synthesis, carbohydrate transport and lipid biosynthesis and transport were identified in this study. A significant association of fruit surface anthocyanin intensity co-localised with MYB-related protein 308 on chromosome 4. Testing of a genomic selection approach revealed good predictive value (GEBV) for economic traits such as seed number (GEBV = 0.611), seed length (0.6199), seed width (0.5435), seed length to width ratio (0.5503), seed/cotyledon mass (0.6014) and ovule number (0.6325). The findings of this study could facilitate genomic selection and marker-assisted breeding of cacao thereby expediting improvement in the yield potential of cacao planting material.

## Introduction

Cacao, *Theobroma cacao* L., Malvaceae *sensu lato* [1], is an important Neotropical, perennial crop, on which the thriving global cocoa and chocolate industry is based. The World Cocoa Foundation, in 2012, reported that 40-50 million people worldwide depend on cocoa for their livelihood. The global export value of cocoa beans has been fluctuating between 8 billion USD and 10.5 billion USD over the past decade. The global chocolate market was valued at USD 106.6 billion in 2019 [2].

*T. cacao* L. is a diploid (2n = 10), allogamous species. Its genome is small; reported by Argout et al. [3] to span 411-494 Mb. Its putative centre of genetic diversity is at the headwaters of the Amazon River, South America [4], and it is indigenous to the Amazon and Orinoco rainforests. Currently, the majority of cacao cultivation, concentrated in West Africa, is still based on traditional varieties collected in South America prior to 1950 [5, 6]. There is much scope for enhancement of cacao planting material towards realizing the full potential of the crop. The significant progress made in cacao genomics in the last decade [7] should accelerate progress in improvement of cacao planting material.

Phenotypic characterisation and evaluation of cacao, based on conventional techniques, provide a relatively inexpensive and easy method of selection of superior genotypes where attributes are easy to score by observation of morphology or simple screening procedures. However, polygenic traits with significant environmental effects on their expression are more difficult to score without a tool for identifying and following the major genes influencing the phenotypes. For these, marker-assisted and genomic selections are advantageous [8]. Some traits of economic importance in cacao, such as yield and resistance to Black Pod disease, have been characterized as polygenic [9, 10], and it is thus desirable to tag these traits with molecular markers to facilitate early selection for the desired genotype. This involves the identification of quantitative trait loci (QTLs) [11] and the establishment of linkage maps [12-19, 10, 20-28, 11, 3, 29-33].

Fouet et al. [32] mapped approximately one Simple Sequence Repeat (SSR) tag for every 2 cM on the 10 linkage groups of *T. cacao* L. This set of Expressed Sequence Tag SSRs included 14 candidate genes for disease resistance and quality traits. The development of a genomic resource database [34] was crucial in the construction of a high-density gene map for *T. cacao* L. Cacao’s phylogenetic proximity to *Arabidopsis*, which has a well-described gene ontology, was useful for elucidating some of the metabolic pathways investigated by Argout et al. [34]. The sequencing of the cacao genome [3, 29] should facilitate the identification of candidate genes for traits of interest. Version 2.0 of the Criollo genome of Argout et al. [29] has 99% of the assembly anchored to the 10 chromosomes of the *T. cacao* L. genome. This will assist researchers in more easily developing superior cacao plants with disease and pest resistance, high yield, desirable flavour, favourable flavanol (antioxidant) content, self-compatibility [7], and other traits of economic interest or with potential health benefits [35].

The availability of robust phenotypic data for hundreds of accessions at the International Cocoa Genebank Trinidad (ICGT) [36–39] along with whole reference genome sequences for cacao allow us to fully explore genetic diversity in a large and diverse cacao collection and its relationship to phenotypic diversity, as outlined by Varshney et al. [40]. Since there is tight coverage with molecular markers along the cacao genome [3, 29, 41, 30], admixture studies and genome-wide association studies are facilitated.

Genome-wide association studies (GWAS) entail detecting significant associations between individual genetic markers, such as Single Nucleotide Polymorphisms (SNPs), linked to functional alleles from a dense genome-wide panel, with the phenotypic traits of a group of individuals [42, 43]. Population-wide associations may be detected between SNPs and causal polymorphisms (*viz*., QTL) that affect traits of interest such as yield. The genetic profiles of detected superior and other variants can be used for genomic prediction and selection [44, 45]. GWAS also involves searching for genotype-phenotype correlations in unrelated individuals [46, 47], and is based on non-random association of alleles in a population or linkage disequilibrium (LD) [48, 49].

Classical QTL bi-parental mapping (linkage) studies have demonstrated that some molecular markers explain a considerable amount of phenotypic variance in quantitative traits [47], but are constrained by the “paucity of large productive progenies from known parental origin for perennial crops” such as cacao [6, 50]. An advantage of GWAS over classical QTL studies is attributed to the fact that with cumulated cycles of recombination at the population level, associations are broken between a genetic factor determining a phenotypic trait and any marker that is not tightly linked to it. GWAS exploits all of the recombination events that have occurred in the evolutionary history of the germplasm under study, which allows a much higher mapping resolution compared with classical QTL mapping [51, 43]. However, factors such as selection, population admixture and family structure may result in spurious associations between phenotypes and a given marker. In this case, the phenotypes are not physically linked to the marker and co-localized genes, but are inferred to be. Robust GWAS models, using population structure as co-variate, allow the identification of mainly authentic (non-spurious) associations [43].

GWAS may have little or no advantage over QTL mapping in cases where LD is extensive [43]. In cross-pollinated crops like *T. cacao* L., where LD has been observed to decay rapidly (within 1–2 Mbps to an r^2^ value (measure of LD) of about 0.1) in wild genotypes [49]), GWAS is expected to have high resolution. In this case, any marker showing a significant association with a trait is expected to be tightly linked to the gene affecting that trait.

GWAS have been used successfully for identifying phenotype-genotype associations for many traits [52–54]. In *Arabidopsis thaliana*, such traits include shade avoidance, heavy metal and salt tolerance, flowering time and life history traits [55, 51]. GWAS have been conducted in cacao [6] and several other crops including rice [56]; maize [57]; wheat [58]; sorghum [59]; barley [60]; rapeseed [61]; soybean [62]; peanut [63] and other plant species [64].

Allegre et al. [33] and Fouet et al. [32] identified and mapped SNPs and SSR markers useful as expressed sequence tags (ESTs) and constructed a high-density genetic map for *T. cacao* L. Both kinds of markers are co-dominant and thus powerful for genetic analysis. Of the 5,246 SNPs screened by Allegre et al. [33], 1536 were found corresponding to genes with putative functions. Of these, 851 SNPs displayed a distinct polymorphic pattern across a selection of cacao germplasm. The latter are SNPs located within a gene expressed sequence and are thus valuable for identifying candidate genes with functional roles in cacao. The average distance between adjacent markers, in the genetic map constructed by Allegre et al. [33], was 0.6 cM. The data are available at http://tropgenedb.cirad.fr.

The objectives of this research were to exploit the naturally occurring genetic variation in a large collection of cacao trees of diverse origin, including wild genotypes, which are conserved *ex situ* at the ICGT to:

1. Facilitate, via GWAS, the identification of SNP markers significantly associated with phenotypic traits (Marker-Trait Associations or MTAs) and putative candidate genes; and
2. Establish predictive values for phenotypic traits of interest, using a genomic selection approach, to examine the efficiency of this breeding strategy to improve cacao yield.

## Materials and Methods

### Germplasm studied

Four hundred and twenty-one (421) cacao accessions, including 263 wild genotypes (collected in the Amazon Basin [36]), were included in this study. Complete phenotypic data were available for 346 of these accessions (S1 Table). They represent 23 “accession groups”, as described by Bekele et al. [36] as well as most of the genetic groups defined by Motamayor et al. [65], which are conserved *ex situ* at the ICGT. Wild cacao types such as those of the AMAZ, GU, IMC, MO, NA, PA, POUND, RB, SCA and SPEC (1-54) accession groups [36], which have evolved over a long period of time, were included to improve the power of detection of associations between SNP markers and phenotypic traits of interest as recommended by Stack et al. [49].

#### Management of germplasm under study

The ICGT is situated at the University Cocoa Research Station, Centeno, Trinidad at an altitude of 15 m above sea level. Shade is provided by trees of *Erythrina* sp. planted 6 m apart, and bananas (*Musa* sp.) placed 4 m apart. The cacao trees are planted 1.8 m apart with typically up to 16 trees per plot for each accession. The soil type is Cunupia fine sandy clay with restricted internal drainage. Over a 30-year period from 1981, the average temperature was 26.3°C. It was 26°C for the period 1961 to 1991. This satisfied the optimal temperature requirements for growing cacao. The mean annual rainfall for 1981 to 2011 was 1945.2 mm, lower than the 2,392 mm recorded for 1961 to 1991 (Trinidad and Tobago Meteorological Office https://www.metoffice.gov.tt). The plants are irrigated as necessary during the dry season (January-June) each year. Regular weeding and pruning of the trees are undertaken. However, disease and pest control are avoided to facilitate scientific monitoring of these conditions. Fertilizers are applied at planting and regularly for only young trees. The trees are maintained within a low input system.

### Phenotypic data collection

Cacao accessions were assessed in terms of 27 flower, fruit and seed traits as described by Bekele et al. and Bekele and Butler (Table 1) [36, 66]. The traits studied were found to be the most discriminative and taxonomically useful descriptors, which avoid redundancy. They were also selected for ease of observation, reliability of scoring, and, in the case of seed characters, agronomic and economic value [37, 38]. Sample collection was done at the ICGT and spanned the period 1992-2012. When possible, the full complement of samples was collected for each accession at a given time, but in most cases, samples were obtained over multiple years during the same season to preclude the effect of seasonality on phenotypic trait expression. The fruits characterized were the products of open pollination. These data are available online in the International Cocoa Germplasm Database (ICGD) (http://www.icgd.rdg.ac.uk/).

**Table 1.** Descriptors used for phenotypic characterisation (n).

| Descriptor | State [sample size] |
| --- | --- |
| 1. Flower, anthocyanin intensity in column of pedicel (FL_PCOL_C) | 1=green, 2=reddish, 3=red [n=10]. |
| 2. Flower, sepal length (mm) [n=10] (FL_SEP_LEN) |  |
| 3. Flower, anthocyanin intensity on ligule (FL_LIG_COL) | 0=absent, 3=slight, 5=intermediate, 7=intense [n=10] |
| 4. Flower, ligule width (mm) [n=10] (FL_LIG_WID) |  |
| 5. Flower, anthocyanin intensity in filament (FL_FIL_COL) | 0=absent, 3=slight, 5=intermediate, 7=intense [n=10] |
| 6. Flower, style length (mm) [n=10] (FL_STY_LEN) |  |
| **7. Flower, ovule number [n=10] (FL_OV_NUM)** |  |
| 8. Fruit, shape (FR_POSH) | 1=oblong, 2=elliptic, 3=obovate, 4=orbicular [n=10], 5=oblate or other (specified). |
| 9. Fruit, basal constriction (FR_PBC) | 0=absent, 1=slight, 2=intermediate, 3=strong, 4=wide shoulder [n=10] |
| 10. Fruit, apex form (FR_PAF) | 1=attenuate, 2=acute, 3=obtuse, 4=rounded, 5=mammillate, 6=indented [n=10] |
| 11. Fruit, surface texture (rugosity or degree of wartiness) (FR_PST) | 0=absent, 3=slight, 5=intermediate, 7=intense [n=10] |
| 12. Fruit, surface anthocyanin intensity in mature ridges (FR_PRCM) | 0=absent, 3=slight, 5=intermediate, 7=intense [n=10] |
| 13. Fruit, ridge disposition (FR_PFD) | 1=equidistant, 2=paired [n=10] |
| 14. Fruit, primary ridge separation (FR_PFS) | 1=slight, 2=intermediate, 3=wide [n=10] |
| **15. Fruit, pod wall hardness [n=10] (FR_HH (qualitative); FR_QHH (quantitative))** | 3= ≤ 1.6 megapascals (MPa), 5 = > 1.6 MPa ≤ 2.0 MPa, 7= > 2.0 MPa |
| 16. Fruit, length (cm) [n=10] (FR_POL) |  |
| 17. Fruit, width (cm) [n=10] (FR_POW) |  |
| 18. Fruit length to width ratio (FR_PODL_W) |  |
| **19. Seed, number [n=10] (FR_BEN)** |  |
| 20. Seed, shape (FR_BES) | 1=oblong 2=elliptic 3=ovate |
| 21. Seed, cotyledon colour (FR_BEC) | 1=white, 2=grey, 3=light purple, 4=medium purple, 5=dark purple, 6=mottled [n=40] |
| 22. Wet bean mass (total) (g) [n=10] (FR_WABW) |  |
| **23. Cotyledon length (cm) [n=20]. (FR_BL)** |  |
| **24. Cotyledon width (cm) [n=20]. (FR_BEW)** |  |
| **25. Cotyledon mass (g) [n=20] (FR_BW)** |  |
| **26. Cotyledon length to width ratio (FR_BENL_W)** |  |
| **27. Pod index [n=10]** | The number of pods/fruits required to produce 1 kg of dried cocoa (1000/ (average dried cotyledon mass × average seed or bean number) |

2 Traits of economic importance are highlighted in bold

#### Yield-related traits in cacao

The yield-related traits under investigation are listed in Table 1. Since seed/cotyledon mass and seed number per fruit have been reported to have moderate to high heritability [67–69], information on pod index (the number of pods/fruits required to produce 1kg of dried cocoa) is particularly useful to breeders. A low pod index is desirable since it is normally associated with large seed size, which is preferred by chocolate manufacturers, and is a reliable indicator of good yield potential. A maximum pod index of 16.5 fruits was a standard set for selection in Trinidad and Tobago [70].

### Collection of genotypic data

SNP markers that provide good coverage of the cacao genome [33] were employed in this study. The selection of 836 SNPs in coding sequences, which displayed significant similarity with known protein sequences, as described by Argout et al. [34], was carried out by Allegre et al. [33]. Illumina SNP genotyping was performed with the Illumina BeadArray platform at the French National Genotyping Centre (CNG, CEA-IG, Evry, France), according to the GoldenGate Assay manufacturer’s protocol. The genotype calling of each marker was verified using reference genotypes and filtered, as described by Argout et al. [34] and Allegre et al. [33]. The QualitySNP pipeline was used for detection of SNPs in the unigenes. All of the SNPs employed for genotyping have been identified in orthologous genes or gene families and this facilitates reference to genetic information, made available via the genome browser, CocoaGen DB (http://cocoa-genome-hub.southgreen.fr/jbrowse).

### Statistical analyses

#### Phenotypic data analysis

Qualitative data such as fruit shape classes were first converted to binary form. The quantitative traits that were found to deviate from normality were log-transformed. Tests of normality, log transformation of data that were not normally distributed, derivation of descriptive statistics and correlation analysis of the collated phenotypic data were performed using Minitab Version 18.

### Genotypic data analysis

#### Determining population structure

Population structure was determined to allow estimation of marker-trait associations without including spurious associations [71]. This was necessary to satisfy the independence assumption under the null hypothesis on which the marker-trait association is based [72]. The Bayesian clustering software, *STRUCTURE*, [72–75, 54] was employed for this purpose. It defined the inferred ancestry of individuals, studied as coefficients of the individuals across sub-populations. Individuals with coefficients of membership of less than 0.7 were classified as admixed. The Q matrix was used to remove associations due to evolution and to keep only data that have close association to the marker trait. The allele frequencies correlated model used Markov Chain Monte Carlo (MCMC) simulations to estimate the group (cluster, *K*) membership of each individual studied, assuming Hardy-Weinberg and linkage equilibrium within groups, random mating within populations and free recombination between loci [72].

Multi-locus genotype data for 200 SNPs, distributed over all 10 chromosomes, with minor allele frequency (MAF) greater than 0.05 and low missing values (less than 10%) were analysed in *STRUCTURE* to describe and visualize population structure, based on allele frequencies of the data. The optimum *K* value that best defined the population structure was identified using the admixture model of ancestry, assuming correlated allele frequencies for *K* = 2 to 15 with 150,000 iterations during the burn-in period, 150,000 Markov Chain Monte Carlo repetitions and 10 independent runs for each genetic sub-population (*K*2-*K*15).

#### Analysis of inferred population structure

The *STRUCTURE* outputs were analysed to infer optimal *K* based on the method described by Pritchard [72]. The optimal *K* was chosen by plotting the log probability of the data, Pr (X | *K*), against a range of *K* values and selecting the one after which the curve formed a plateau, as indicated by the arrow in Fig 1, while also considering the consistency of the groupings across multiple runs with the same *K*. Runs for which the variance was not homogeneous with variances of the other runs with the same *K* value were excluded.

**Fig 1.**
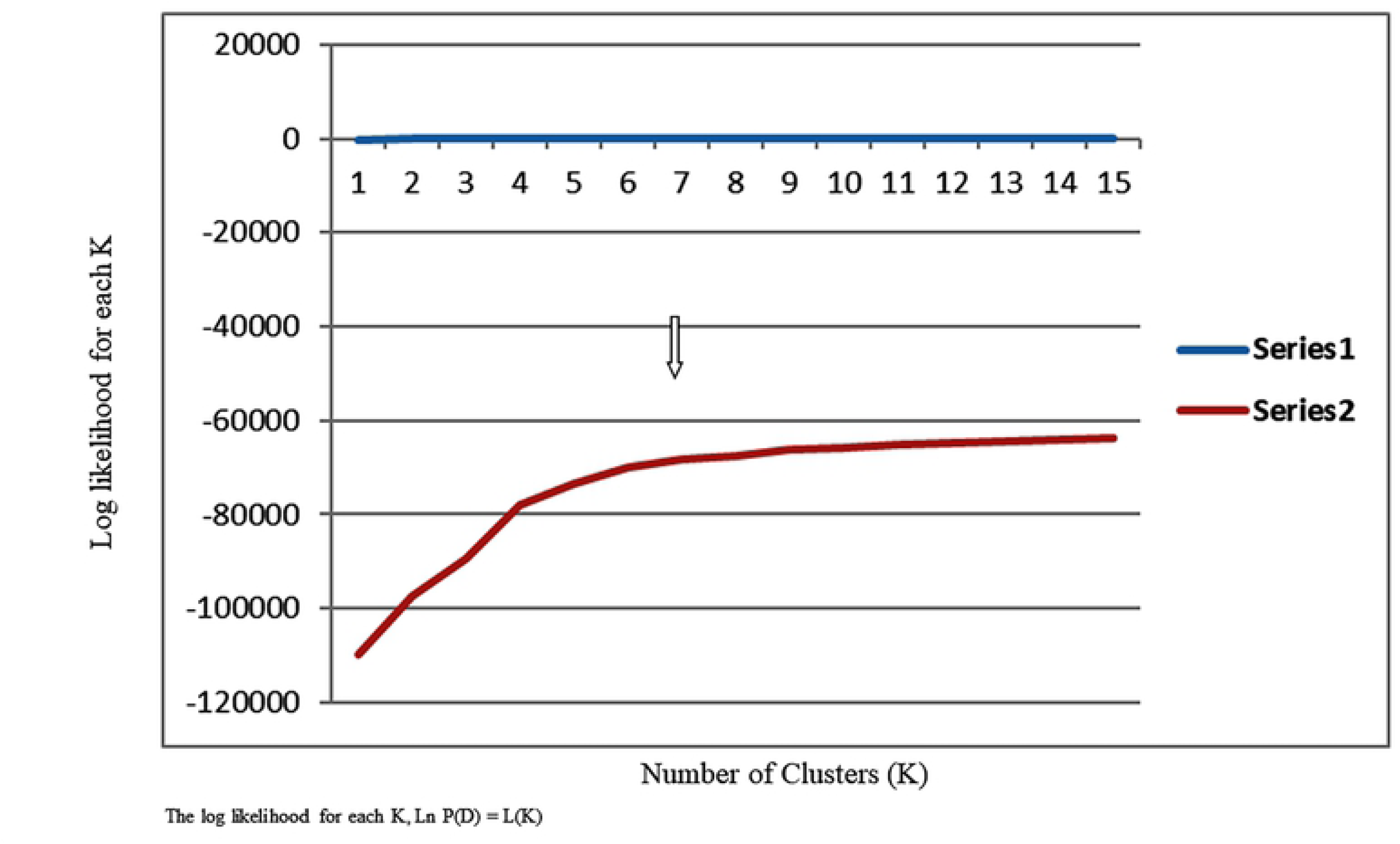
Plot of log of K versus number of clusters based on STRUCTURE analysis. **Legend**: Analysis of population structure of 421 cacao accessions using STRUCTURE - estimated LnP(K) of possible clusters (*K*) from 2 to 15. When *K* is approaching a true value, L(*K*) plateaus (or continues increasing slightly).

The population structure of the 421 accessions studied was visualized using DARwin (Dissimilarity Analysis and Representation for Windows) version 6 (http://darwin.cirad.fr) [76]. DARwin was used to estimate pairwise Jaccard’s genetic dissimilarity indices using the 200 SNP markers employed in *STRUCTURE* Analysis. A tree was constructed by clustering accessions, based on a dissimilarity matrix using the Unweighted Pair Group Method with Arithmetic Mean (UPGMA). Clade strength in the dendrogram was tested using 1000 bootstraps. This tree was rendered with iTOL (https://itol.embl.de/) to produce Fig 2.

**Fig 2.**
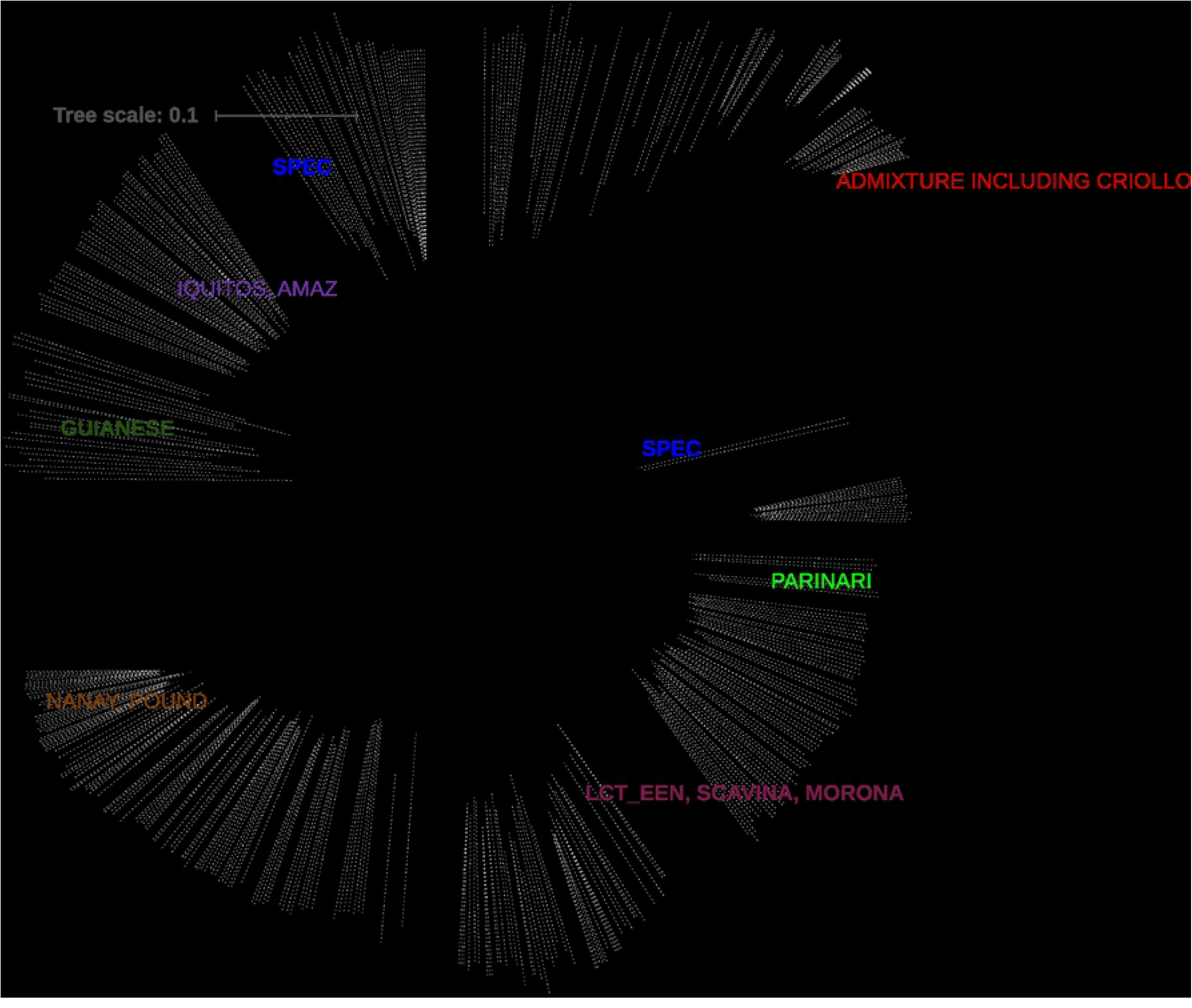
Neighbour-joining tree based on UPGMA of 421 cacao genotypes. **Legend**: The tree was generated in DARwin Version 6 and rendered in iTOL version 6 (https://itol.embl.de/) Bootstrap set at > =90; Seven admixed groups are evident.

### Genome-wide Association Study Analysis (GWAS)

The detection of associations between SNPs and traits is dependent on the phenotypic variance within the population that is explained by the SNP alleles [51]. This variance is determined by the “extent to which the two allelic variants differ in their phenotypic effect (effect size)” in the population under study [51]. *TASSEL* [77], version 5.2.50 for Windows, was employed to conduct GWAS. *TASSEL* employed a fixed effects linear model to test for association between genetic sites and phenotypes. A ‘main effects only’ model was generated using all variables in the input data. Both General Linear (GLM) and the Mixed Linear (MLM) models were used in this study.

MLM was used to correct covariances due to relatedness at the population level between genotypes (due to population structure) [78] as well as co-ancestry (kinship) or identity by descent. The inclusion of the *K* matrix allowed the inclusion of multiple backgrounds QTL as a random factor in the mixed linear model, as explained by Henderson [79]. The scaled, centred ‘IBS’ method, described by Endelman and Jannink [80], was used to estimate additive genetic variance and generate the Kinship matrix of relationships among genotypes using *TASSEL*.

The statistical model used for the MLM is as follows:

Y = Xβ + Zu + e

Where Y is the vector of observations, β an unknown vector containing fixed effects including genetic marker and population structure (Q); u is an unknown vector of random additive genetic effects from multiple background QTL for the individuals; ***X*** and ***Z*** are the known design matrices; and ***e*** is the unobserved vector of random residuals.

Each marker allele is fit as a separate class with heterozygotes fit as an additional class so that the resulting marker effect is not broken down into additive and dominance effects. For the most robust MLM model used, the minor allele frequency (MAF) was set to > 0.05 (the SNPS were ‘filtered’). No missing values were included since SNPs with more than 14% missing values were removed and imputation was performed to replace any other missing values using the Euclidean distance measure in the k-nearest-neighbour algorithm [81] generated for the 10 nearest neighbours [77].

Phenotypic data for 75 accessions were imputed based on their genetic profiles.

#### Testing the robustness of the Mixed Linear Model (MLM) output

Model comparisons were made to test for putative false associations. These involved comparisons of the results of running *TASSEL* using the General Linear Model (GLM) with the Structure (Q) matrix and Principal Components Analysis (PCA) matrix, respectively, as opposed to the MLM with Structure and Kinship (Q+K). Price et al. [82] described how PCA corrects for stratification in GWAS as an alternative to the Q matrix.

GLM and MLM were run with filtered SNPs (612) and also with unfiltered SNPs with missing values ≤ 14% (737 SNPs). MLM was also run with unfiltered SNPs with missing data > 14%, but less than 20% (836 SNPs) for comparison.

#### Tests of significance of association

The test of significance in the *TASSEL* 5 software routine derived from the *F*-distribution assumes that the traits analysed have normally distributed residual error. The stringent Bonferroni correction test to cater for multiple testing [77] was applied to the derived *P* values to test the significance of the associations between traits and markers, based on the association analysis using the MLM routine in *TASSEL* with the Q matrix for *K*=7, Run 7. Manhattan plots, generated in R [83], were also used to check for evidence of *P* value inflation using *TASSEL* and to identify significant MTAs.

Simple M in R [83] was used to determine the number of independent markers that should be used in testing to avoid the penalty of the stringent Bonferroni correction test, as prescribed by Gao et al. [84]. There were 664 SNP markers that were independent and this number of markers was used to infer the Bonferroni threshold of 7.53 × 10^-5^ in the analyses with unfiltered SNPs.

To deduce positive associations, when more than one marker in the same linkage group was related to the same trait, only those separated by more than 2 cM in the reference map [29] were considered. This was reinforced by linkage disequilibrium (LD) decay patterns observed by Stack et al. [49] and in this study.

LD estimates are reported as squared correlations of allele frequencies (*r^2^*) and as D prime (*D^2^*). The correlation between alleles at two loci, *r^2^*, and the standardized disequilibrium coefficient for determining whether recombination or homoplasy had occurred between a pair of alleles, *D^2^*, were derived in this study. Fisher’s exact test was calculated to compare alleles at any two loci. The LD pattern was tested with 100,000 permutations in *TASSEL* to obtain *P* values for the tests. LD heatmap was used to scan for high linkage disequilibrium within chromosomes based on *r^2^* values. High LD was characterized by many red squares in the heatmap generated. Many red blocks together would correspond to haplotype blocks [85, 86].

LD decay was plotted in R [83] to show the points representing the distance, in Mb, on each linkage group/chromosome, at which the mean value of *r^2^* decreased to half of the maximum value. LD decay plots were generated and annotated for the chromosomes where significant marker trait associations were observed.

Quantile-quantile plots were generated in R [83] to search for evidence of bias in the GWAS, such as due to genotyping artifacts, and to discern the extent to which the observed distribution of the test statistic followed the expected (null) distribution.

The proportion of phenotypic variance explained by a marker was determined by the square of the partial correlation coefficient (R^2^%).

### Genomic prediction

The predictive breeding value (GEBV) (along with the predictive error variance (PEV)) for each phenotypic trait was derived using ridge regression in *TASSEL* 5.2.50 software, which performed multiple correlation tests to compute correspondence between genotype and phenotype and accounted for collinearity to avoid bias. The genotypic data (737 SNPs in alpha format) were loaded and kinship analysis performed to generate an Identity by Descent (IBD) file. The 421 phenotypes and *K* matrix were selected and the genomic selection routine was run with 5-fold cross validation with 100 iterations. The Best Linear Unbiased Prediction (BLUP) model (linear mixed model to estimate random effects) (*TASSEL* 4 software) was used to derive BLUPs for each SNP marker.

### Detecting candidate genes located within marker-trait association zones

The identification of candidate genes was done using the latest cocoa genome sequence (*T. cacao* Criollo genome version 2) [29] available on the cocoa genome hub (https://cocoa-genome-hub.southgreen.fr/jbrowse). A genetic linkage map, showing the position of putative candidate genes that co-localised with SNP markers associated with traits of interest (MTAs), was then constructed using *SpiderMap* (Rami, 2007 unpublished, Spidermap v1.7.1, free software, CIRAD).

## Results

### Phenotypic data analysis

The phenotypic data for the fully characterized accessions are presented in S2 Table. The large phenotypic variation expressed in this panel of cacao genotypes, based on the coefficients of variation (Table 2), indicated a diverse genetic background, which was suitable for GWAS. Correlation analysis revealed the interdependence of traits on one another. Of the 91 Pearson correlation coefficients, *r,* calculated for the quantitative traits, only 20 were not significant (*P* ≥ 0.05). It is noteworthy that the yield-related traits, seed number, seed/cotyledon mass and seed dimensions, were highly correlated (*P* ≤ 0.001) (Table 3a). As observed previously at the ICGT by Bekele et al. [37], seed number was negatively correlated (*r* = -0.205*** in this study) with individual dried cotyledon mass. Seed/cotyledon length and width were positively correlated with individual dried seed/cotyledon mass (*r* = 0.688*** and 0.751***, respectively). This suggests that the former traits can be used as indicators of the latter.

**Table 2.**
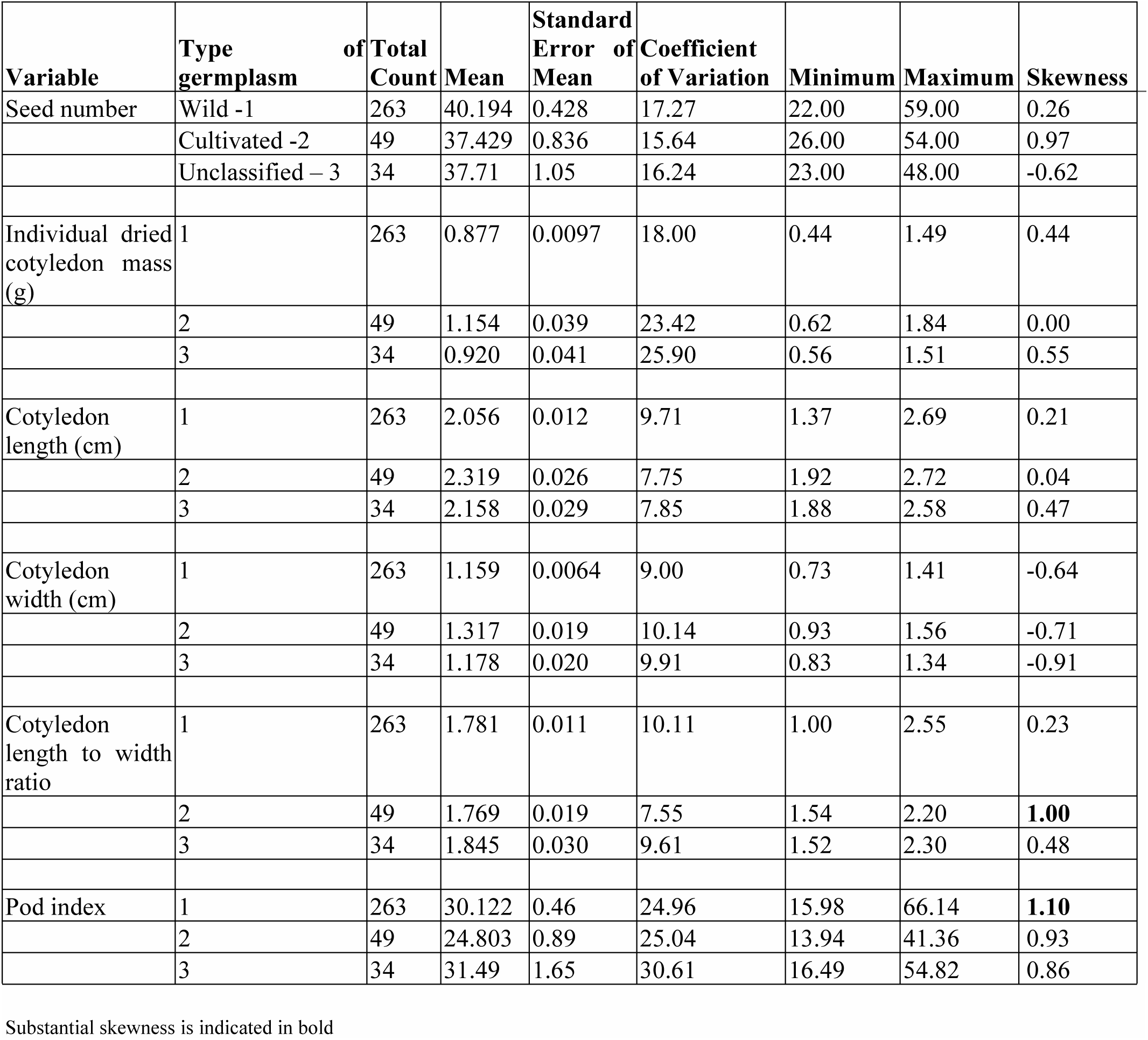
Descriptive statistics for quantitative fruit and seed traits in wild, cultivated and unclassified cacao germplasm studied.

| Variable | Type of germplasm | Total Count | Mean | Standard Error of Mean | Coefficient of Variation | Minimum | Maximum | Skewness |
| --- | --- | --- | --- | --- | --- | --- | --- | --- |
| Seed number | Wild -1 | 263 | 40.194 | 0.428 | 17.27 | 22.00 | 59.00 | 0.26 |
|  | Cultivated -2 | 49 | 37.429 | 0.836 | 15.64 | 26.00 | 54.00 | 0.97 |
|  | Unclassified – 3 | 34 | 37.71 | 1.05 | 16.24 | 23.00 | 48.00 | -0.62 |
| Individual dried cotyledon mass (g) | 1 | 263 | 0.877 | 0.0097 | 18.00 | 0.44 | 1.49 | 0.44 |
|  | 2 | 49 | 1.154 | 0.039 | 23.42 | 0.62 | 1.84 | 0.00 |
|  | 3 | 34 | 0.920 | 0.041 | 25.90 | 0.56 | 1.51 | 0.55 |
| Cotyledon length (cm) | 1 | 263 | 2.056 | 0.012 | 9.71 | 1.37 | 2.69 | 0.21 |
|  | 2 | 49 | 2.319 | 0.026 | 7.75 | 1.92 | 2.72 | 0.04 |
|  | 3 | 34 | 2.158 | 0.029 | 7.85 | 1.88 | 2.58 | 0.47 |
| Cotyledon width (cm) | 1 | 263 | 1.159 | 0.0064 | 9.00 | 0.73 | 1.41 | -0.64 |
|  | 2 | 49 | 1.317 | 0.019 | 10.14 | 0.93 | 1.56 | -0.71 |
|  | 3 | 34 | 1.178 | 0.020 | 9.91 | 0.83 | 1.34 | -0.91 |
| Cotyledon length to width ratio | 1 | 263 | 1.781 | 0.011 | 10.11 | 1.00 | 2.55 | 0.23 |
|  | 2 | 49 | 1.769 | 0.019 | 7.55 | 1.54 | 2.20 | **1.00** |
|  | 3 | 34 | 1.845 | 0.030 | 9.61 | 1.52 | 2.30 | 0.48 |
| Pod index | 1 | 263 | 30.122 | 0.46 | 24.96 | 15.98 | 66.14 | **1.10** |
|  | 2 | 49 | 24.803 | 0.89 | 25.04 | 13.94 | 41.36 | 0.93 |
|  | 3 | 34 | 31.49 | 1.65 | 30.61 | 16.49 | 54.82 | 0.86 |

Substantial skewness is indicated in bold

**Table 3A.** Pearson correlation coefficients for quantitative phenotypic traits.

|  | Flower sepal length | Ligule width | Ovule number | Style length | Fruit length | Fruit width | Fruit length to width ratio | Total wet seed mass |
| --- | --- | --- | --- | --- | --- | --- | --- | --- |
| Ligule width | 0.429*** |  |  |  |  |  |  |  |
| Ovule number | -0.092 | 0.142* |  |  |  |  |  |  |
| Style length | 0.363*** | 0.281*** | 0.098 |  |  |  |  |  |
| Fruit length | 0.291*** | 0.237*** | 0.050 | 0.168* |  |  |  |  |
| Fruit width | 0.338*** | 0.272*** | 0.131* | 0.170* | 0.498*** |  |  |  |
| Fruit length: width | 0.003 | 0.012 | -0.062 | 0.031 | **0.605\*\*\*** | -0.383*** |  |  |
| Total wet seed mass | 0.324*** | 0.310*** | 0.126* | 0.004 | **0.518\*\*\*** | **0.701\*\*\*** | -0.098 |  |
| Seed number | 0.117* | 0.277*** | **0.568\*\*\*** | 0.066 | 0.303*** | 0.326*** | 0.032 | 0.474*** |
| Cotyledon mass (individual) | 0.189*** | 0.124* | -0.233*** | -0.088 | 0.365*** | **0.510\*\*\*** | -0.096 | **0.684\*\*\*** |
| Cotyledon length | 0.293*** | 0.315*** | 0.073 | -0.025 | 0.361*** | **0.680\*\*\*** | -0.239*** | **0.762\*\*\*** |
| Cotyledon width | 0.122* | 0.122* | -0.178* | -0.167* | 0.009 | 0.311*** | -0.292*** | **0.523\*\*\*** |
| Cotyledon length: width | 0.188*** | 0.193*** | 0.263*** | 0.152* | 0.386*** | 0.375*** | 0.085 | 0.229*** |
| Pod index | -0.158* | -0.244*** | -0.177* | 0.045 | -0.496*** | **-0.621\*\*\*** | 0.052 | **-0.862\*\*\*** |

|  | Seed number | Cotyledon mass | Cotyledon length | Cotyledon width | Cotyledon length to width ratio |
| --- | --- | --- | --- | --- | --- |
| Ligule width |  |  |  |  |  |
| Ovule number |  |  |  |  |  |
| Style length |  |  |  |  |  |
| Fruit length |  |  |  |  |  |
| Fruit width |  |  |  |  |  |
| Fruit length: width |  |  |  |  |  |
| Total wet seed mass |  |  |  |  |  |
| Seed number |  |  |  |  |  |
| Cotyledon mass | -0.205*** |  |  |  |  |
| Cotyledon length | 0.235*** | 0.688*** |  |  |  |
| Cotyledon width | -0.165* | 0.751*** | 0.576*** |  |  |
| Cotyledon length to width ratio | 0.419*** | -0.095* | 0.408*** | -0.501*** |  |
| Pod index | -0.506*** | -0.684*** | -0.700*** | -0.523*** | -0.155* |

There was also a strong correlation (0.568***) between ovule number and seed number that justifies prediction of the latter using the former when fruits are unavailable. The correlations of fruit width with pod index and seed length, *r* = -0.621*** and 0.680***, respectively, are also noteworthy (Table 3A).

The Spearman correlations for anthocyanin intensity in the various plant organs are presented in Table 3B. All of these correlations were significant (*P* ≤ 0.05) except for the anthocyanin pigment concentration in the pedicel and anthocyanin pigmentation of the cotyledon (*r* = -0.097). It was notable that the correlations involving mature fruit surface (ridges) anthocyanin intensity and that in the ligule and filament of the flower and in the seed/cotyledon were all significantly (*P* ≤ 0.05) negative (*r* = - 0.150, -0.257, -0.147, respectively).

**Table 3B.**
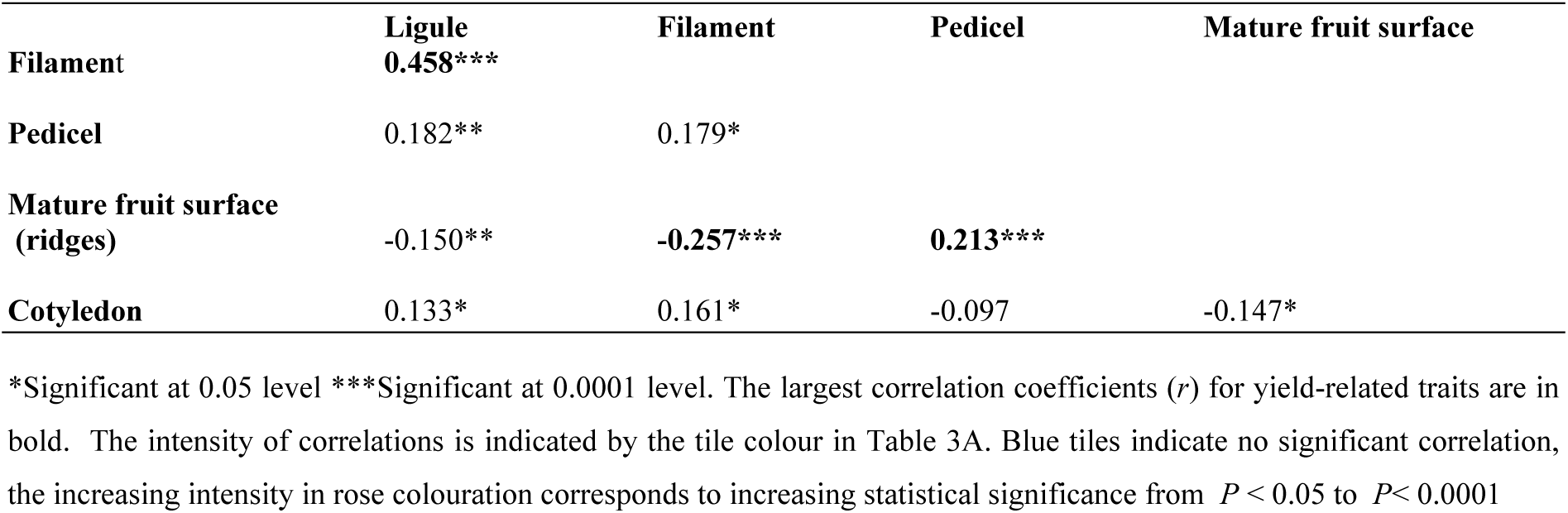
Spearman correlation coefficients for anthocyanin intensity in various plant organs.

Tests of normality indicated significant deviation from normality for fruit length, width, fruit length to width ratio, total fresh seed mass, seed number, individual seed (cotyledon) mass and width, seed length to width ratio, pod index, ovule number, sepal length and width, and style length. Results of tests of normality performed on the natural log transformed values of fruit and seed quantitative traits indicated that natural log transformation corrected for the deviation from normality of these traits (S3 Table). The phenotypic data were thus natural log transformed to conduct GWAS. Untransformed data were also subjected to GWAS for comparison.

### Population structure

Fifteen replicate runs of models using genotypic clusters (*K*) from 2 to 15 confirmed that *K* = 7 had the highest log-likelihood probability (log Pr (X | *K*) versus *K*) (Fig 1).

The population structure analysis revealed that 74% of the accessions could be stratified into seven sub-populations, while 26% could be regarded as admixtures. The constitution of the seven genetic clusters is presented in Table 4 and S4 Table. The IMC and AMAZ groups were clustered together as were also the SCA, MO and LCTEEN groups (Table 4; S4 Table). Several Upper Amazon accessions were genotyped as Trinitario, and are putatively mislabelled. The genetic diversity of the accessions studied is represented in the neighbour-joining tree in Fig 2, which depicts the seven clusters differentiated. (One SPEC accession was grouped alone and this was not considered a cluster).

**Table 4.** The fixation index as a measure of population differentiation due to genetic structure for each cluster identified by *STRUCTURE* analysis.

| Cluster | Main genetic clusters represented | Mean value of Fixation Index |
| --- | --- | --- |
| 1 | PARINARI (PA) | 0.5106 |
| 2 | LCT_EEN, SCAVINA (SCA), MORONA (MO) | 0.7204 |
| 3 | NANAY (NA), POUND | 0.8665 |
| 4 | SPEC, ICS | 0.8653 |
| 5 | CRIOLLO, ICS | 0.9951 |
| 6 | GUIANESE (GU) | 0.8378 |
| 7 | AMAZ, IQUITOS (IMC) | 0.5096 |

Mean value of Fixation Index (membership coefficient) $\geq 0.7$ is indicative of strong population differentiation
Individuals with coefficients of less than 0.7 were classified as admixed

### Linkage disequilibrium (LD)

*D^2^* greater than 0.6, where *D^2^* of 1 represents the highest amount of disequilibrium possible, was indicative of recombination or homoplasy between pairs of alleles (Table 5). With regard to the squared correlations of allele frequencies, *r^2^* values, none were observed to be greater than 0.2 and the mean *r^2^* across the 10 chromosomes was 0.1 (Table 5). Chromosomes 1 and 3 had mean *r^2^* > 0.14. These findings seem to support that of Stack et al. [49], who found that the wild genotypes studied (such as those from the Purus, Contamana, Curaray, Iquitos, Nanay, Marañón and Guiana genetic groups) exhibited very low overall LD, measured by *r^2^*, which rapidly decayed within 1–2 Mbps to a value of around 0.1. In this study, an average decay of *r^2^* to 50% over chromosomes 1, 4, 5, 7 and 9 occurred at a relatively short distance of 5.21 Mb (9 cM) on average (S5 Table). Motilal et al. [87] reported decay to half over a distance of 9.3 cM on chromosomes 1 to 9 for the germplasm they studied. LD decay plots, based on *r^2^*, for this study are presented in Fig 3A. These findings suggest that MTAs could be “localized in the genome with relatively high precision using an association mapping approach” [49], particularly when wild germplasm is studied.

**Fig 3A.**
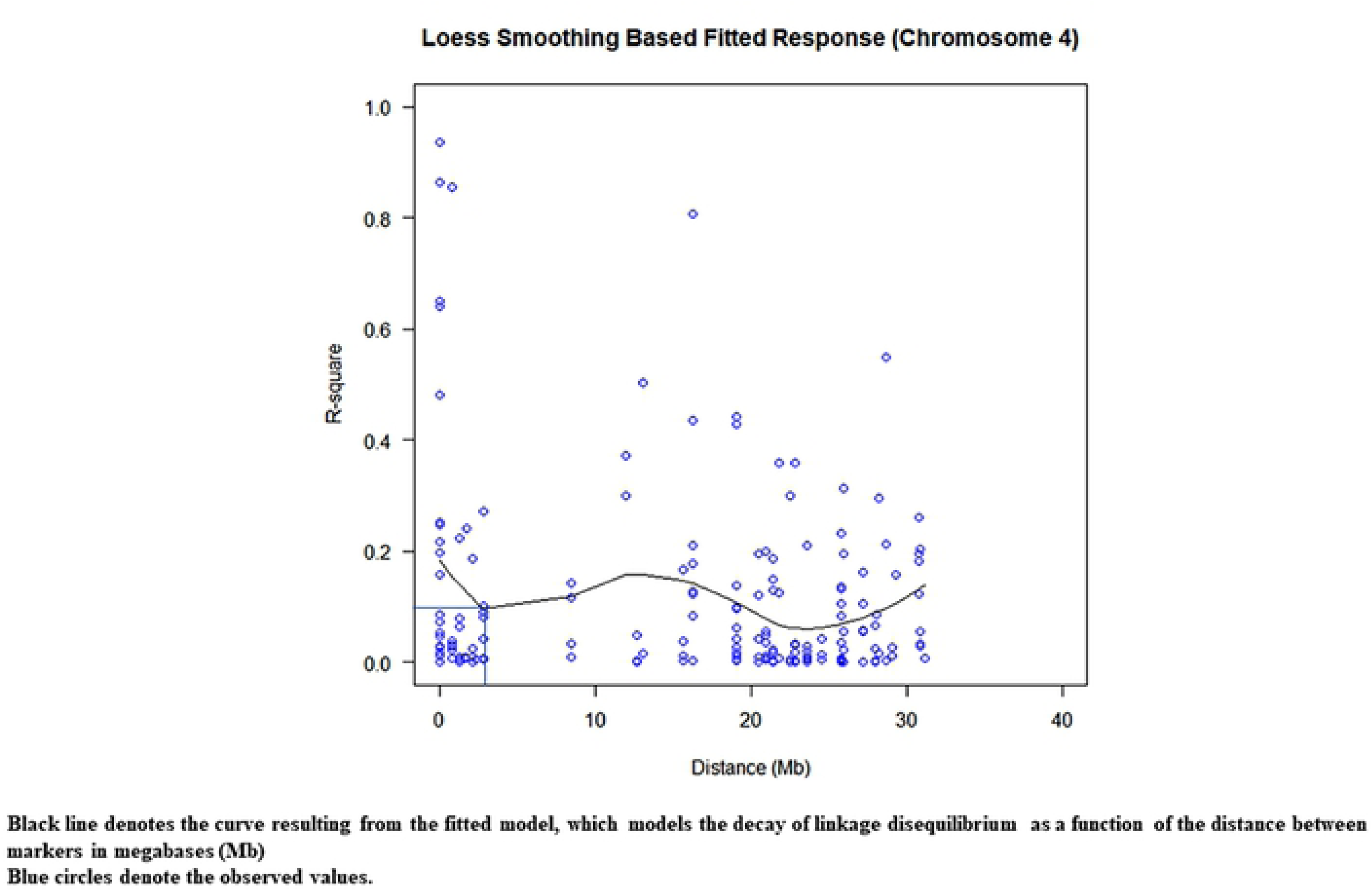
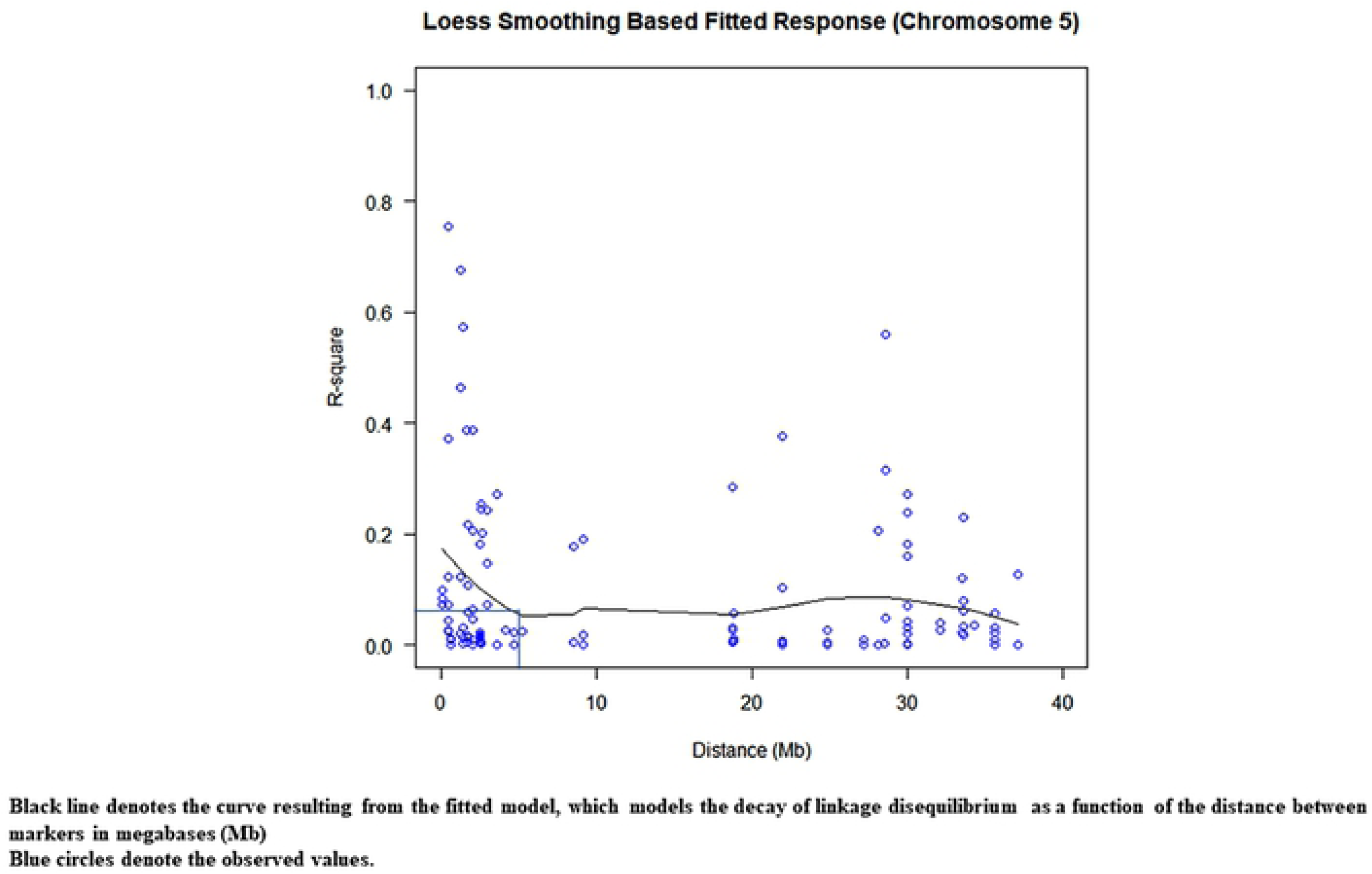
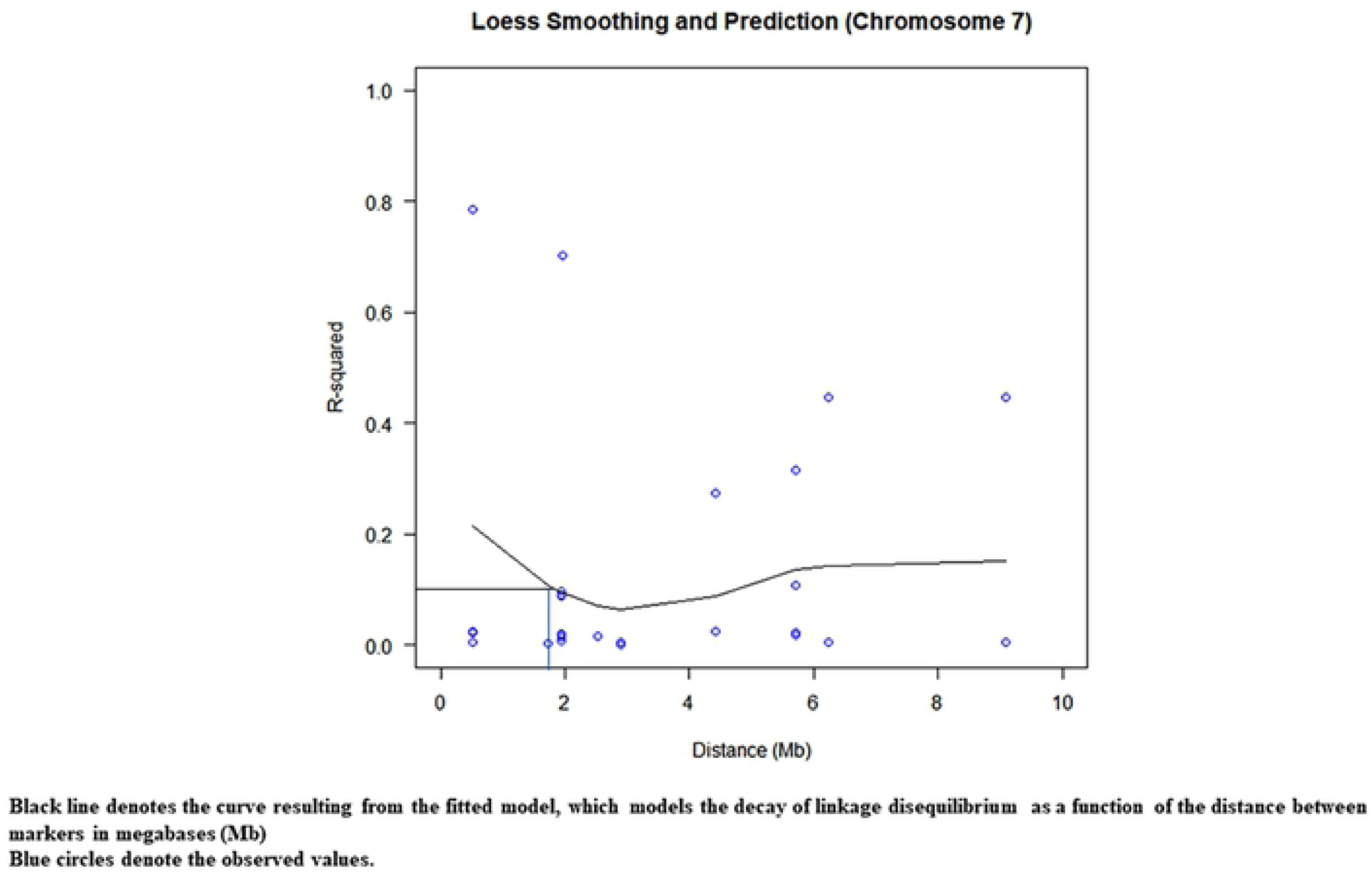
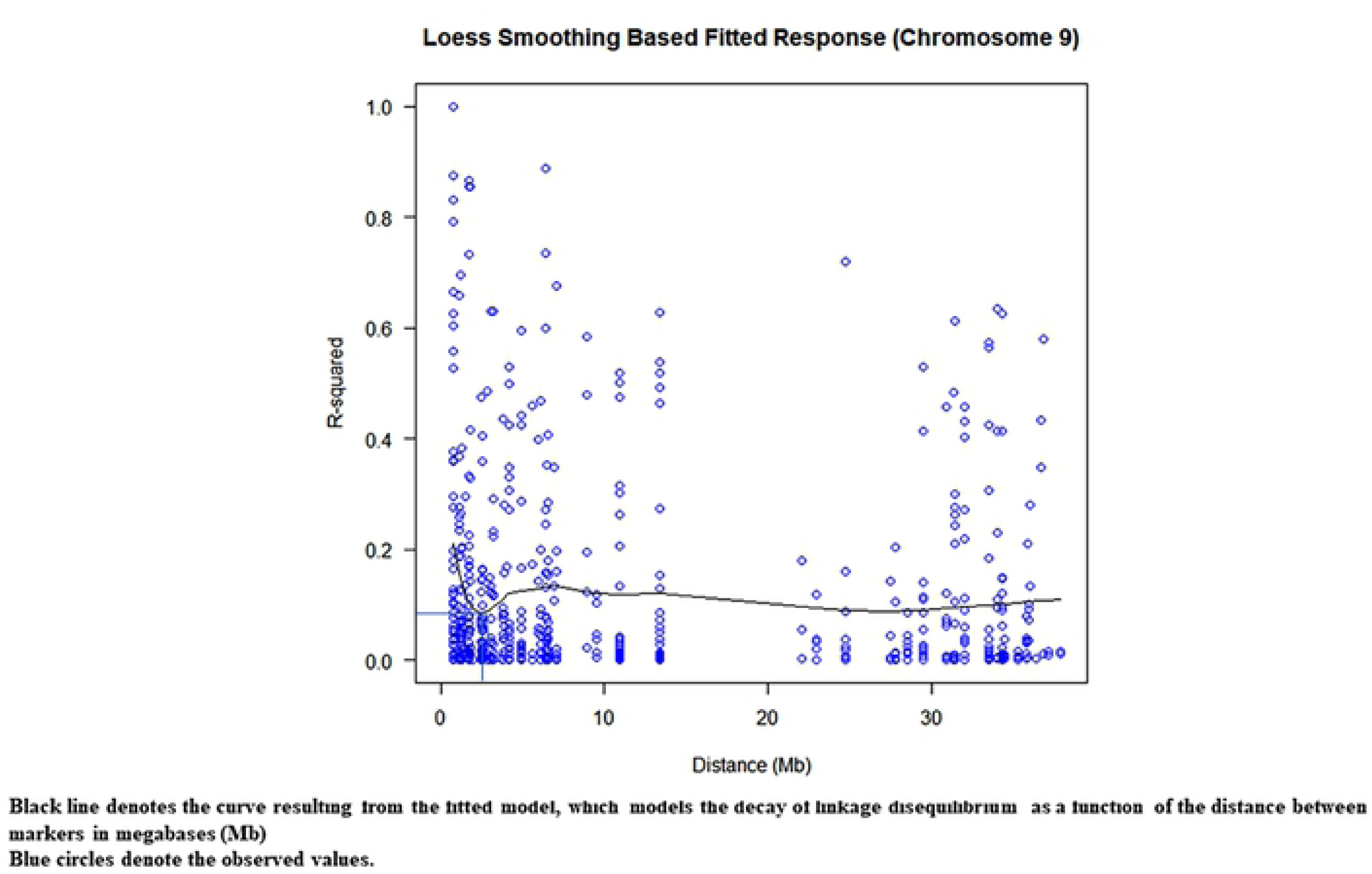
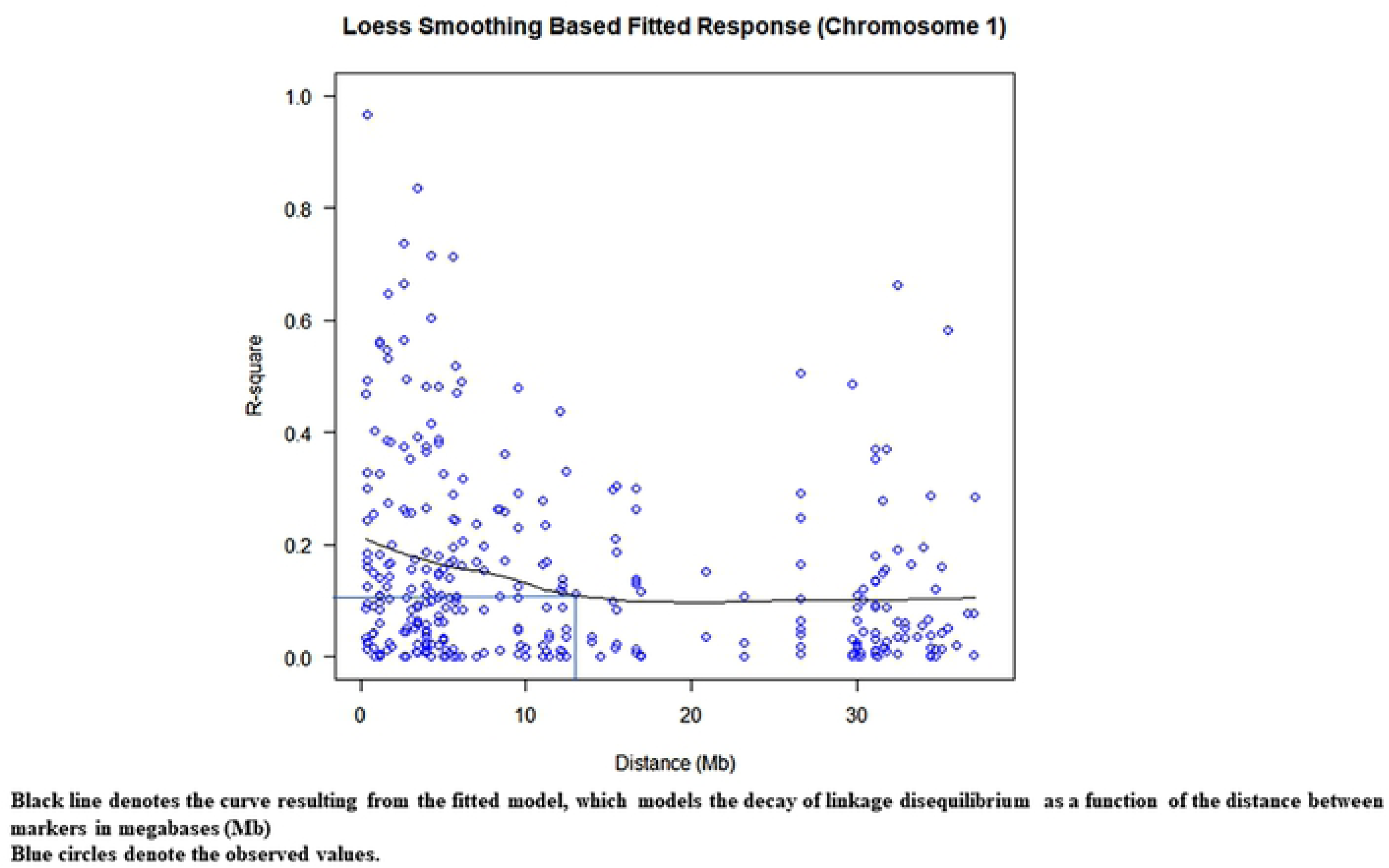
Plots modelling the decay in pairwise linkage disequilibrium coefficients (*r*^2^) as a function of the distance between markers in megabases (Mb). Plot of pairwise linkage disequilibrium coefficients (r2) on chromosome 1; Plot of pairwise linkage disequilibrium coefficients (r2) on chromosome 4; Plot of pairwise linkage disequilibrium coefficients (r2) on chromosome 5; Plot of pairwise linkage disequilibrium coefficients (r2) on chromosome 7; Plot of pairwise linkage disequilibrium coefficients (r2) on chromosome 9.

**Table 5.**
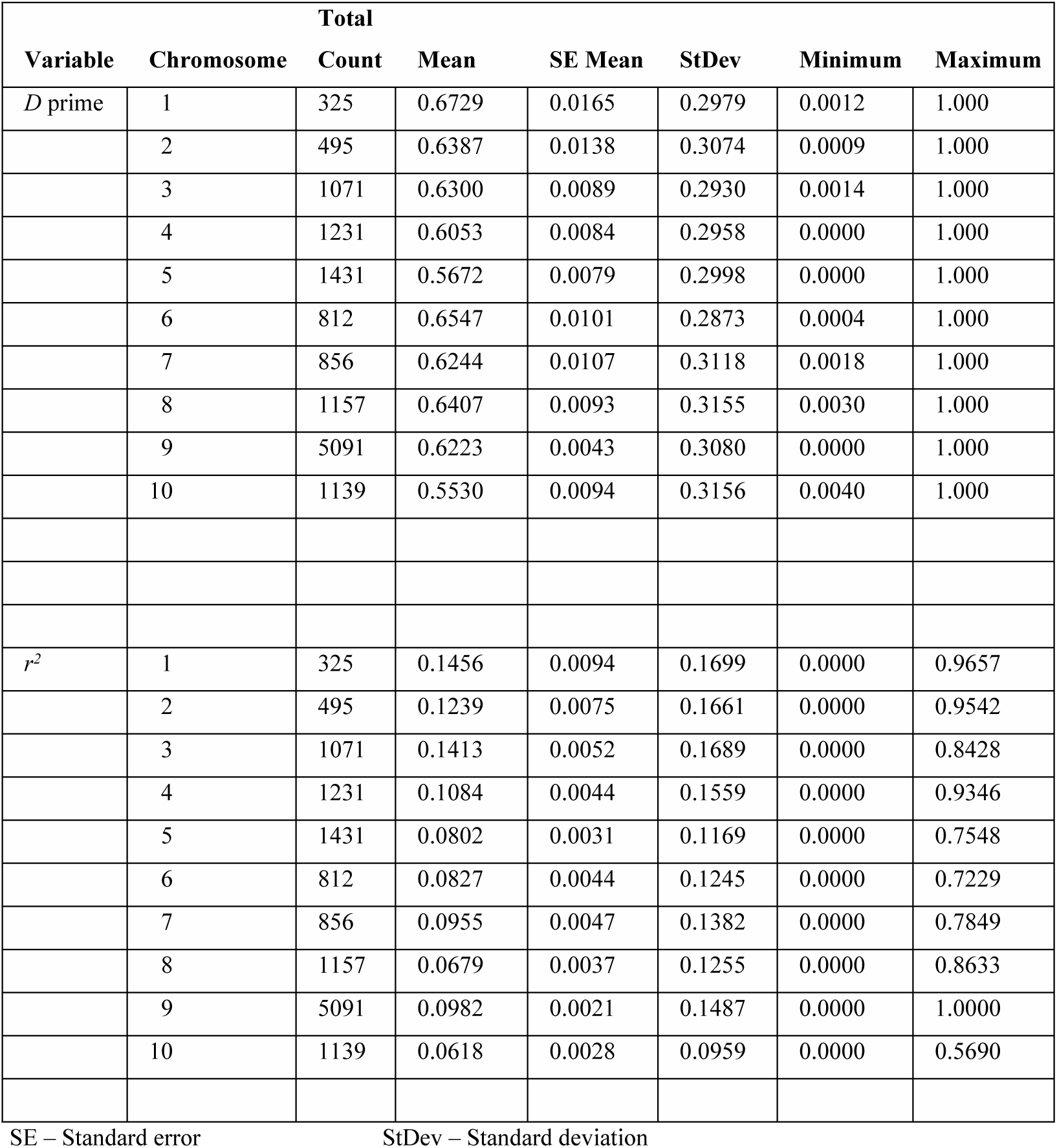
Results of linkage disequilibrium analysis.

It is noteworthy that for cultivated germplasm such as Trinitario (*e.g*., the ICS accession group), LD decay was observed by Stack et al. [49] to be very gradual with increasing marker distance, falling below 0.1 at approximately 30 Mbps. Marcano et al. [88] found LD to decay to half over 25-35 cM for the recently admixed populations (Meso-American Criollo and South American Forastero) they studied.

The linkage disequilibrium (LD) decay patterns, observed for the chromosomes in this study population and a LD heat map (Figs 3A and 3B, S5 Table), were used to inform the process of searching for putative candidate genes. JBrowse in version 2 of the Criollo B97-61/B2 genome (https://cocoa-genome-hub.southgreen.fr/) was used to locate candidate genes upstream or downstream of MTA zones. The search for putative candidate genes with relevant functional roles was conducted within defined intervals, based on linkage disequilibrium decay, spanning MTAs on each chromosome. For chromosomes 3, 5, 6 and 8, this was over a distance of 2.5 Mb upstream or downstream of the significant MTAs. It was over a distance of 5 Mb for chromosome 1, 1.6 Mb for chromosome 4, 0.87 Mb for chromosome 7 and 0.86 Mb for chromosome 9.

**Fig 3B.**
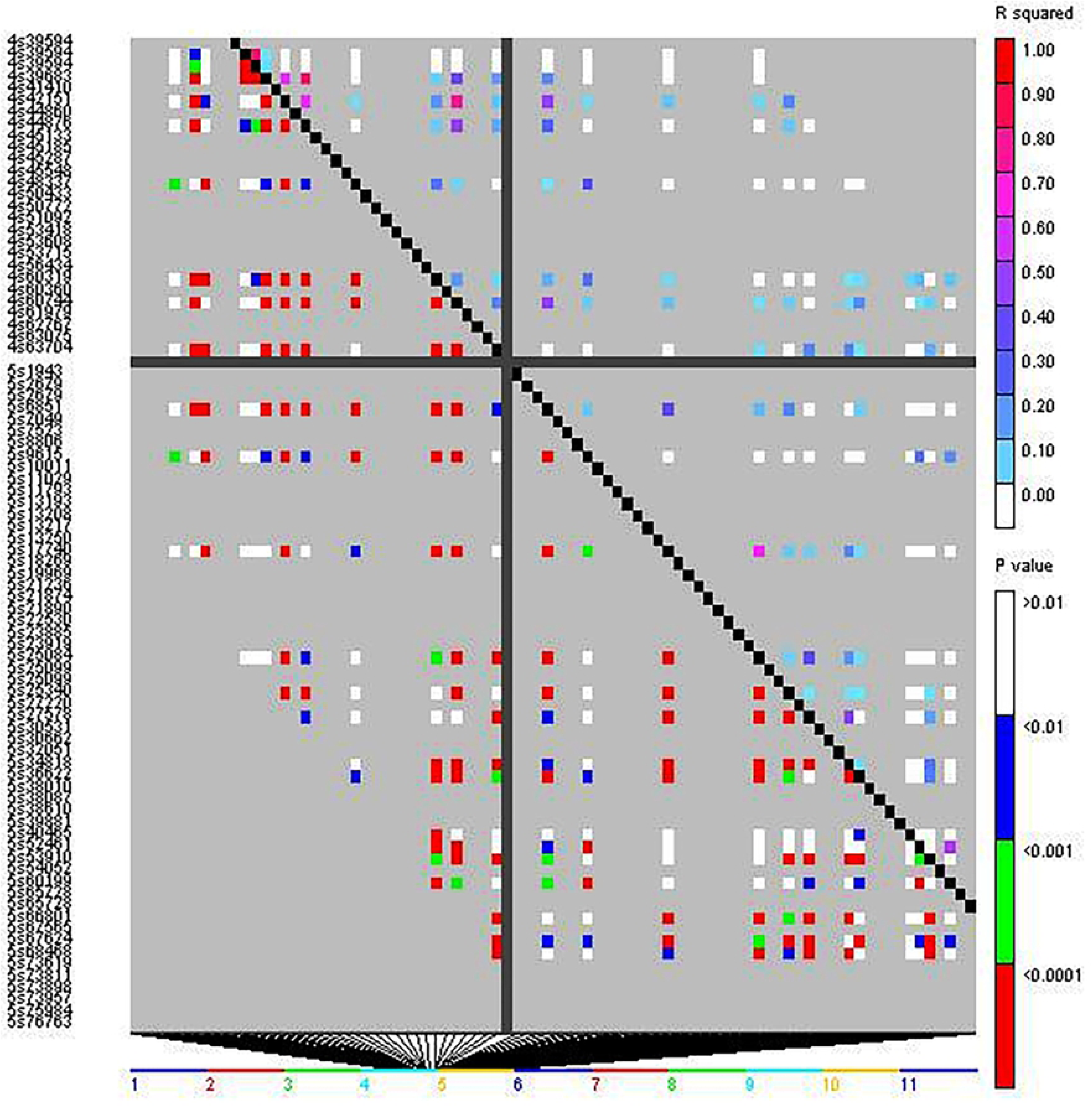
Heatmap of linkage disequilibrium (*r*^2^) across the chromosomes 4 and 5 based on data for 421 cacao accessions genotyped using 612 filtered SNPs. Legend: Markers were ordered on the x and y axes according to location along the chromosomes and each cell of the heat map represents a single marker pair. The upper triangle, above the black diagonal, is colour-coded based on the *r*^2^ value between SNPs while colours depicted in the lower triangle are based on *P*-values for the corresponding *r*^2^ values.

### Comparison of results generated for different models utilized in GWAS

When 836 SNPs (unfiltered and without correction for missing values >14% <20%) were used for the MLM routine with Bonferroni threshold of *P* ≤ 7.53 ×10^-5^ (0.05/664; 664 being the number of independent SNP markers employed)), 81 significant associations were found with MLM + PCA (accounting for population structure) while 71 significant associations were found with MLM + Q (matrix derived using *STRUCTURE*). In comparison, when the SNPS were filtered (with MAF ≥ 0.05) in *TASSEL* prior to executing MLM with Q+K, 612 SNPS were retained, and 17 significant associations were observed.

Performing *TASSEL* without correcting for kinship using the GLM model, GLM + Q with SNPS filtered (MAF ≥ 0.05), resulted in 5410 significant marker-trait associations at *P* ≤ 7.53 × 10^-5^. It was inferred that most of these MTAs were spurious. In addition, the Manhattan plots generated for the chromosomes showed no clearly distinct peaks. When the results were sorted on marker effects, *R^2^*, greater than 0.2, 95 positive associations were obtained using GLM with filtered SNPs. One of these involved fruit wall hardness and TcSNP1411 on chromosome 3 (position 25,585,284) and TcSNP1334 also on chromosome 3 (position 31,031,130). However, when the robust MLM was used for GWAS, no significant MTAs were found involving this trait, which is of interest due to its putative correlation with resistance to Cocoa Pod Borer attack [37].

### Putatively robust marker-trait associations

The close observed and expected distributions of the –log10(P) values in the MLM+K+Q model, suggested a reduction of potential spurious MTAs for MLM analysis based on 612 filtered SNPS. Consequently, significant associations obtained using this MLM model and Bonferroni threshold of *P* ≤ 8.17 × 10^-5^ (–log10 (P) = 4.088) were scrutinized for potentially authentic/stable marker-trait associations with putative functional value. There was a predominance of positive associations between highly heritable qualitative traits such as fruit shape, anthocyanin intensity on fruit surface [89], fruit apex form, and flower filament anthocyanin intensity and SNP markers. However, there were also some significant associations between SNPs and quantitative traits. Seventeen significant (*P* ≤ 8.17 × 10^-5^) MTAs were identified, including between seed number and log seed length and TcSNP 1335 on chromosome 7 (Fig 4) (at *P* ≤ 1.15 × 10^-14^ and *P* ≤ 6.75 × 10^-05^, respectively), and log seed number and TcSNP 785 on chromosome 1 (*P* ≤ 2.38 × 10^-05^) (Table 6). Manhattan plots and associated Quantile-Quantile plots are presented for yield-related traits in Fig 5A and 5B. The relatively low number of SNPs significantly associated with phenotypic traits, 2.4% of all SNPs used, was probably partly due to the stringent statistical thresholds applied in this study.

**Fig 4.**
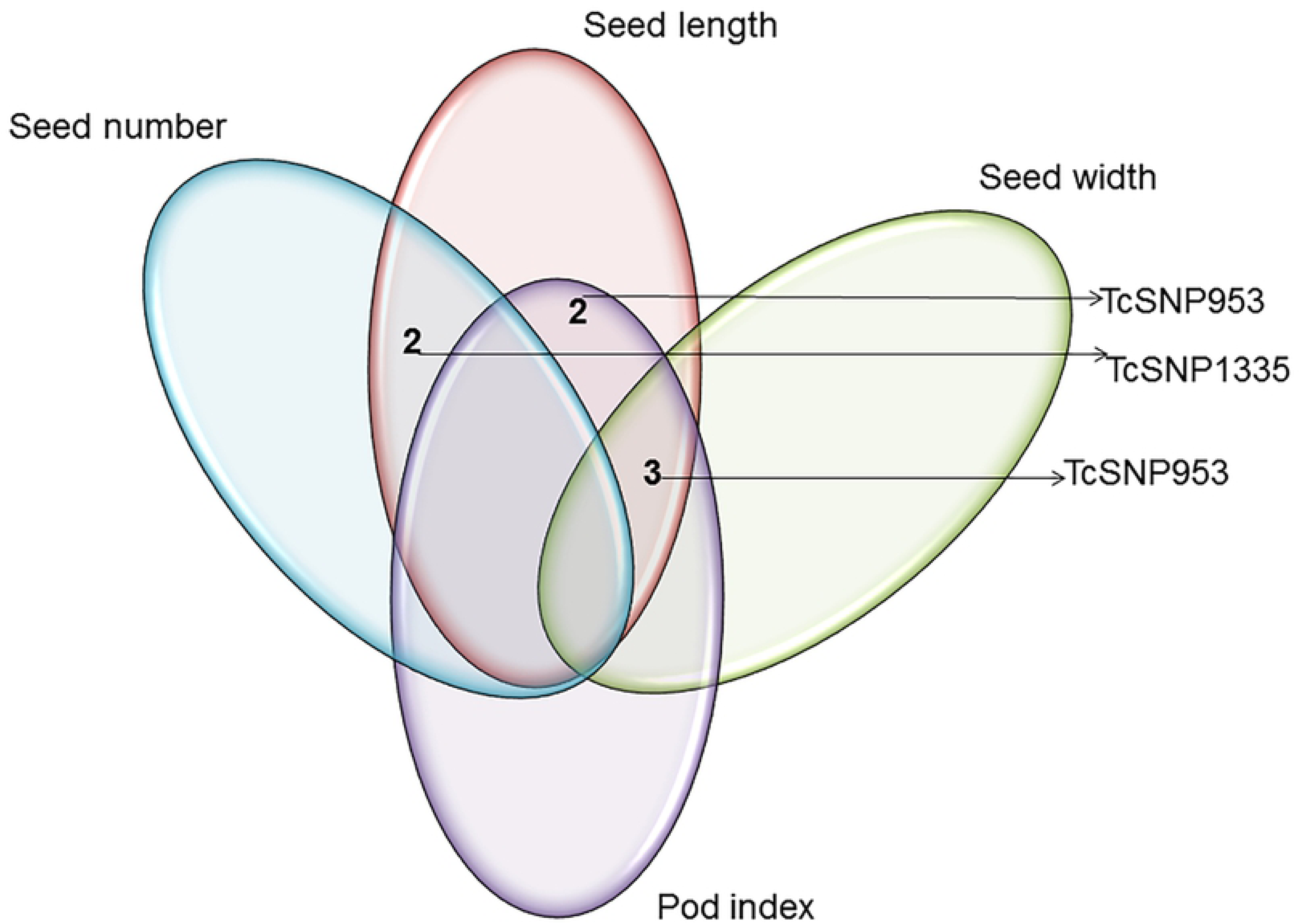
Venn diagram depicting relationships among seed traits of interest based on common associations with SNP markers.

**Fig 5A.**
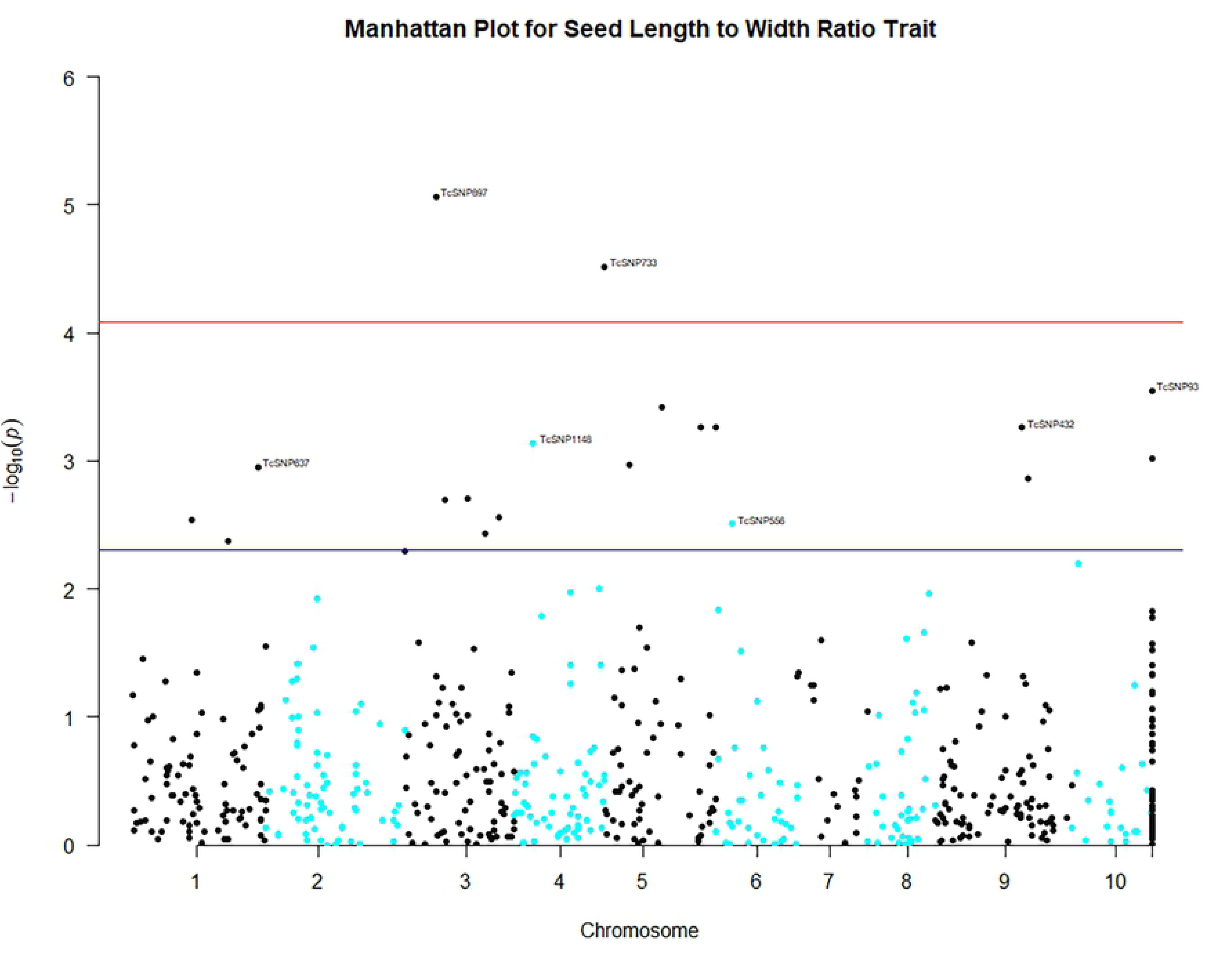
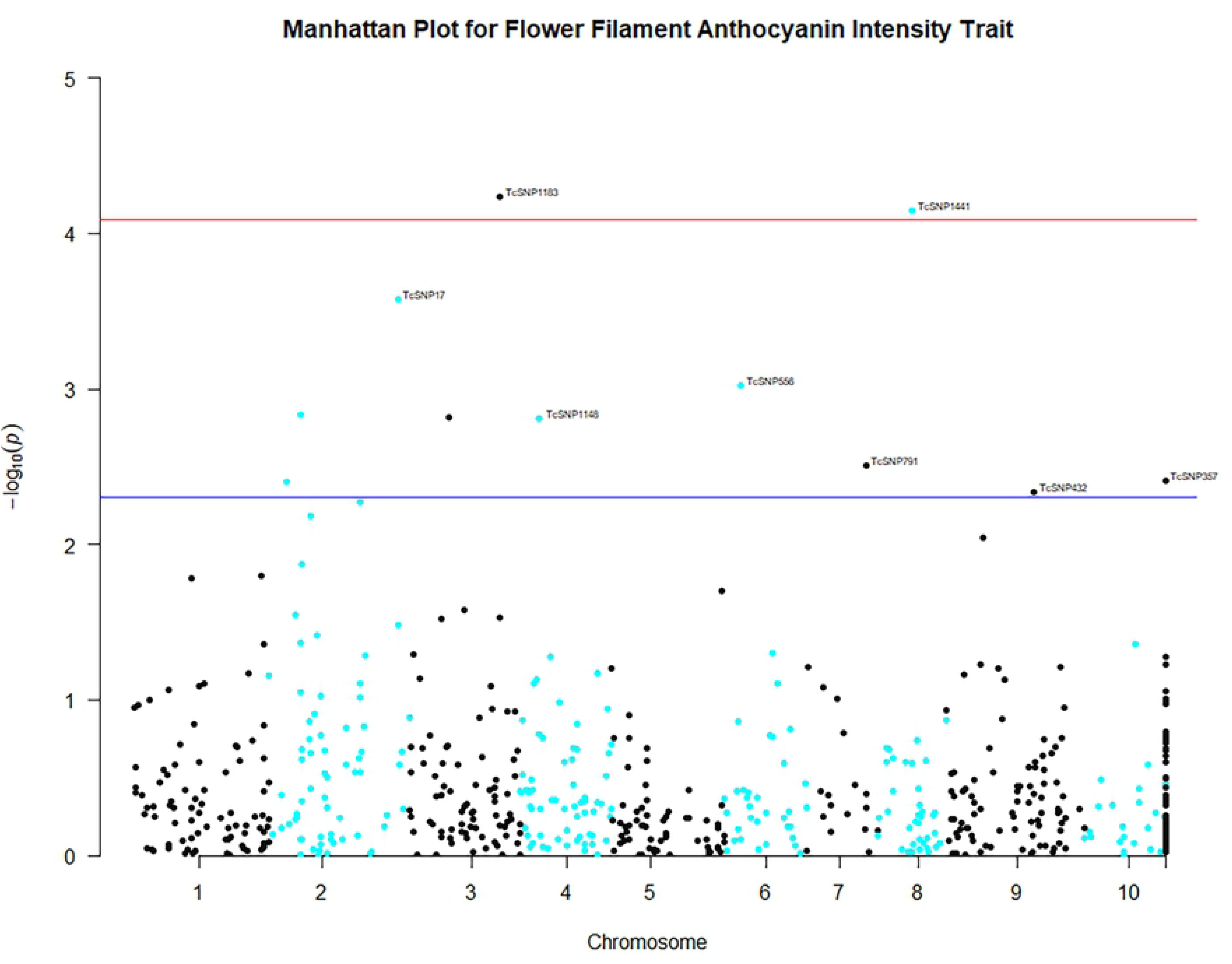
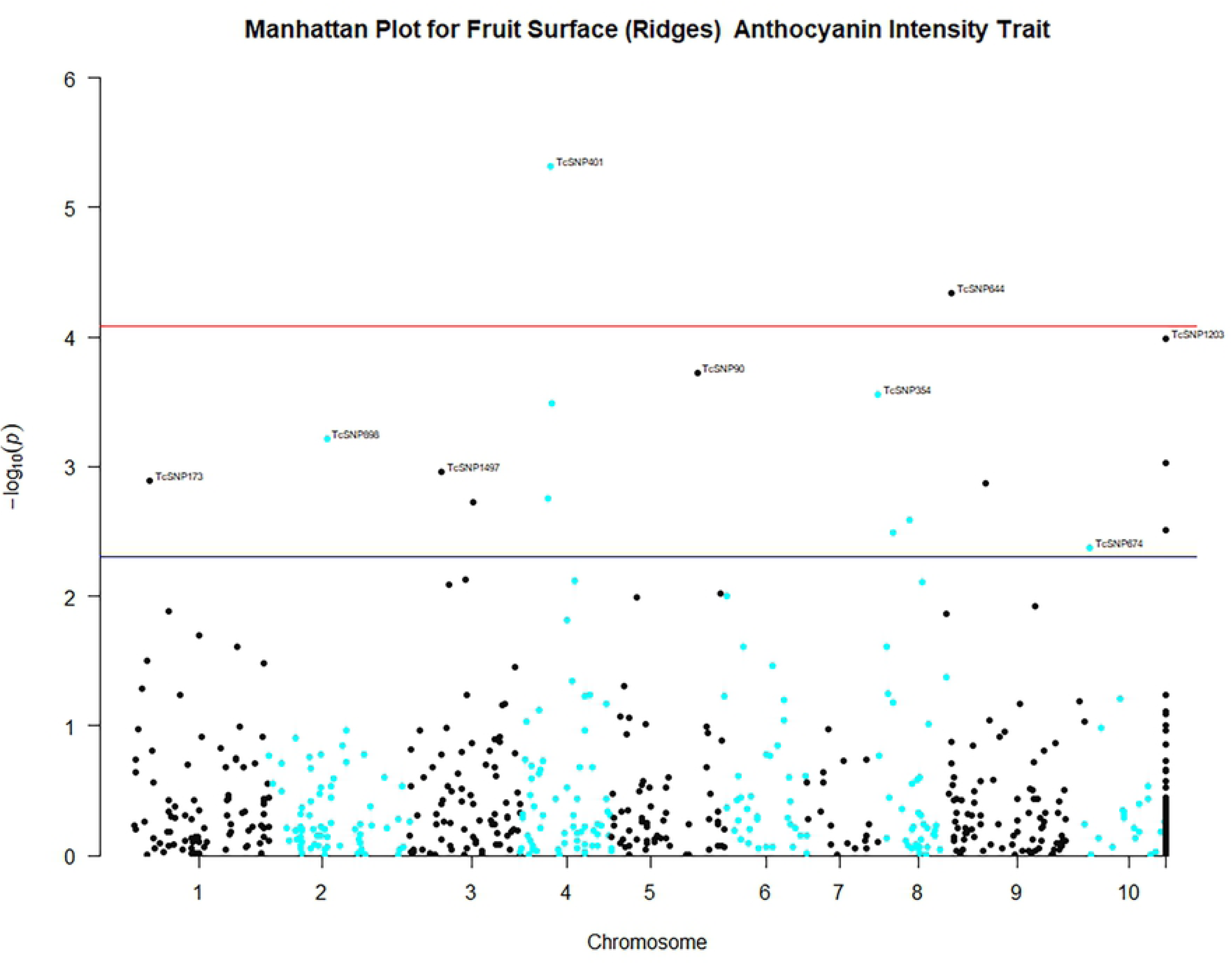
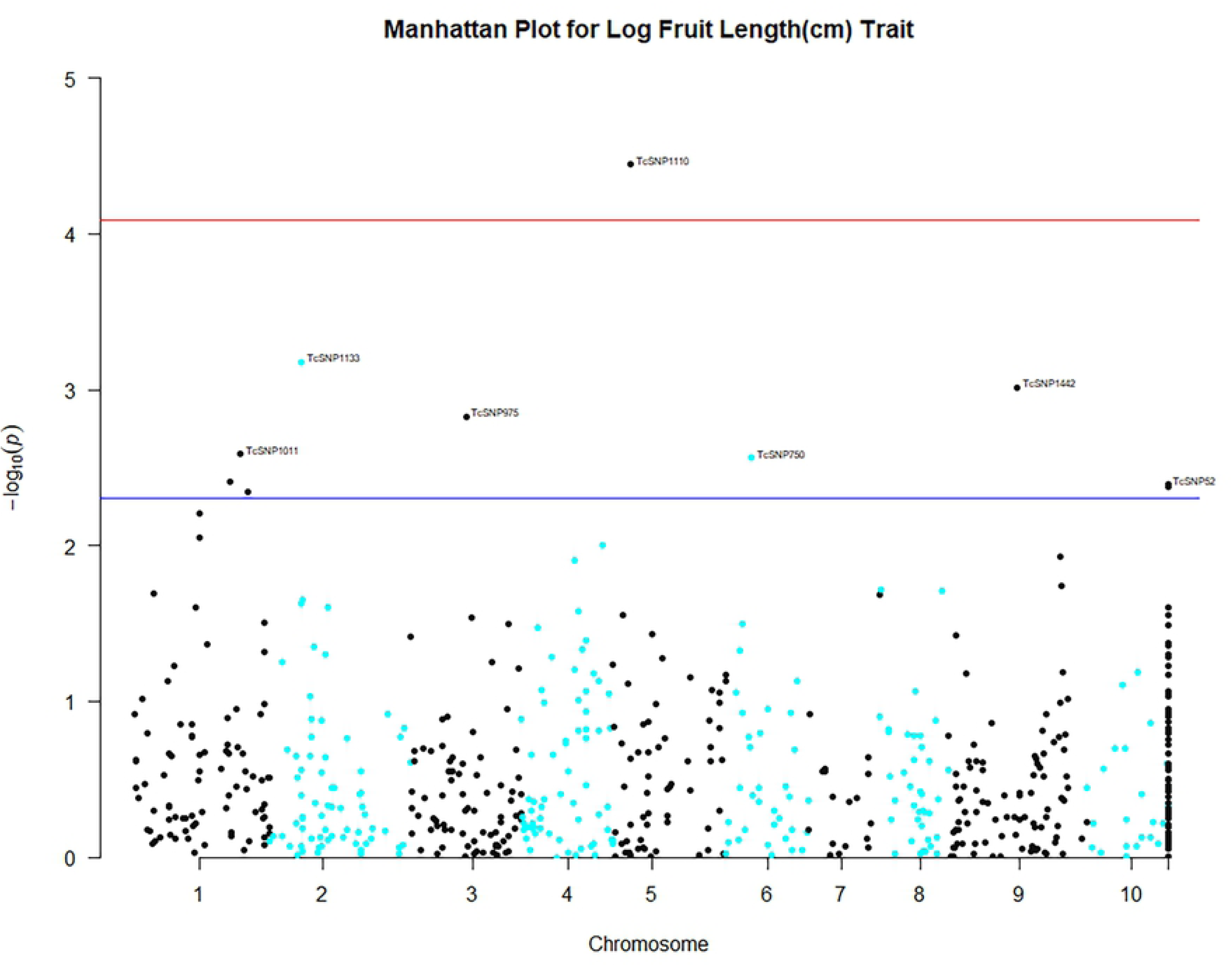
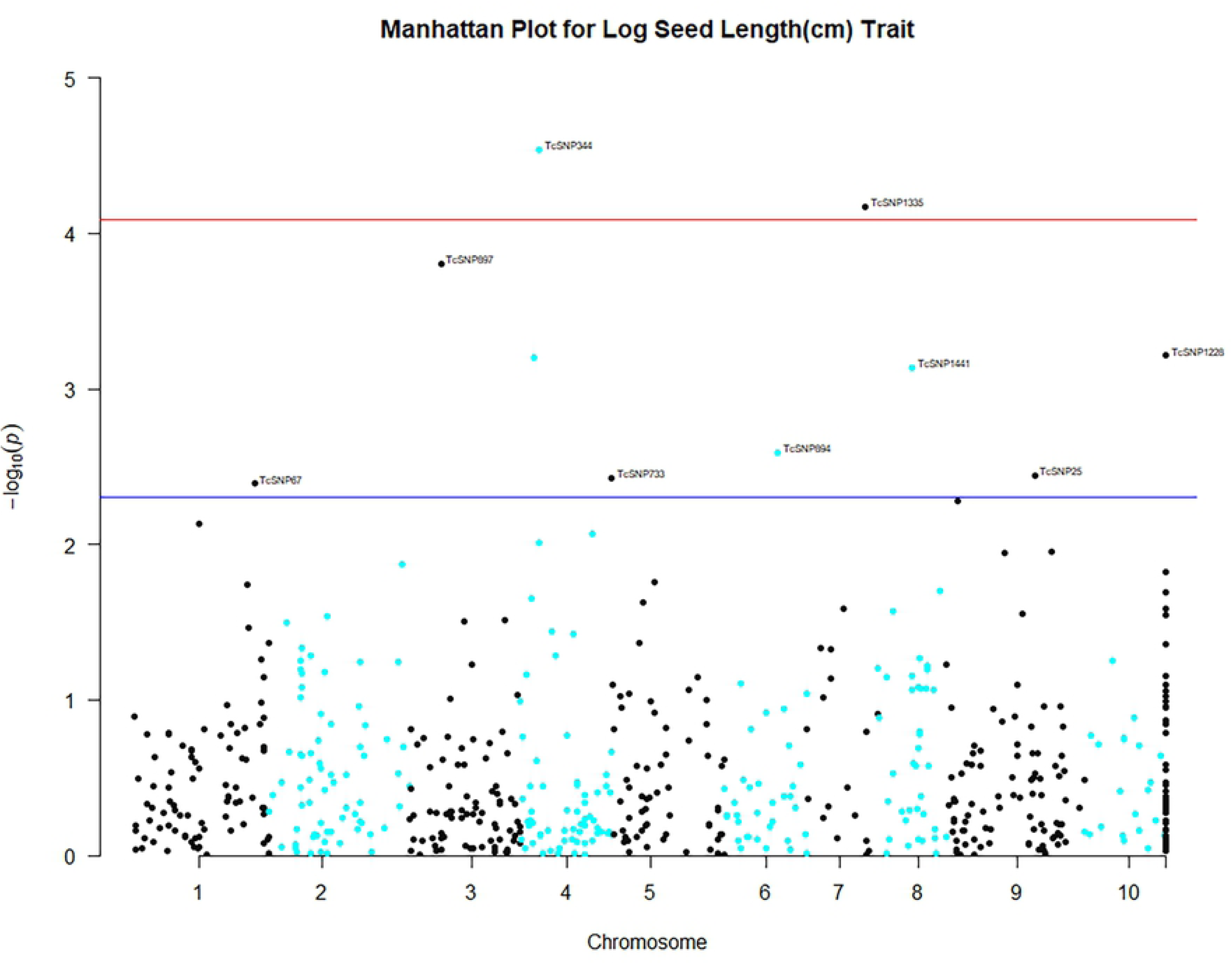
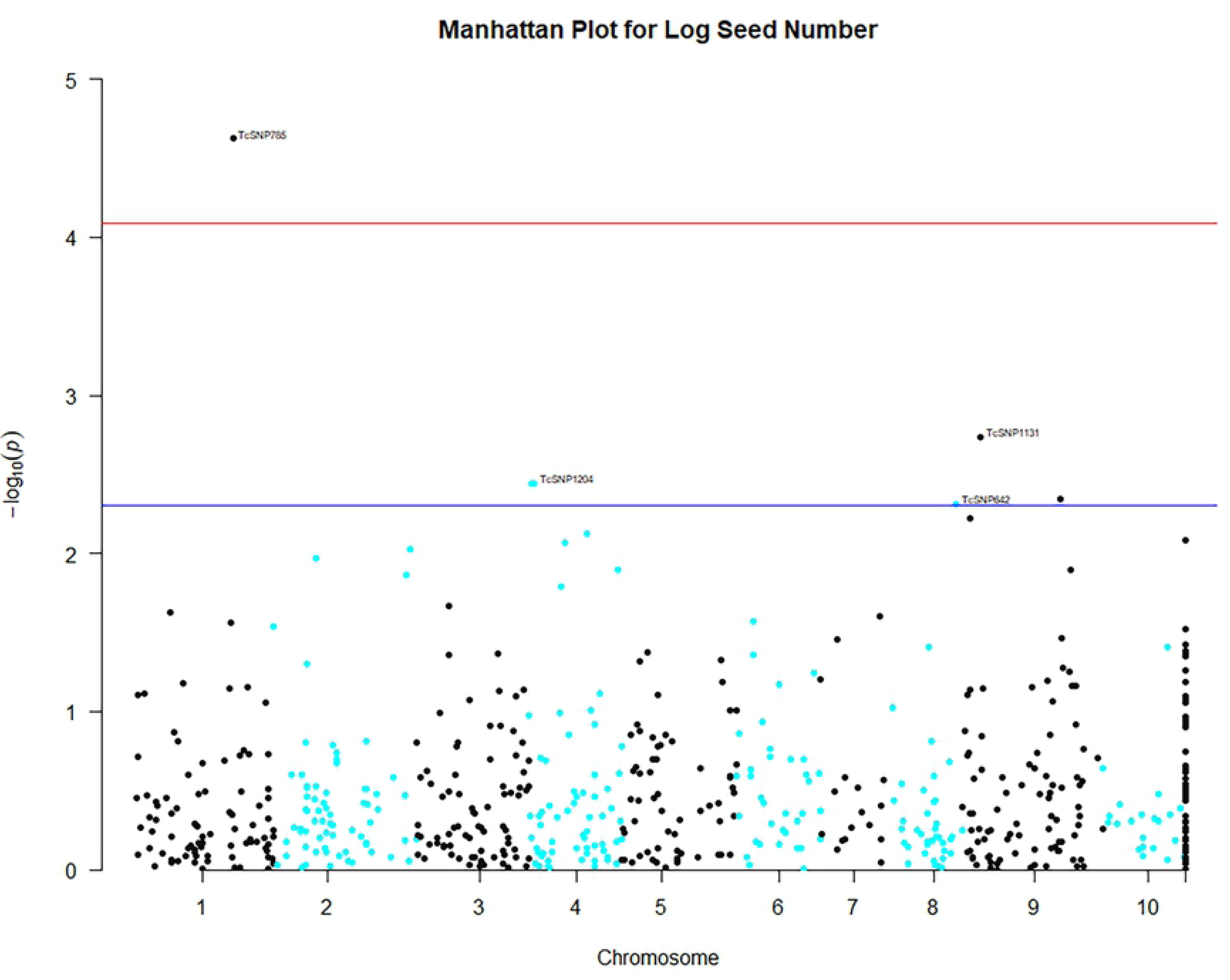
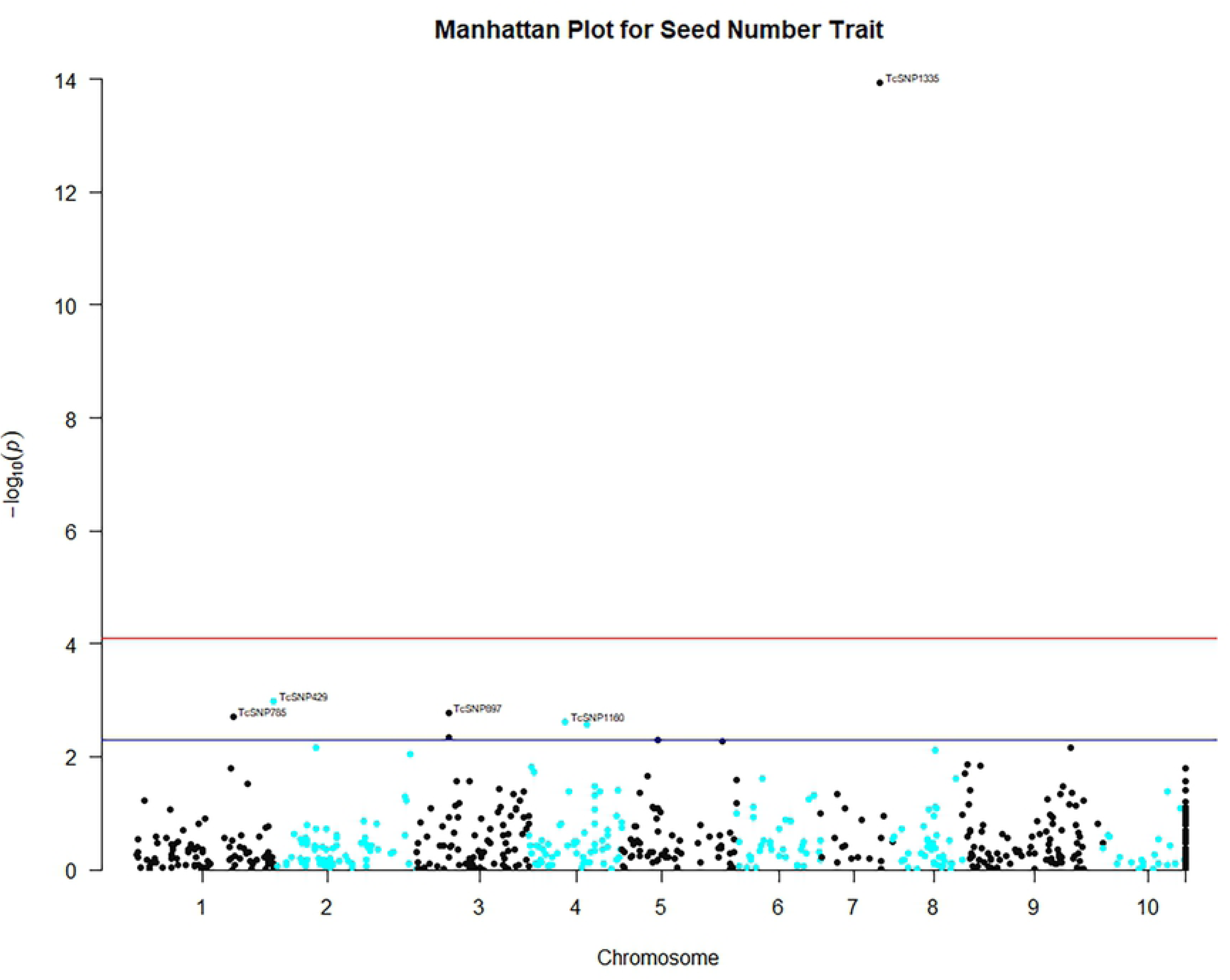
Manhattan plots from genome-wide association analysis. **Legend**: Genome-wide association plots across 8 cacao chromosomes for seven phenotypic traits that had statistically significant MTAs: filament anthocyanin intensity, fruit surface (ridges) anthocyanin intensity, log fruit length, log seed length, log seed number, seed length to width ratio, seed number.
- Based on TASSEL version 5.2.50 results for 421 cacao accessions, 612 filtered SNPs and the Mixed Linear Model.
- Chromosome “11” was designated for unmapped SNP markers.
- X- and Y-axes represent the SNP markers along each chromosome and the -log10(*P*-value), respectively.
- The red horizontal line corresponds to the Bonferonni significance threshold of P-values ≤ 8.17 × 10-5 (–log10 (P) = 4.088) and the blue line corresponds to a significance level of 0.005.

**Fig 5B.**
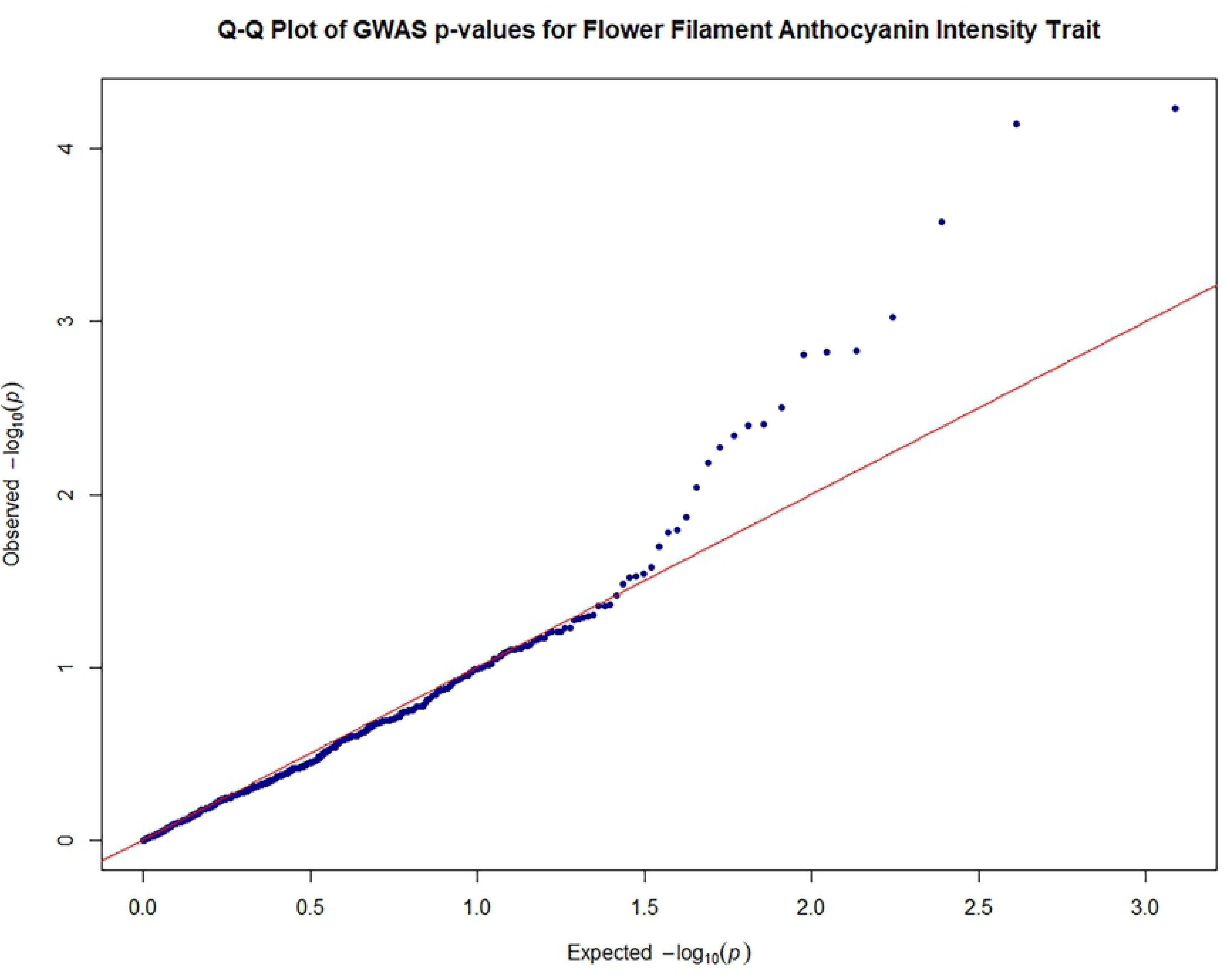
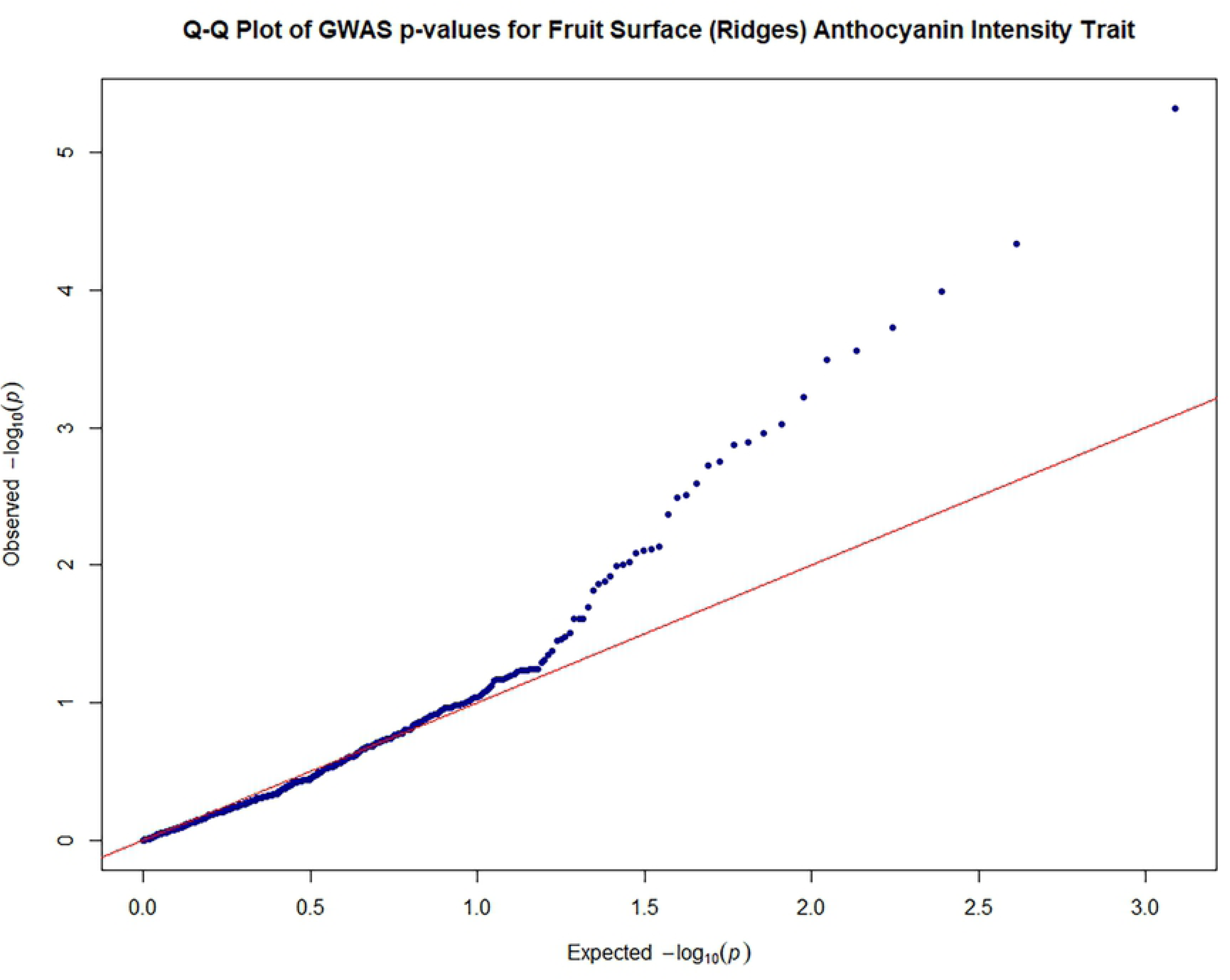
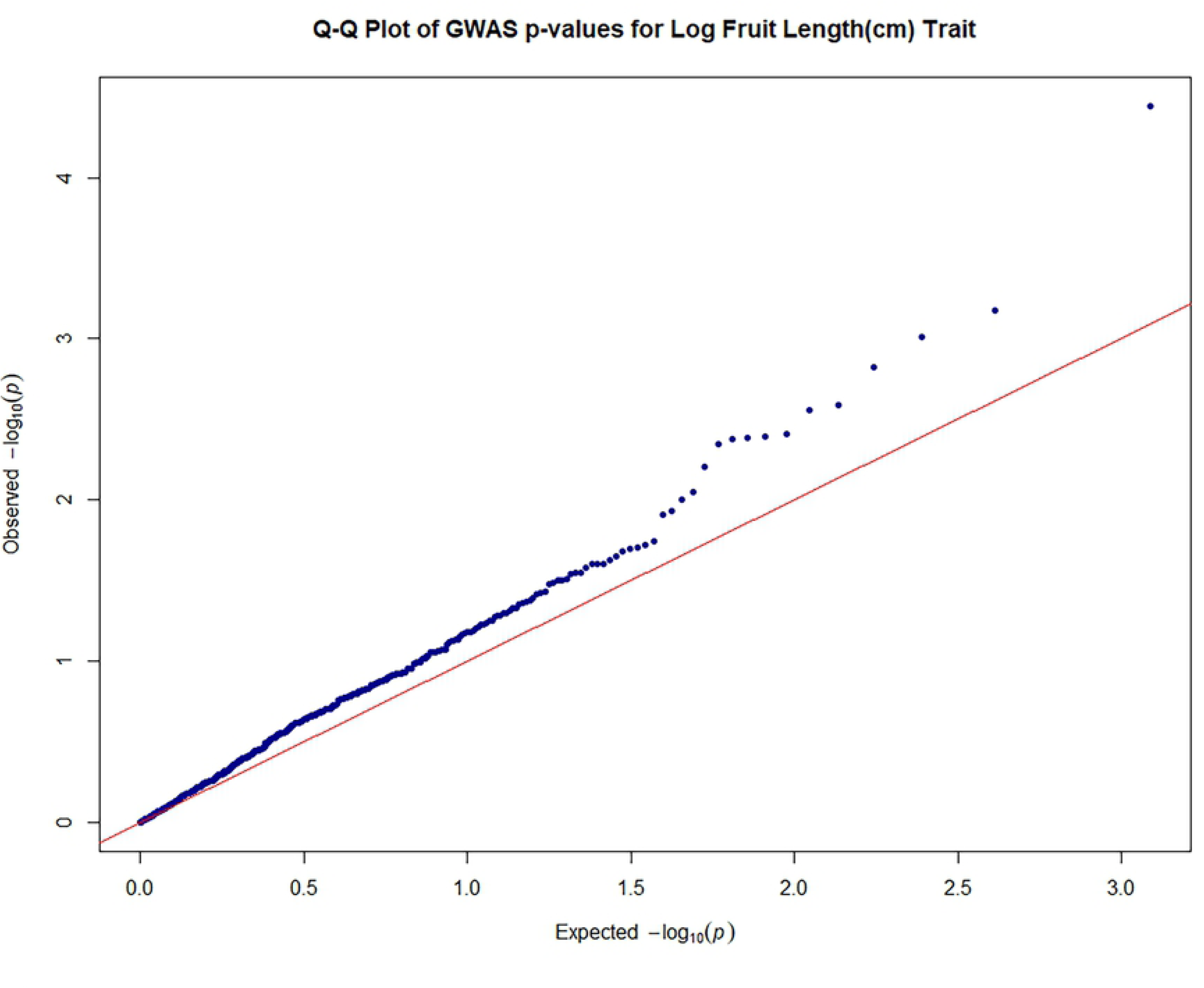
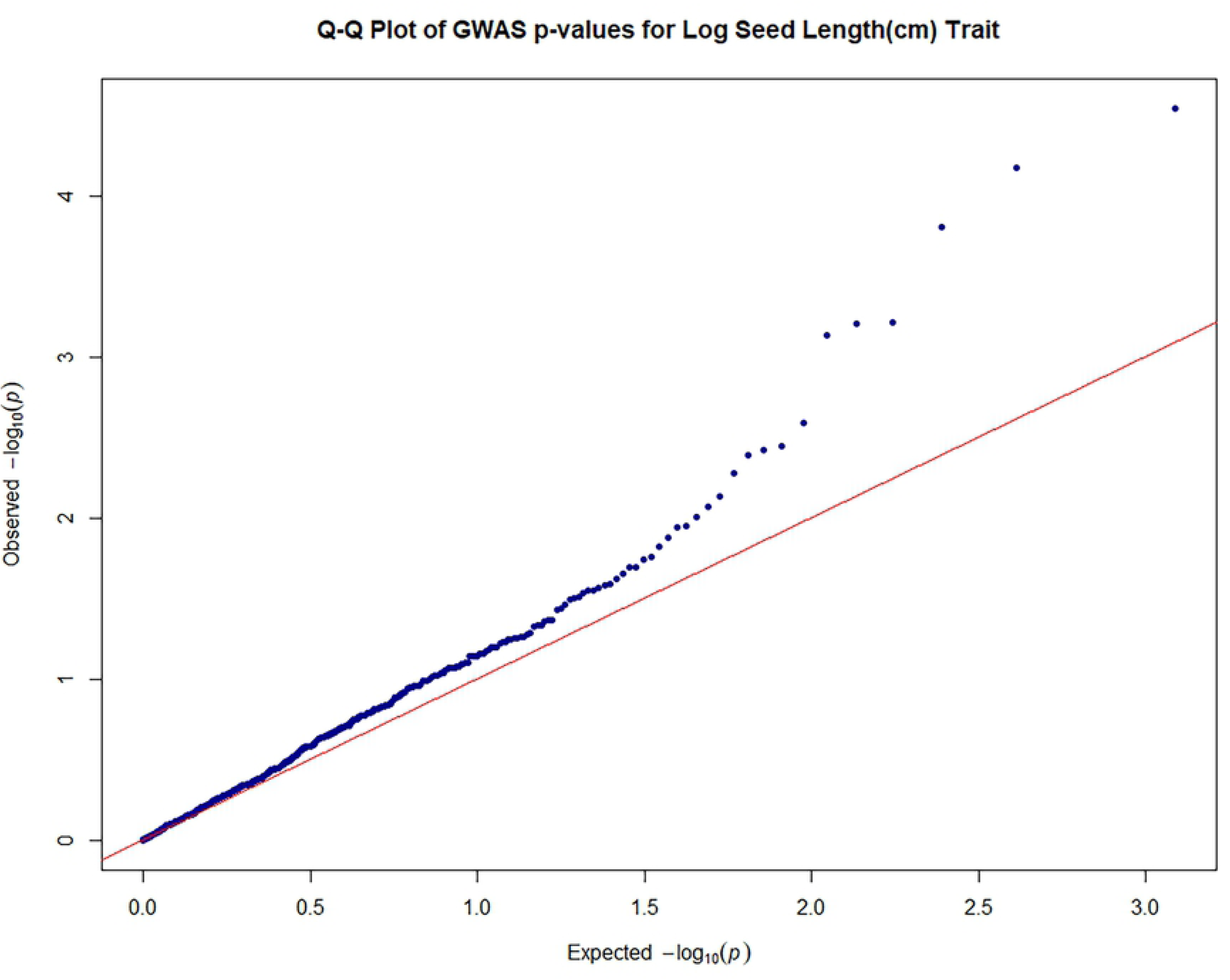
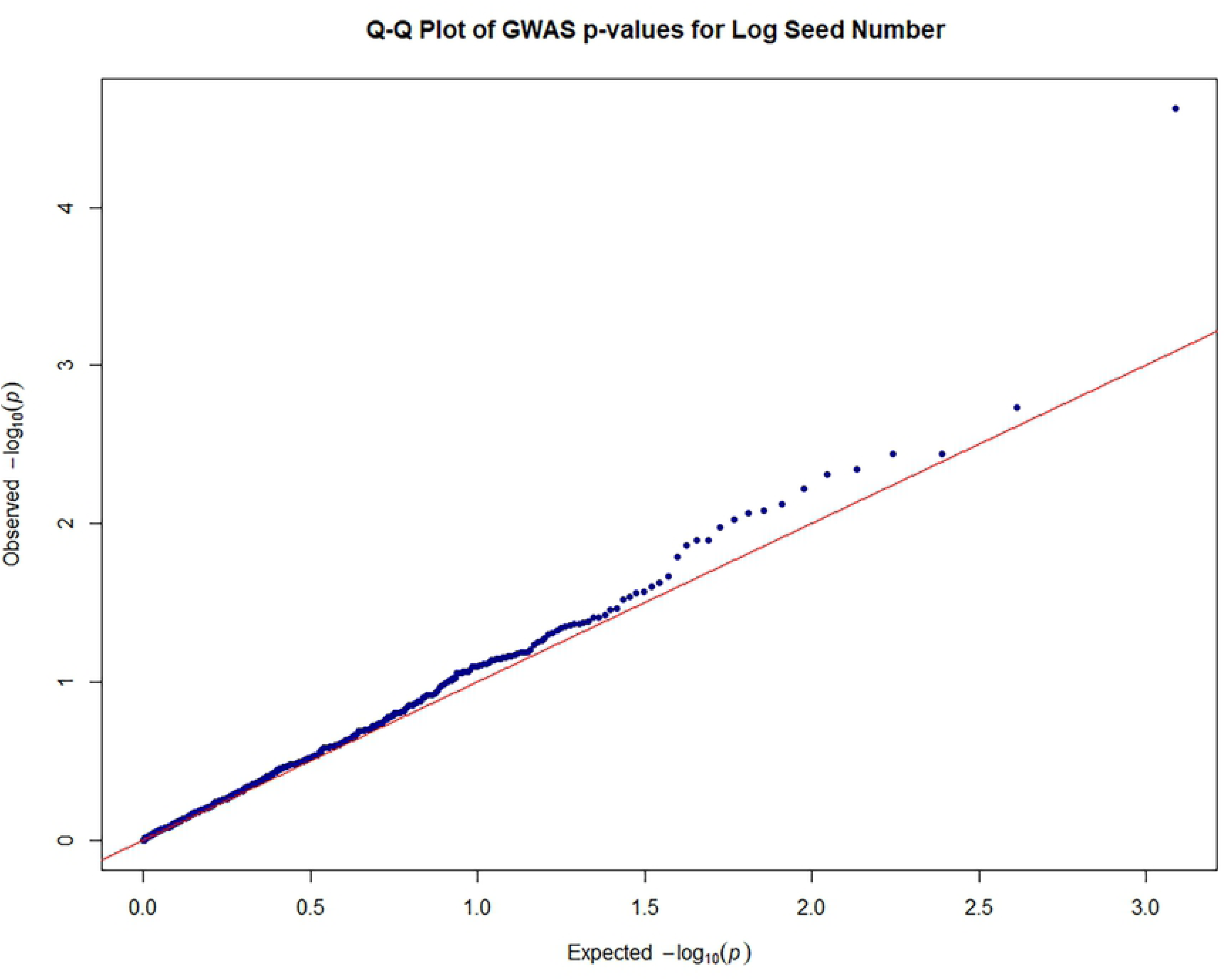
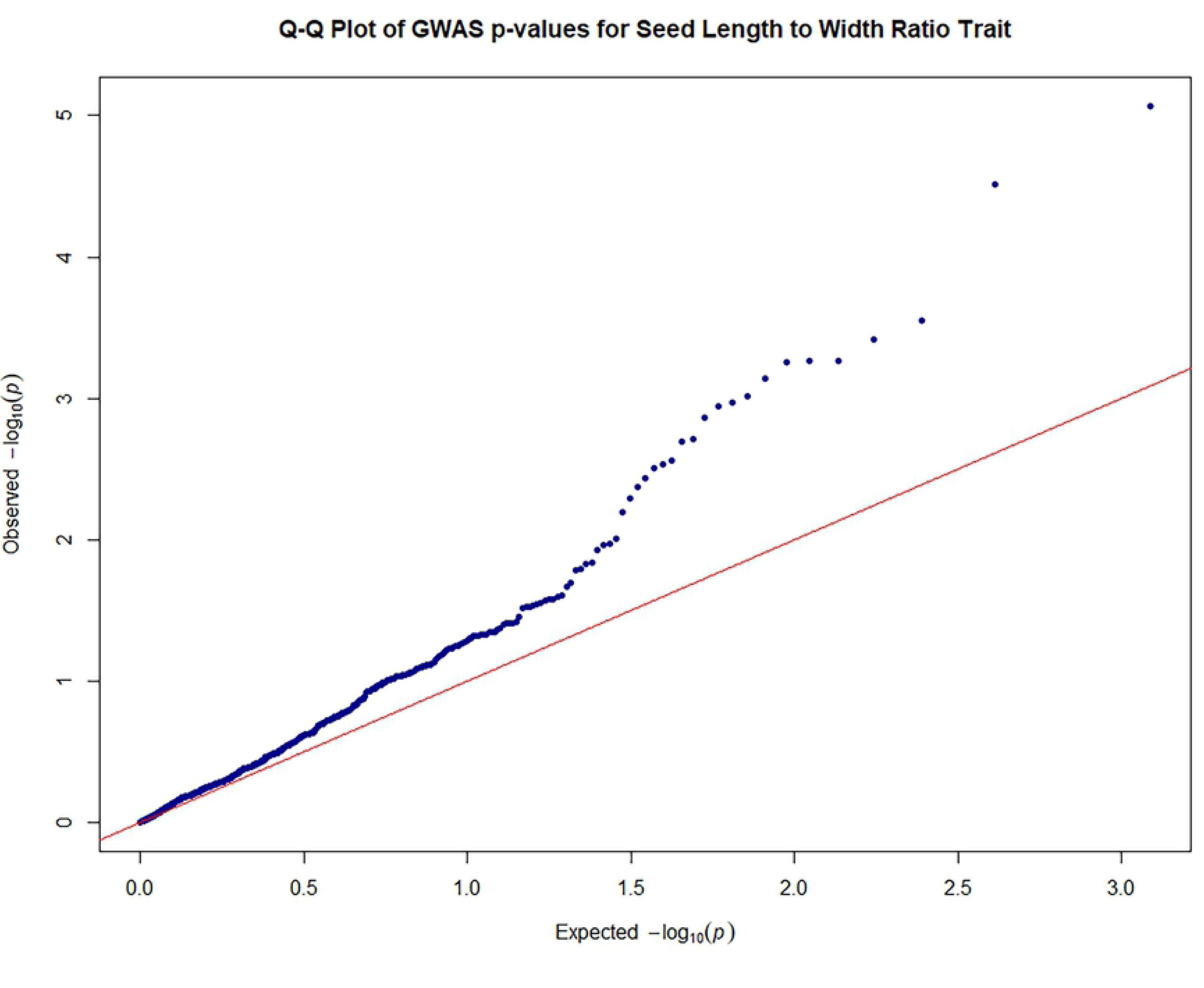
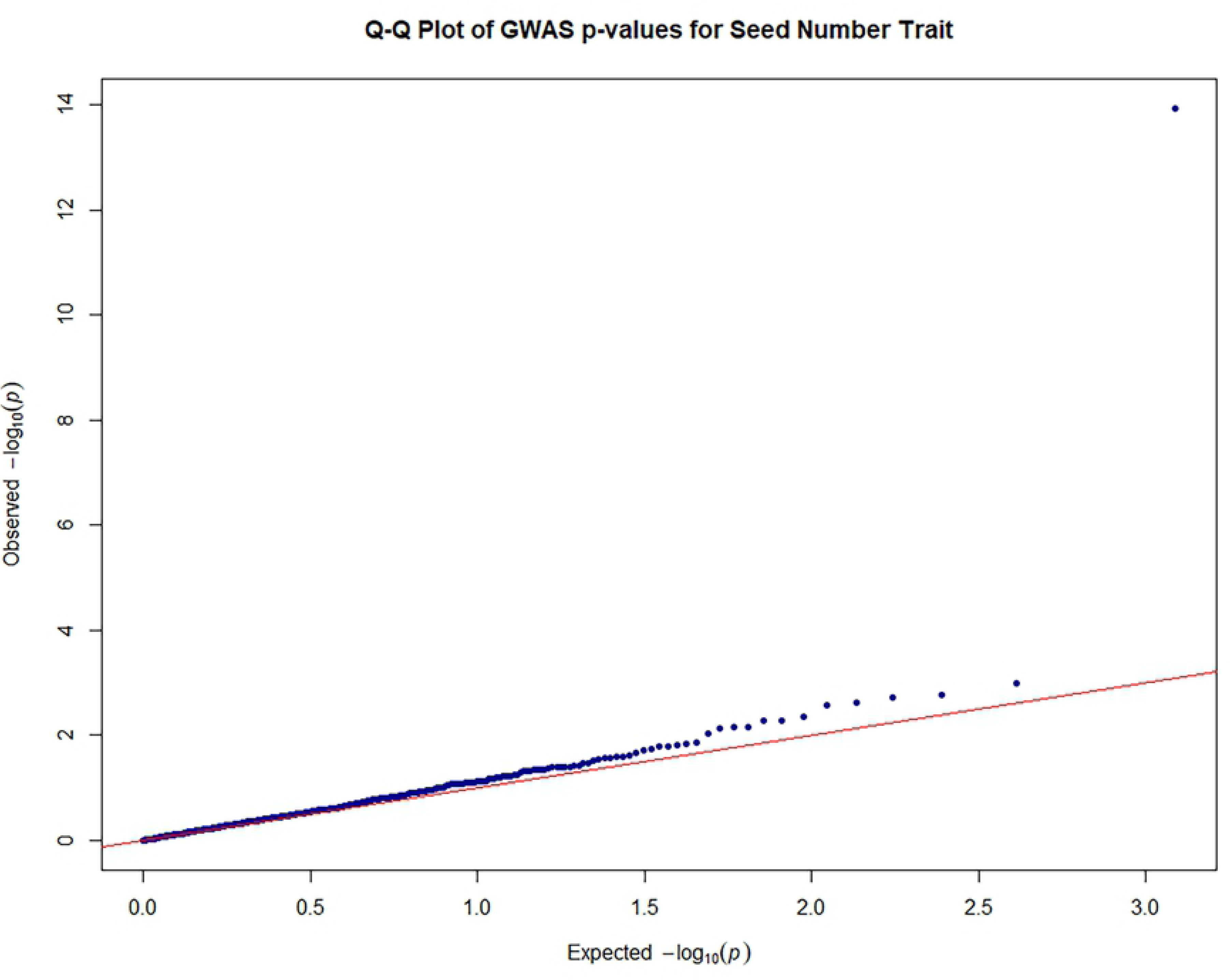
Quantile–quantile plots of estimated−log10 (*P*) from genome-wide association studies. Quantile–quantile plots of estimated−log10 (*P*) for filament anthocyanin intensity; Quantile–quantile plots of estimated−log10 (*P*) for fruit surface (ridges) anthocyanin intensity; Quantile–quantile plots of estimated−log10 (*P*) for log fruit length; Quantile–quantile plots of estimated−log10 (*P*) for log seed length; Quantile–quantile plots of estimated−log10 (*P*) for log seed number; Quantile–quantile plots of estimated−log10 (*P*) for seed length to width ratio; Quantile–quantile plots of estimated−log10 (*P*) for seed number **Legend**: The plots provide no evidence of bias in the GWAS, such as due to genotyping artifacts, and display the extent to which the observed distribution of the test statistic followed the expected (null) distribution The red line represents expected *P*-values with no associations.

**Table 6.** Most significant, yield-related and other marker-trait associations and variation explained

| Trait | Marker | Chr | Position | Df | F | P | Additive Effect | Additive P | Dominant Effect | Dominant P | Marker R<sup>2</sup> |
| --- | --- | --- | --- | --- | --- | --- | --- | --- | --- | --- | --- |
| Flower filament anthocyanin intensity | TcSNP1183 | 3 | 32,973,816 | 2 | 10.01 | 5.84E<sup>-05</sup> | 0.84 | 7.90E<sup>-04</sup> | -0.59 | 0.0517 | 0.049 |
| Flower filament anthocyanin intensity | TcSNP1441 | 8 | 25,585,284 | 2 | 9.8 | 7.14E<sup>-05</sup> | -1.55 | 2.15E<sup>-05</sup> | -0.55 | 0.199 | 0.048 |
| Fruit surface (ridges) anthocyanin intensity | TcSNP401 | 4 | 20,485,872 | 2 | 12.73 | 4.78E<sup>-06</sup> | 0.76 | 0.002228 | 0.41 | 0.1923 | 0.071 |
| Fruit surface (ridges) anthocyanin intensity | TcSNP644 | 9 | 1,220,437 | 2 | 10.31 | 4.57E<sup>-05</sup> | -0.21 | 0.5374 | -1.54 | 4.69E<sup>-05</sup> | 0.059 |
| Fruit shape orbicular | TcSNP1353 | 3 | 34,216,420 | 2 | 19.96 | 6.73E<sup>-09</sup> | 0.09 | 8.80E<sup>-10</sup> | -0.09 | 8.80E<sup>-10</sup> | 0.109 |
| Fruit shape oblate | TcSNP1353 | 3 | 34,216,420 | 2 | 19.96 | 6.73E<sup>-09</sup> | 0.09 | 8.80E<sup>-10</sup> | -0.09 | 8.80E<sup>-10</sup> | 0.109 |
| Fruit shape orbicular | TcSNP1477 | 3 | 17,127,233 | 2 | 12.15 | 8.20E<sup>-06</sup> | 0.07 | 1.33E<sup>-06</sup> | -0.07 | 1.33E<sup>-06</sup> | 0.069 |
| Fruit shape oblate | TcSNP1477 | 3 | 17,127,233 | 2 | 12.15 | 8.20E<sup>-06</sup> | 0.07 | 1.33E<sup>-06</sup> | -0.07 | 1.33E<sup>-06</sup> | 0.069 |
| **Fruit length** | TcSNP1110 | 5 | 1,969,054 | 1 | 16.79 | 5.29E<sup>-05</sup> | NaN | NaN | NaN | NaN | 0.047 |
| **Log fruit length** | TcSNP1110 | 5 | 1,969,054 | 1 | 17.56 | 3.60E<sup>-05</sup> | NaN | NaN | NaN | NaN | 0.049 |
| **Log seed length** | TcSNP344 | 4 | 8,445,308 | 1 | 18.01 | 2.88E<sup>-05</sup> | NaN | NaN | NaN | NaN | 0.050 |
| **Seed length: width** | TcSNP897 | 3 | 23,362,296 | 2 | 12.09 | 8.57E<sup>-06</sup> | -0.11 | 4.96E<sup>-04</sup> | 0.19 | 3.24E<sup>-06</sup> | 0.067 |
| **Seed length: width** | TcSNP733 | 5 | 2,070,839 | 1 | 17.89 | 3.05E<sup>-05</sup> | NaN | NaN | NaN | NaN | 0.051 |
| **Seed number** | TcSNP1335 | 7 | 9,085,336 | 2 | 35.51 | 1.15E<sup>-14</sup> | 20.04 | 8.93E<sup>-13</sup> | 23.42 | 1.18E<sup>-15</sup> | 0.170 |
| **Log seed number** | TcSNP785 | 1 | 31,785,158 | 2 | 11 | 2.38E<sup>-05</sup> | -0.06 | 2.17E<sup>-04</sup> | 0.07 | 1.88E<sup>-04</sup> | 0.059 |
| **Log Seed length** | TcSNP1335 | 7 | 9,085,336 | 2 | 9.89 | 6.75E<sup>-05</sup> | -0.20 | 1.21E<sup>-05</sup> | -0.18 | 6.91E<sup>-05</sup> | 0.055 |
| Seed length: width | TcSNP823 | 5 | 2,546,863 | 2 | 9.49 | **9.97E<sup>-05</sup>** | -0.01 | 0.3114 | 0.05 | 2.90E<sup>-05</sup> | 0.053 |
| Seed number | TcSNP1160 | 4 | 21,847,012 | 1 | 116.1 | **1.00E<sup>-04</sup>** | NaN | 2.15E<sup>-23</sup> | NaN | NaN | 0.261 |
| Pod Index | TcSNP642 | 8 | 16,399,147 | 1 | 13.88 | **2.30E<sup>-04</sup>** | NaN | NaN | NaN | 330 | 0.219 |

**P values below the Bonferroni correction threshold are in bold Chr = chromosome**
**Bonferroni threshold $\log_{10} 8.17 \times 10^{-5} = -4.0877$**

Consequently, the results were carefully scrutinized to discern MTAs just below the level of significance, which may also have functional importance.

Nine putative candidate genes with functional roles related to seed development, lipid biosynthesis and transfer and carbohydrate transport were identified on chromosome 1 (Table 7). It is noteworthy that TcSNP 785 on chromosome 1 was co-localized with gene *Tc01v2_g025880*, which is functionally significant since it encodes protein disulfide isomerase that may be required for proper pollen development, ovule fertilization and embryo development (Table 7). On chromosome 3, 11 putative candidate genes were detected, which encode traits associated with seed development and lipid accumulation (Table 7).

**Table 7.**
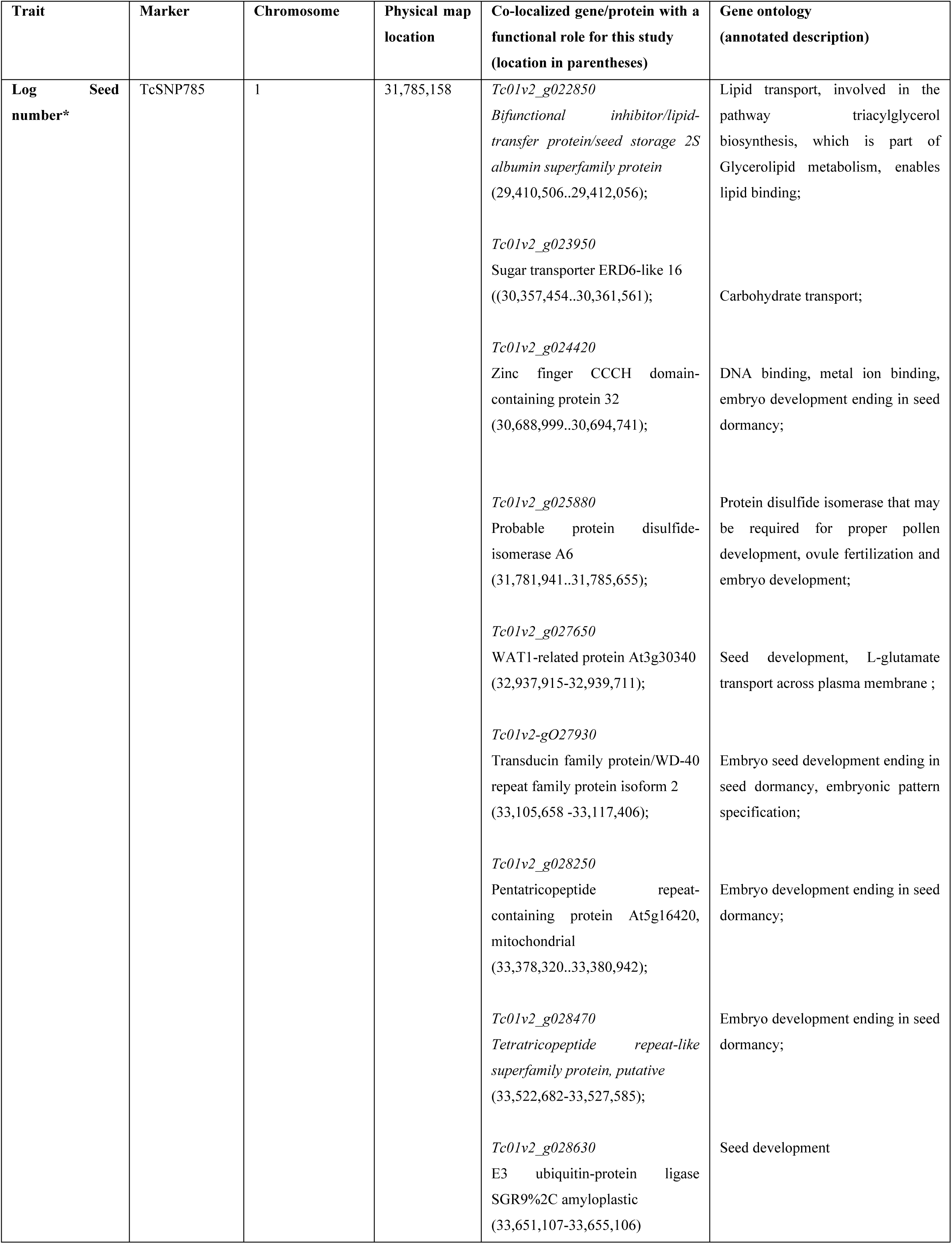
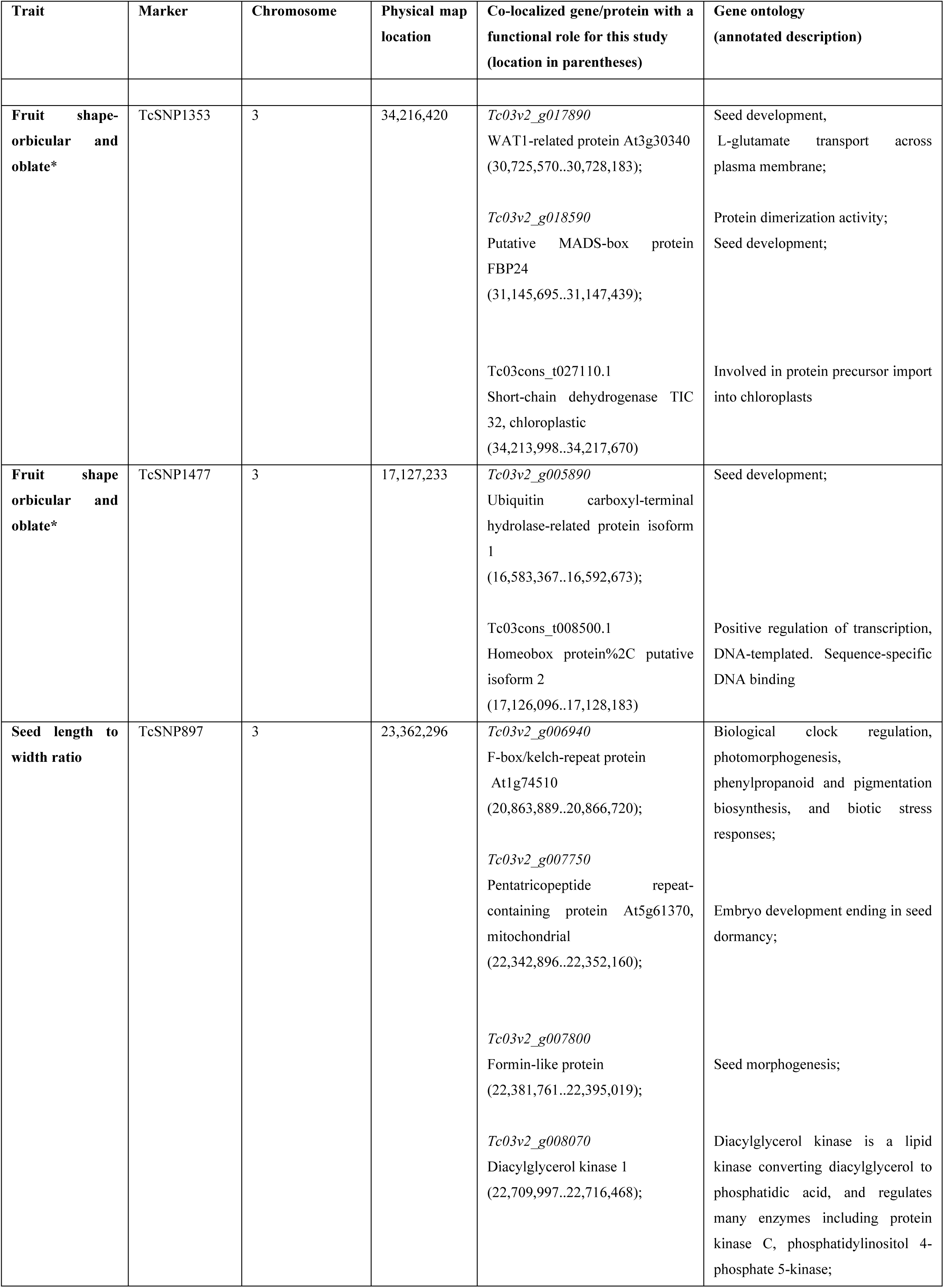
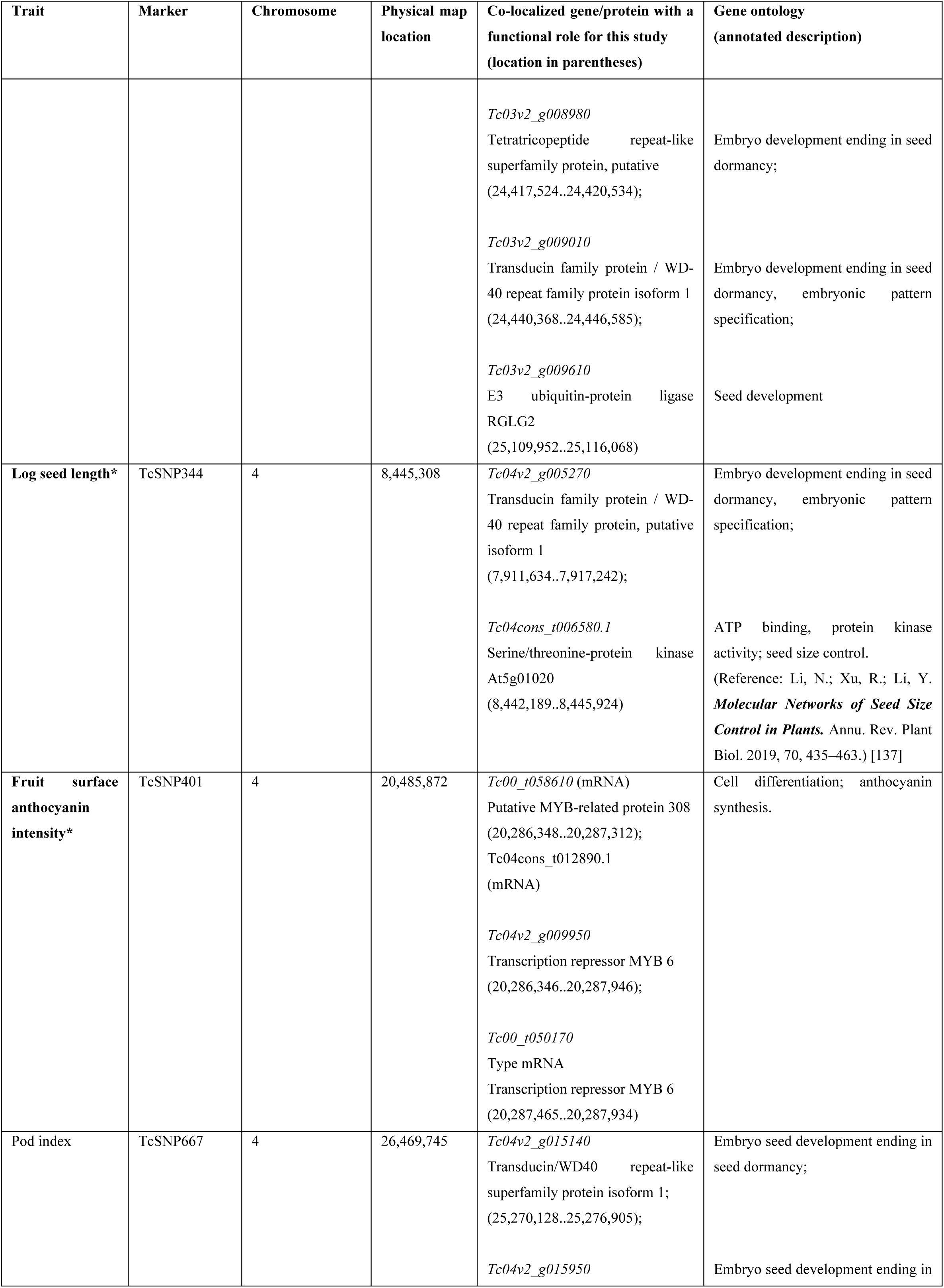
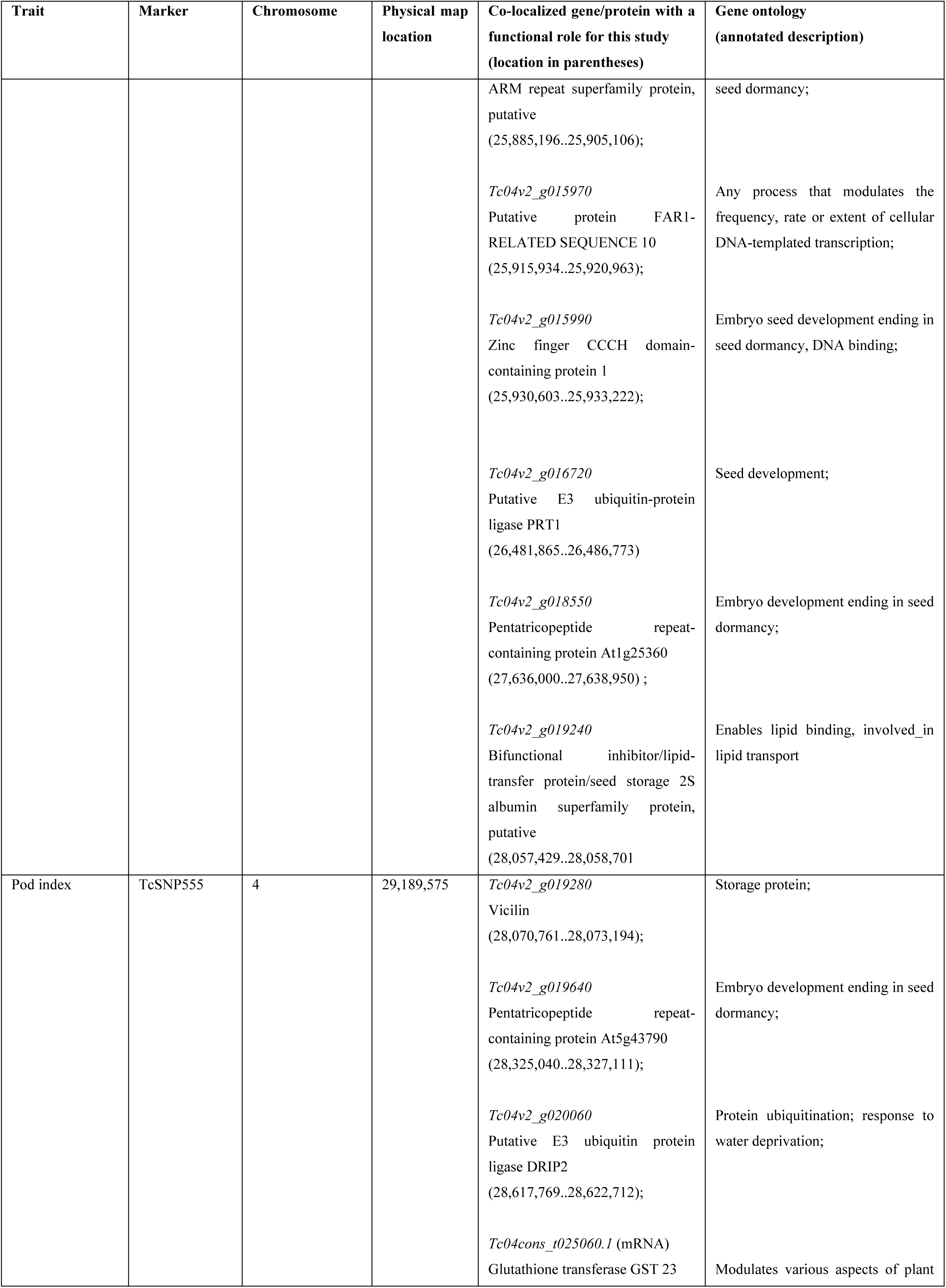
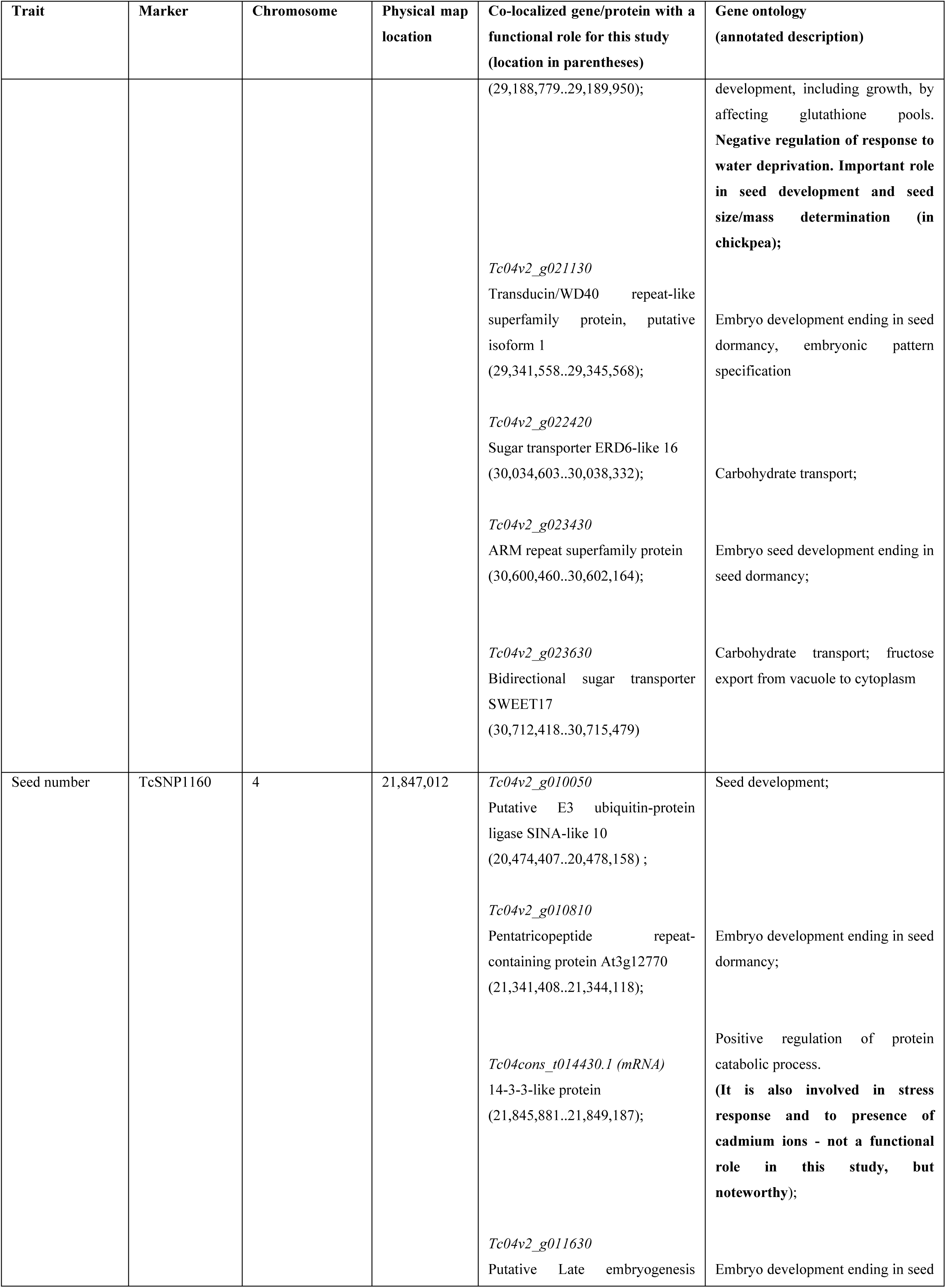
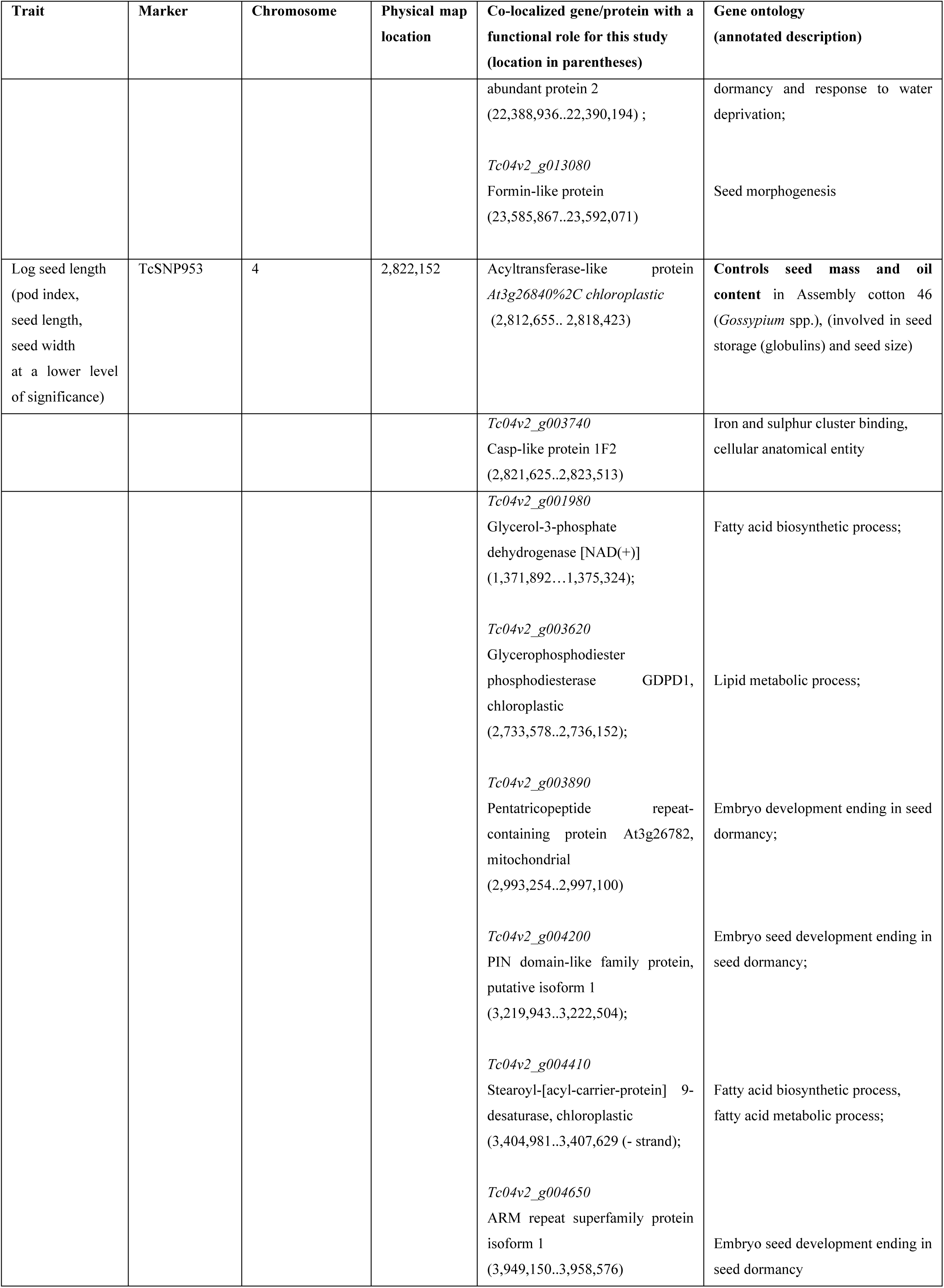
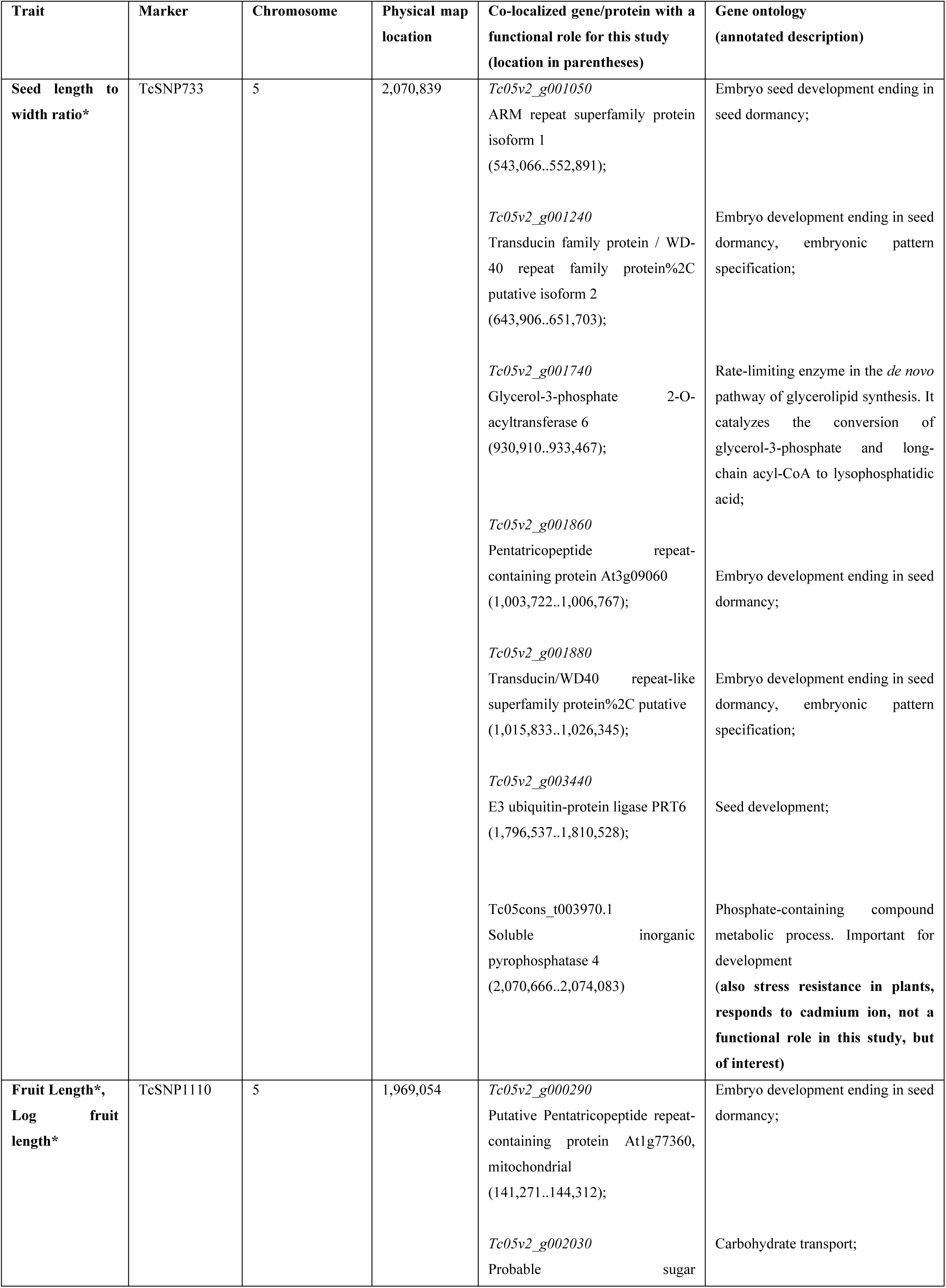
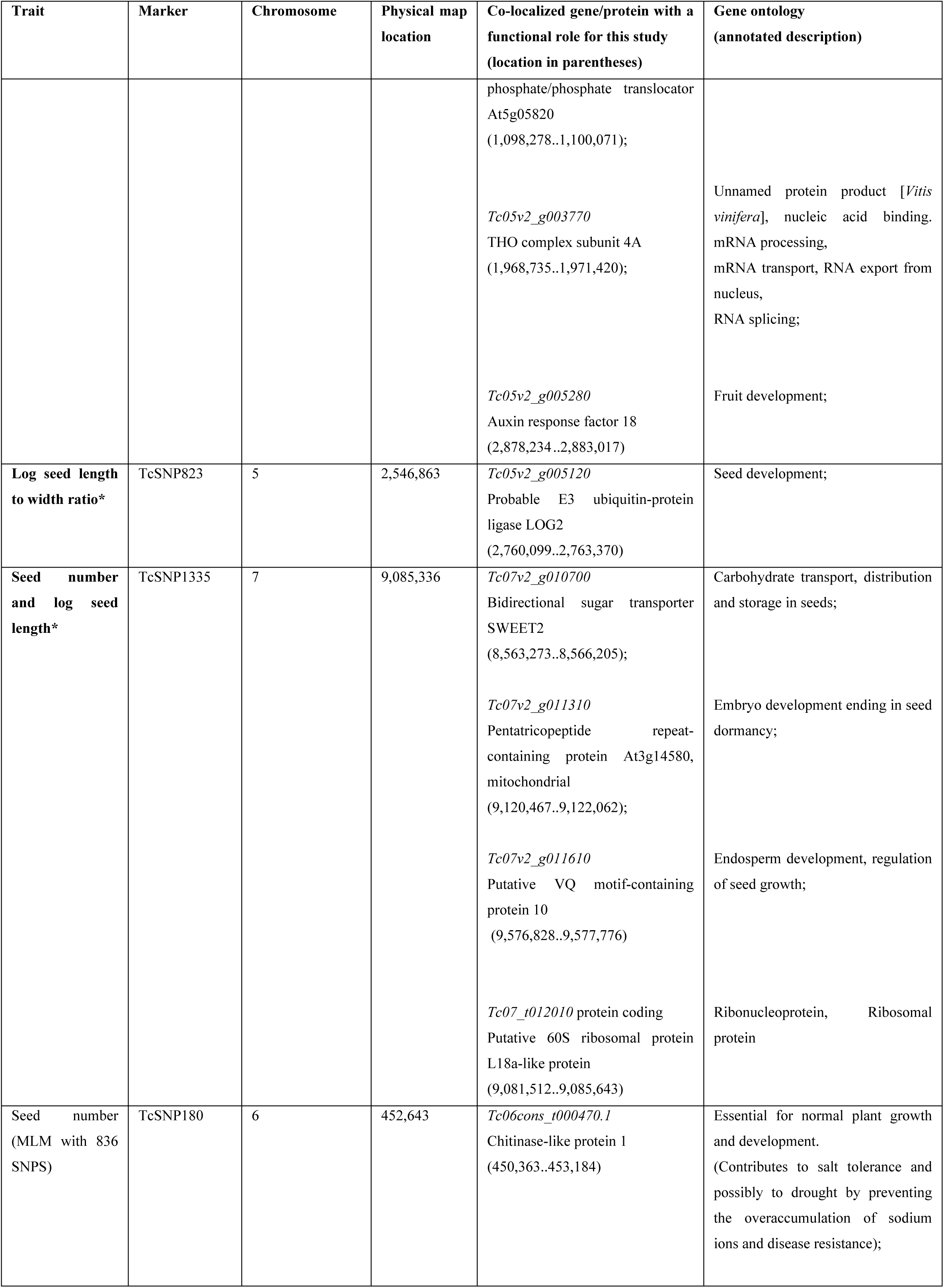
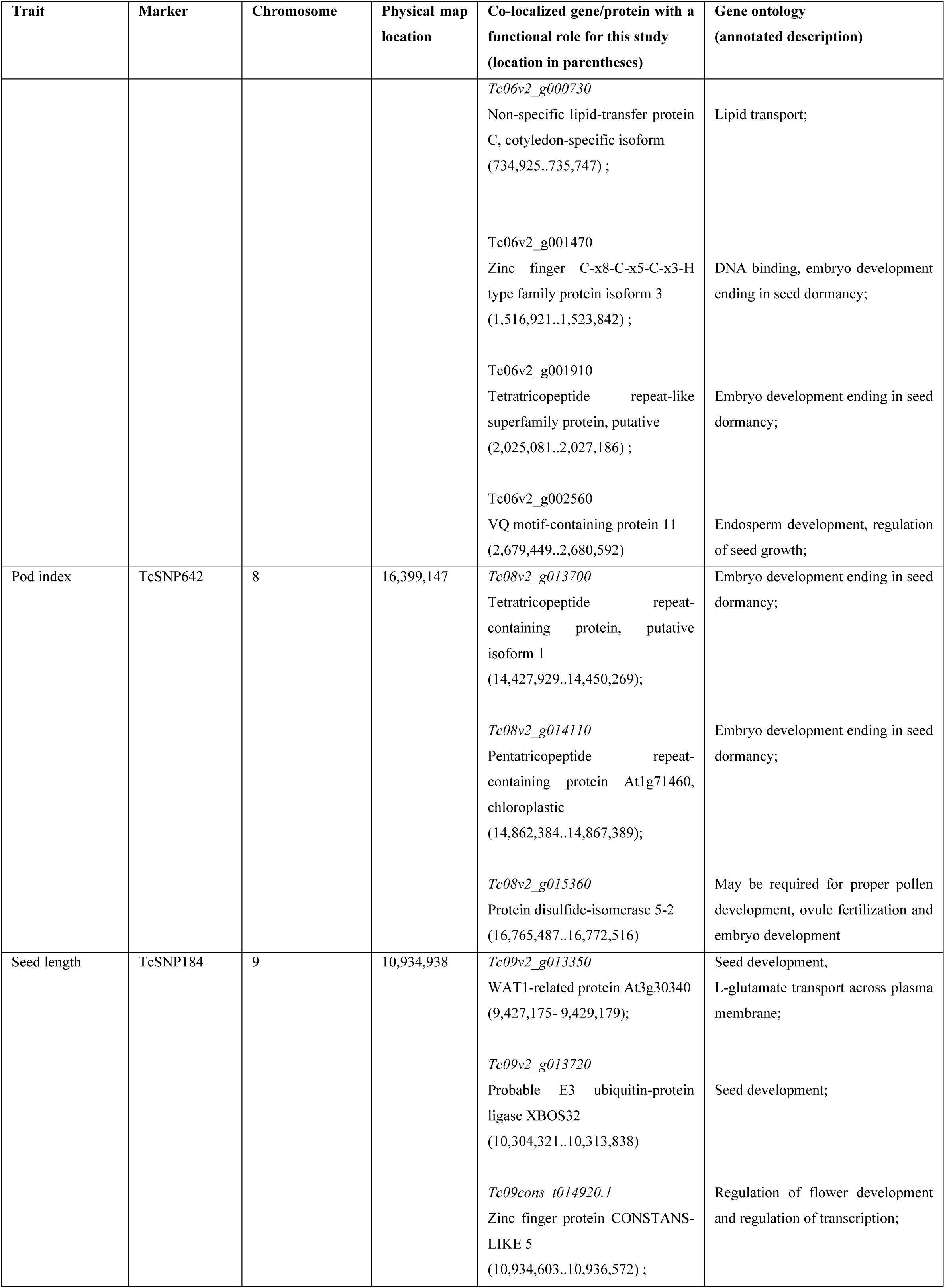
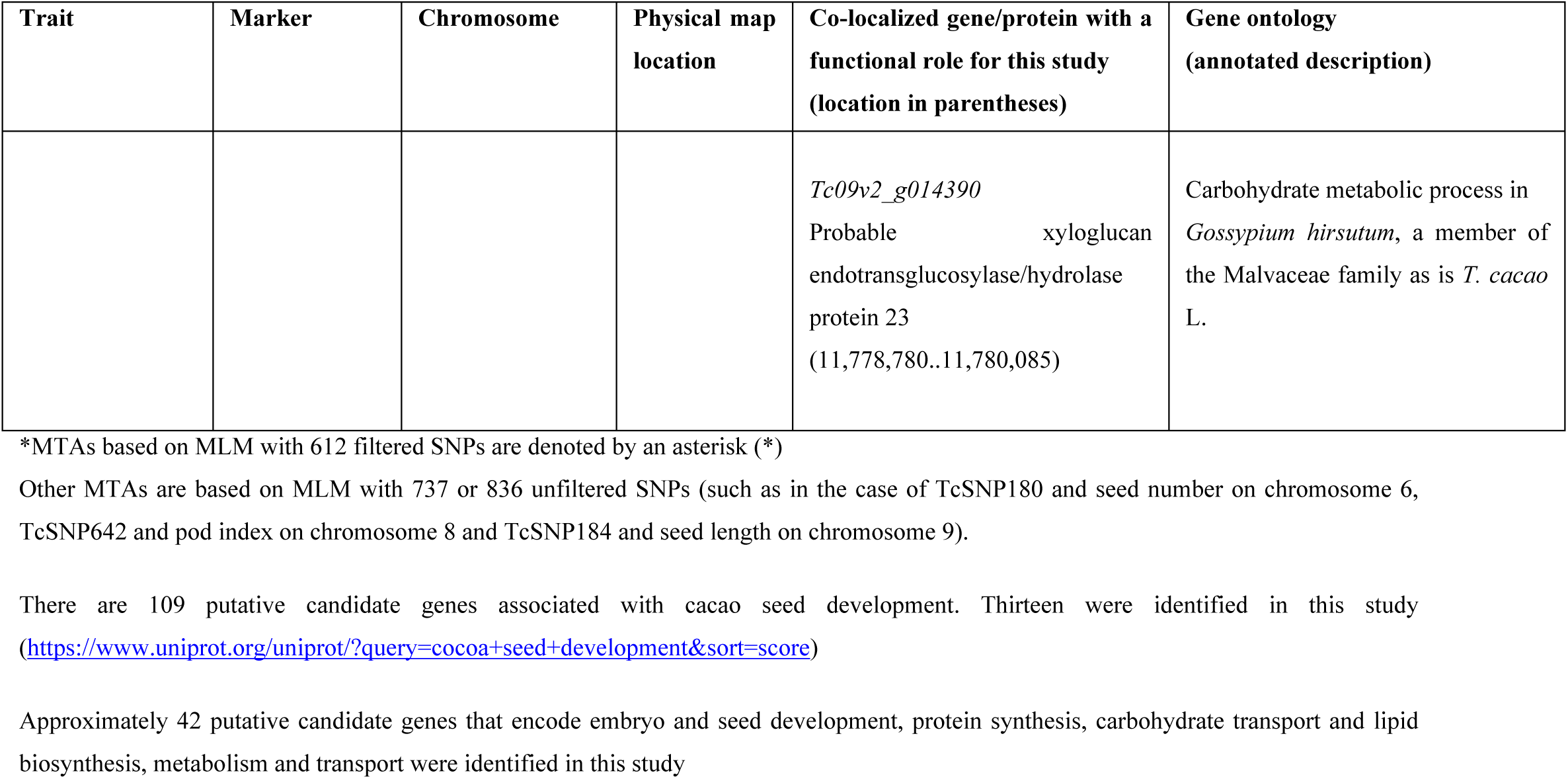
Genes co-localized with SNP markers significantly associated with phenotypic traits.

| Trait | Marker | Chromosome | Physical map location | Co-localized gene/protein with a functional role for this study (location in parentheses) | Gene ontology (annotated description) |
| --- | --- | --- | --- | --- | --- |
| Log Seed number* | TcSNP785 | 1 | 31,785,158 | *Tc01v2_g022850*<br>Bifunctional inhibitor/lipid-transfer protein/seed storage 2S albumin superfamily protein (29,410,506..29,412,056);<br>*Tc01v2_g023950*<br>Sugar transporter ERD6-like 16 ((30,357,454..30,361,561);<br>*Tc01v2_g024420*<br>Zinc finger CCCH domain-containing protein 32 (30,688,999..30,694,741);<br>*Tc01v2_g025880*<br>Probable protein disulfide-isomerase A6 (31,781,941..31,785,655);<br>*Tc01v2_g027650*<br>WAT1-related protein At3g30340 (32,937,915-32,939,711);<br>*Tc01v2-gO27930*<br>Transducin family protein/WD-40 repeat family protein isoform 2 (33,105,658 -33,117,406);<br>*Tc01v2_g028250*<br>Pentatricopeptide repeat-containing protein At5g16420, mitochondrial (33,378,320..33,380,942);<br>*Tc01v2_g028470*<br>Tetratricopeptide repeat-like superfamily protein, putative (33,522,682-33,527,585);<br>*Tc01v2_g028630*<br>E3 ubiquitin-protein ligase SGR9%2C amyloplastic (33,651,107-33,655,106) | Lipid transport, involved in the pathway triacylglycerol biosynthesis, which is part of Glycerolipid metabolism, enables lipid binding;<br>Carbohydrate transport;<br>DNA binding, metal ion binding, embryo development ending in seed dormancy;<br>Protein disulfide isomerase that may be required for proper pollen development, ovule fertilization and embryo development;<br>Seed development, L-glutamate transport across plasma membrane ;<br>Embryo seed development ending in seed dormancy, embryonic pattern specification;<br>Embryo development ending in seed dormancy;<br>Embryo development ending in seed dormancy;<br>Seed development |
| **Fruit shape-orbicular and oblate\*** | TcSNP1353 | 3 | 34,216,420 | *Tc03v2_g017890*<br>WAT1-related protein At3g30340 (30,725,570..30,728,183);<br>*Tc03v2_g018590*<br>Putative MADS-box protein FBP24 (31,145,695..31,147,439);<br>Tc03cons_t027110.1<br>Short-chain dehydrogenase TIC 32, chloroplastic (34,213,998..34,217,670) | Seed development, L-glutamate transport across plasma membrane;<br>Protein dimerization activity; Seed development;<br>Involved in protein precursor import into chloroplasts |
| **Fruit shape orbicular and oblate\*** | TcSNP1477 | 3 | 17,127,233 | *Tc03v2_g005890*<br>Ubiquitin carboxyl-terminal hydrolase-related protein isoform 1 (16,583,367..16,592,673);<br>Tc03cons_t008500.1<br>Homeobox protein%2C putative isoform 2 (17,126,096..17,128,183) | Seed development;<br>Positive regulation of transcription, DNA-templated. Sequence-specific DNA binding |
| **Seed length to width ratio** | TcSNP897 | 3 | 23,362,296 | *Tc03v2_g006940*<br>F-box/kelch-repeat protein At1g74510 (20,863,889..20,866,720);<br>*Tc03v2_g007750*<br>Pentatricopeptide repeat-containing protein At5g61370, mitochondrial (22,342,896..22,352,160);<br>*Tc03v2_g007800*<br>Formin-like protein (22,381,761..22,395,019);<br>*Tc03v2_g008070*<br>Diacylglycerol kinase 1 (22,709,997..22,716,468); | Biological clock regulation, photomorphogenesis, phenylpropanoid and pigmentation biosynthesis, and biotic stress responses;<br>Embryo development ending in seed dormancy;<br>Seed morphogenesis;<br>Diacylglycerol kinase is a lipid kinase converting diacylglycerol to phosphatidic acid, and regulates many enzymes including protein kinase C, phosphatidylinositol 4-phosphate 5-kinase; |
|  |  |  |  | *Tc03v2_g008980*<br>Tetratricopeptide repeat-like superfamily protein, putative (24,417,524..24,420,534);<br>*Tc03v2_g009010*<br>Transducin family protein / WD-40 repeat family protein isoform 1 (24,440,368..24,446,585);<br>*Tc03v2_g009610*<br>E3 ubiquitin-protein ligase RGLG2 (25,109,952..25,116,068) | Embryo development ending in seed dormancy;<br>Embryo development ending in seed dormancy, embryonic pattern specification;<br>Seed development |
| Log seed length* | TcSNP344 | 4 | 8,445,308 | *Tc04v2_g005270*<br>Transducin family protein / WD-40 repeat family protein, putative isoform 1 (7,911,634..7,917,242);<br>*Tc04cons_t006580.1*<br>Serine/threonine-protein kinase At5g01020 (8,442,189..8,445,924) | Embryo development ending in seed dormancy, embryonic pattern specification;<br>ATP binding, protein kinase activity; seed size control. (Reference: Li, N.; Xu, R.; Li, Y. *Molecular Networks of Seed Size Control in Plants*. Annu. Rev. Plant Biol. 2019, 70, 435–463.) [137] |
| Fruit surface anthocyanin intensity* | TcSNP401 | 4 | 20,485,872 | *Tc00_t058610* (mRNA)<br>Putative MYB-related protein 308 (20,286,348..20,287,312);<br>*Tc04cons_t012890.1* (mRNA)<br>*Tc04v2_g009950*<br>Transcription repressor MYB 6 (20,286,346..20,287,946);<br>*Tc00_t050170*<br>Type mRNA<br>Transcription repressor MYB 6 (20,287,465..20,287,934) | Cell differentiation; anthocyanin synthesis. |
| Pod index | TcSNP667 | 4 | 26,469,745 | *Tc04v2_g015140*<br>Transducin/WD40 repeat-like superfamily protein isoform 1; (25,270,128..25,276,905);<br>*Tc04v2_g015950* | Embryo seed development ending in seed dormancy;<br>Embryo seed development ending in |
|  |  |  |  | ARM repeat superfamily protein, putative<br>(25,885,196..25,905,106);<br>*Tc04v2_g015970*<br>Putative protein FAR1-RELATED SEQUENCE 10<br>(25,915,934..25,920,963);<br>*Tc04v2_g015990*<br>Zinc finger CCCH domain-containing protein 1<br>(25,930,603..25,933,222);<br>*Tc04v2_g016720*<br>Putative E3 ubiquitin-protein ligase PRT1<br>(26,481,865..26,486,773)<br>*Tc04v2_g018550*<br>Pentatricopeptide repeat-containing protein At1g25360<br>(27,636,000..27,638,950) ;<br>*Tc04v2_g019240*<br>Bifunctional inhibitor/lipid-transfer protein/seed storage 2S albumin superfamily protein, putative<br>(28,057,429..28,058,701) | seed dormancy;<br>Any process that modulates the frequency, rate or extent of cellular DNA-templated transcription;<br>Embryo seed development ending in seed dormancy, DNA binding;<br>Seed development;<br>Embryo development ending in seed dormancy;<br>Enables lipid binding, involved in lipid transport |
| Pod index | TcSNP555 | 4 | 29,189,575 | *Tc04v2_g019280*<br>Vicilin<br>(28,070,761..28,073,194);<br>*Tc04v2_g019640*<br>Pentatricopeptide repeat-containing protein At5g43790<br>(28,325,040..28,327,111);<br>*Tc04v2_g020060*<br>Putative E3 ubiquitin protein ligase DRIP2<br>(28,617,769..28,622,712);<br>*Tc04cons_t025060.1* (mRNA)<br>Glutathione transferase GST 23 | Storage protein;<br>Embryo development ending in seed dormancy;<br>Protein ubiquitination; response to water deprivation;<br>Modulates various aspects of plant |
|  |  |  |  | (29,188,779..29,189,950);<br>*Tc04v2_g021130*<br>Transducin/WD40 repeat-like superfamily protein, putative isoform 1<br>(29,341,558..29,345,568);<br>*Tc04v2_g022420*<br>Sugar transporter ERD6-like 16<br>(30,034,603..30,038,332);<br>*Tc04v2_g023430*<br>ARM repeat superfamily protein<br>(30,600,460..30,602,164);<br>*Tc04v2_g023630*<br>Bidirectional sugar transporter SWEET17<br>(30,712,418..30,715,479) | development, including growth, by affecting glutathione pools. **Negative regulation of response to water deprivation. Important role in seed development and seed size/mass determination (in chickpea);**<br>Embryo development ending in seed dormancy, embryonic pattern specification<br>Carbohydrate transport;<br>Embryo seed development ending in seed dormancy;<br>Carbohydrate transport; fructose export from vacuole to cytoplasm |
| Seed number | TcSNP1160 | 4 | 21,847,012 | *Tc04v2_g010050*<br>Putative E3 ubiquitin-protein ligase SINA-like 10<br>(20,474,407..20,478,158) ;<br>*Tc04v2_g010810*<br>Pentatricopeptide repeat-containing protein At3g12770<br>(21,341,408..21,344,118);<br>*Tc04cons_t014430.1 (mRNA)*<br>14-3-3-like protein<br>(21,845,881..21,849,187);<br>*Tc04v2_g011630*<br>Putative Late embryogenesis | Seed development;<br>Embryo development ending in seed dormancy;<br>Positive regulation of protein catabolic process.<br>**(It is also involved in stress response and to presence of cadmium ions - not a functional role in this study, but noteworthy);**<br>Embryo development ending in seed |
|  |  |  |  | abundant protein 2<br>(22,388,936..22,390,194) ;<br>*Tc04v2_g013080*<br>Formin-like protein<br>(23,585,867..23,592,071) | dormancy and response to water deprivation;<br>Seed morphogenesis |
| Log seed length (pod index, seed length, seed width at a lower level of significance) | TcSNP953 | 4 | 2,822,152 | Acyltransferase-like protein<br>*At3g26840%2C chloroplastic*<br>(2,812,655..2,818,423) | **Controls seed mass and oil content** in Assembly cotton 46 (*Gossypium* spp.), (involved in seed storage (globulins) and seed size) |
|  |  |  |  | *Tc04v2_g003740*<br>Casp-like protein 1F2<br>(2,821,625..2,823,513) | Iron and sulphur cluster binding, cellular anatomical entity |
|  |  |  |  | *Tc04v2_g001980*<br>Glycerol-3-phosphate dehydrogenase [NAD(+)]<br>(1,371,892...1,375,324);<br>*Tc04v2_g003620*<br>Glycerophosphodiester phosphodiesterase GDPD1, chloroplastic<br>(2,733,578..2,736,152);<br>*Tc04v2_g003890*<br>Pentatricopeptide repeat-containing protein At3g26782, mitochondrial<br>(2,993,254..2,997,100)<br>*Tc04v2_g004200*<br>PIN domain-like family protein, putative isoform 1<br>(3,219,943..3,222,504);<br>*Tc04v2_g004410*<br>Stearoyl-[acyl-carrier-protein] 9-desaturase, chloroplastic<br>(3,404,981..3,407,629 (- strand));<br>*Tc04v2_g004650*<br>ARM repeat superfamily protein isoform 1<br>(3,949,150..3,958,576) | Fatty acid biosynthetic process;<br>Lipid metabolic process;<br>Embryo development ending in seed dormancy;<br>Embryo seed development ending in seed dormancy;<br>Fatty acid biosynthetic process, fatty acid metabolic process;<br>Embryo seed development ending in seed dormancy |
| Seed length to width ratio* | TcSNP733 | 5 | 2,070,839 | *Tc05v2_g001050*<br>ARM repeat superfamily protein isoform 1<br>(543,066..552,891);<br>*Tc05v2_g001240*<br>Transducin family protein / WD-40 repeat family protein%2C putative isoform 2<br>(643,906..651,703);<br>*Tc05v2_g001740*<br>Glycerol-3-phosphate 2-O-acyltransferase 6<br>(930,910..933,467);<br>*Tc05v2_g001860*<br>Pentatricopeptide repeat-containing protein At3g09060<br>(1,003,722..1,006,767);<br>*Tc05v2_g001880*<br>Transducin/WD40 repeat-like superfamily protein%2C putative<br>(1,015,833..1,026,345);<br>*Tc05v2_g003440*<br>E3 ubiquitin-protein ligase PRT6<br>(1,796,537..1,810,528);<br>Tc05cons_t003970.1<br>Soluble inorganic pyrophosphatase 4<br>(2,070,666..2,074,083) | Embryo seed development ending in seed dormancy;<br>Embryo development ending in seed dormancy, embryonic pattern specification;<br>Rate-limiting enzyme in the *de novo* pathway of glycerolipid synthesis. It catalyzes the conversion of glycerol-3-phosphate and long-chain acyl-CoA to lysophosphatidic acid;<br>Embryo development ending in seed dormancy;<br>Embryo development ending in seed dormancy, embryonic pattern specification;<br>Seed development;<br>Phosphate-containing compound metabolic process. Important for development<br>(also stress resistance in plants, responds to cadmium ion, not a functional role in this study, but of interest) |
| Fruit Length*, Log fruit length* | TcSNP1110 | 5 | 1,969,054 | *Tc05v2_g000290*<br>Putative Pentatricopeptide repeat-containing protein At1g77360, mitochondrial<br>(141,271..144,312);<br>*Tc05v2_g002030*<br>Probable sugar | Embryo development ending in seed dormancy;<br>Carbohydrate transport; |
|  |  |  |  | phosphate/phosphate translocator<br>At5g05820<br>(1,098,278..1,100,071);<br>*Tc05v2_g003770*<br>THO complex subunit 4A<br>(1,968,735..1,971,420);<br>*Tc05v2_g005280*<br>Auxin response factor 18<br>(2,878,234..2,883,017) | Unnamed protein product [*Vitis vinifera*], nucleic acid binding.<br>mRNA processing,<br>mRNA transport, RNA export from nucleus,<br>RNA splicing;<br>Fruit development; |
| Log seed length to width ratio* | TcSNP823 | 5 | 2,546,863 | *Tc05v2_g005120*<br>Probable E3 ubiquitin-protein ligase LOG2<br>(2,760,099..2,763,370) | Seed development; |
| Seed number and log seed length* | TcSNP1335 | 7 | 9,085,336 | *Tc07v2_g010700*<br>Bidirectional sugar transporter SWEET2<br>(8,563,273..8,566,205);<br>*Tc07v2_g011310*<br>Pentatricopeptide repeat-containing protein At3g14580, mitochondrial<br>(9,120,467..9,122,062);<br>*Tc07v2_g011610*<br>Putative VQ motif-containing protein 10<br>(9,576,828..9,577,776)<br>*Tc07_t012010* protein coding<br>Putative 60S ribosomal protein L18a-like protein<br>(9,081,512..9,085,643) | Carbohydrate transport, distribution and storage in seeds;<br>Embryo development ending in seed dormancy;<br>Endosperm development, regulation of seed growth;<br>Ribonucleoprotein, Ribosomal protein |
| Seed number (MLM with 836 SNPS) | TcSNP180 | 6 | 452,643 | *Tc06cons_t000470.1*<br>Chitinase-like protein 1<br>(450,363..453,184) | Essential for normal plant growth and development.<br>(Contributes to salt tolerance and possibly to drought by preventing the overaccumulation of sodium ions and disease resistance); |
|  |  |  |  | *Tc06v2_g000730*<br>Non-specific lipid-transfer protein C, cotyledon-specific isoform (734,925..735,747) ;<br>*Tc06v2_g001470*<br>Zinc finger C-x8-C-x5-C-x3-H type family protein isoform 3 (1,516,921..1,523,842) ;<br>*Tc06v2_g001910*<br>Tetratricopeptide repeat-like superfamily protein, putative (2,025,081..2,027,186) ;<br>*Tc06v2_g002560*<br>VQ motif-containing protein 11 (2,679,449..2,680,592) | Lipid transport;<br>DNA binding, embryo development ending in seed dormancy;<br>Embryo development ending in seed dormancy;<br>Endosperm development, regulation of seed growth; |
| Pod index | TcSNP642 | 8 | 16,399,147 | *Tc08v2_g013700*<br>Tetratricopeptide repeat-containing protein, putative isoform 1 (14,427,929..14,450,269);<br>*Tc08v2_g014110*<br>Pentatricopeptide repeat-containing protein At1g71460, chloroplastic (14,862,384..14,867,389);<br>*Tc08v2_g015360*<br>Protein disulfide-isomerase 5-2 (16,765,487..16,772,516) | Embryo development ending in seed dormancy;<br>Embryo development ending in seed dormancy;<br>May be required for proper pollen development, ovule fertilization and embryo development |
| Seed length | TcSNP184 | 9 | 10,934,938 | *Tc09v2_g013350*<br>WAT1-related protein At3g30340 (9,427,175- 9,429,179);<br>*Tc09v2_g013720*<br>Probable E3 ubiquitin-protein ligase XBOS32 (10,304,321..10,313,838)<br>*Tc09cons_t014920.1*<br>Zinc finger protein CONSTANS-LIKE 5 (10,934,603..10,936,572) ; | Seed development, L-glutamate transport across plasma membrane;<br>Seed development;<br>Regulation of flower development and regulation of transcription; |
|  |  |  |  | *Tc09v2_g014390*<br>Probable xyloglucan endotransglucosylase/hydrolase protein 23<br>(11,778,780..11,780,085) | Carbohydrate metabolic process in *Gossypium hirsutum*, a member of the Malvaceae family as is *T. cacao* L. |

\*MTAs based on MLM with 612 filtered SNPs are denoted by an asterisk (\*)
Other MTAs are based on MLM with 737 or 836 unfiltered SNPs (such as in the case of TcSNP180 and seed number on chromosome 6, TcSNP642 and pod index on chromosome 8 and TcSNP184 and seed length on chromosome 9).
There are 109 putative candidate genes associated with cacao seed development. Thirteen were identified in this study (https://www.uniprot.org/uniprot/?query=cocoa+seed+development&sort=score)
Approximately 42 putative candidate genes that encode embryo and seed development, protein synthesis, carbohydrate transport and lipid biosynthesis, metabolism and transport were identified in this study

Associations between seed length and TcSNP 953 (*P* ≤ 6.98 × 10^-04^), pod index and TcSNP 667 (*P* ≤ 4.15 × 10^-04^), pod index and TcSNP 555 (*P* ≤ 3.85 × 10^-04^) and seed number and TcSNP 1160 (*P* ≤ 1.35 × 10^-04^), all on chromosome 4 (Table 6), although below the prescribed level of significance with Bonferroni correction, suggest that chromosome 4 may contain a cluster or ‘hotspot’ of QTLs for yield-related traits. There were two putative candidate genes involved with seed development co-localised with TcSNP 344, six that encode for seed development that were co-localised with TcSNP 667, eight responsible for seed protein and development and sugar transport linked to TcSNP 555 and five with seed development functions co-localised with TcSNP 1160 (Table 7). In addition, seven genes involved with lipid formation and seed development were localised close to TcSNP 953 on chromosome 4.

It was noteworthy that when 612 filtered SNPs were employed for GWAS using the MLM, a minor association (at *P* ≤ 6.98 × 10^-04^), below the stringent level of significance with Bonferroni correction, was observed between log seed length and TcSNP 953 at position 2,822,152 on chromosome 4. The latter marker is located 3.7 Kb upstream of the gene that encodes acyl transferase-like protein At3g26840, which controls seed mass and oil content in Assembly cotton 46 (*Gossypium* spp.). At3g26840 is involved in seed storage (globulins) and seed size in Assembly cotton 46 (Jako et al. 2001) [90]. *Gossypium* spp. are related to cacao, both being members of the Malvaceae family.

On chromosome 5, seven putative candidate genes were co-localised with TcSNP 733. These were all involved in seed development. One candidate gene encodes soluble inorganic pyrophosphatase 4, which is important for development, but also in stress resistance (including to cadmium ion response) in plants (https://www.uniprot.org/uniprot/Q9LFF9). The latter was not of functional significance in this study, but is important for optimised cocoa production systems. In addition, three functional candidate genes involved with seed development, sugar transport and fruit development were found to be co-localised with TcSNP 1110, and one involved with seed development was co-localised with TcSNP 823 on chromosome 5 (Table 7).

Five putative candidate genes, including a lipid transfer protein, were identified on chromosome 6, co-localised with a MTA involving TcSNP 180. Three putative candidate genes were found on chromosome 7, one of which was a sugar transporter. Likewise, three functional candidate genes with seed development roles were detected on chromosome 8 although the MTA, involving pod index, was below the stringent Bonferroni threshold (*P* ≤ 2.30 × 10^-04^) (Table 7).

On chromosome 9, TcSNP 184 was co-localized with four putative candidate genes, one of which encodes Zinc finger protein CONSTANS-LIKE 5 (Table 7). The latter is involved in the regulation of flower development and regulation of transcription (https://www.uniprot.org/uniprot/Q9FHH8). Other genes detected were all involved in seed development.

It was notable that seed length, seed length to width ratio and seed number were significantly associated with markers on different chromosomes. This suggests putative oligogenic or polygenic inheritance of these yield-related traits.

In addition, seed number and orbicular and oblate fruit shapes were significantly associated with TcSNP 390 on chromosome 7, a possible indication of gene linkage or perhaps pleiotropy. This requires further investigation.

It is also significant that TcSNP 401 (20,485,872), located 199 Kb upstream of *Tc00_t058610*, which encodes a putative MYB-related protein 308, 2 Mb upstream of the gene, *Tc04v2_g008890*, which encodes a putative MYB family transcription factor, and 1.3 Mb upstream of the gene, *Tc04v2_g009300*, which encodes a MYB domain protein 20, was significantly associated with fruit surface (ridge) anthocyanin intensity on chromosome 4. These associations were similar to those found previously by Marcano et al. [50] and Motamayor et al. [89]. A significant association (*P* ≤ 10 × 4.57^-05^) was also found between fruit surface anthocyanin intensity and SNP 644 on chromosome 9.

Other interesting results were obtained when MLM was performed with 737 (unfiltered) SNPs. There was a significant (*P* ≤ 3.81 × 10^-05^) MTA, which was detected between sepal length and TcSNP 1334 on chromosome 3. Furthermore, an association between TcSNP 180 and seed number on chromosome 6, when MLM was performed with 836 unfiltered SNPs, may be of interest despite not being considered stable or robust. TcSNP 180 was co-localized with genes encoding seed development as well as drought tolerance and disease resistance (Table 7). The observation regarding drought tolerance is of considerable significance since the conditions at the ICGT, where these accessions were observed, are considered sub-optimal in terms of soil moisture content [37].

Of the 17 significant MTAs unravelled when MLM with 612 (filtered) SNPS was performed, there were 2 for fruit shape oblate (an uncommon phenotype in this diverse cacao germplasm sample), and orbicular on chromosome 3. These MTAs involved TcSNPs 1353 and 1477. The oblate shape is a trait associated with certain wild types, which have evolved over a long period of time. A well-known accession with this trait is CATONGO. Interestingly, loci controlling fruit shape were dispersed over several chromosomes, representing independent linkage groups, when MLM was performed using 737 (unfiltered) SNPs.

The highly significant MTAs involving quantitative yield-related traits, observed in this study, suggest stability of genomic regions involved, as was also reported by Marcano et al. [50]. However, the genetic markers, co-localized with genes, were not major (accounting for ≥ 20% of the phenotypic variation expressed) since 5 to 17 percent of the phenotypic variation expressed was explained by the marker effect (R^2^).

In summary, about 40 putative candidate genes of functional importance were identified in this study. These included those that encode protein precursors, carbohydrate transport, lipid synthesis/bioassembly, binding and metabolism, lipid transfer and seed storage, endosperm development and regulation of seed growth, embryo development leading to seed dormancy, seed development/morphogenesis and regulation of flower and pollen development and ovule fertilization as well as responses to water deprivation, cadmium contamination and other abiotic stresses and biotic (disease) stresses (Table 7).

### Genomic prediction value of traits

The predictive (GEBV) values of the phenotypic traits studied are presented in Table 8. Of the qualitative traits studied, fruit basal constriction and filament anthocyanin intensity had predictive values greater than 0.5. The quantitative traits, seed number, seed mass, seed length, seed width, seed length to width ratio, pod index, fruit wall hardness, ovule number and fruit width had GEBV values greater than 0.5. The detection of several markers associated with yield-related traits with good predictive value, in this study, could facilitate genomic selection and marker-assisted selection in cacao. The yield-related traits, seed number and dried seed (cotyledon) mass had GEBV values of 0.611 and 0.6014, respectively. The GEBV value of cotyledon length and width, indicators of seed size, were 0.6199 and 0.5435, respectively. Interestingly, ovule number had the largest GEBV value of 0.6325. It is regarded as a reliable predictor of seed number, which is dependent on successful pollination.

**Table 8.** Predictive values (GEBV) of phenotypic traits associated with SNPS.

| Phenotypic Trait | GEBV |
| --- | --- |
| Ovule number | 0.6325 |
| Seed/bean number | 0.6110 |
| Seed/bean mass (g) | 0.6014 |
| Seed length (cm) | 0.6199 |
| Pod width (cm) | 0.5849 |
| Total wet seed mass (g) | 0.5656 |
| Seed width (cm) | 0.5435 |
| Seed length to width ratio (log transformed) | 0.5503 |
| Filament anthocyanin intensity | 0.5486 |
| Pod Index (log transformed) | 0.5394 |
| Pod basal constriction | 0.5116 |
| Pod index (not transformed) | 0.4975 |
| Pod wall hardness | 0.5134 |
| Sepal width (mm) | 0.4919 |
| Sepal length (mm) | 0.4366 |
| Ligule width (mm) | 0.4720 |

GEBV indicates the correlation between the predicted value per trait based on genotype and the observed value based on the measured phenotype

## Discussion

It is noteworthy that more than 17 potentially useful MTAs were detected in this study. Several hundred marker trait associations or QTLs have previously been identified in cacao. Allegre et al. [33] referred to 300 of these associations. In addition, Motilal et al. [91] identified a QTL for resistance to *P. palmivora* close to the region identified by Clément et al. [10] on chromosome 4. Queiroz et al. [92] identified a major QTL linked to resistance to Witches’ Broom disease. Royaert et al. [93, 94] identified marker-trait associations for self-compatibility and resistance to Witches’ Broom, respectively, in a segregating mapping population of cacao. Sounigo et al. [95] found several associations related to SSRs and yield. Motamayor et al. [89] identified candidate genes regulating fruit colour. Da Silva et al. [96] identified markers on chromosome 4, which were putatively co-localized with a major gene encoding self-incompatibility. Osorio-Guarín et al. [97] detected two genes putatively associated with productivity (number of healthy fruits) and seven encoding Frosty Pod disease resistance.

It must be borne in mind that genetic variation of quantitative (polygenic, continuous) traits such as yield and disease resistance are controlled by the combined effects of QTL, epistasis (interactions between QTLs) [14], the environment and interaction between environment and QTL [98]. The use of only biallelic subsets of SNPs, in this study, could have excluded multiallelic loci, which may have contributed to additional variance expressed in the study population for polygenic traits, including those related to yield potential. Mir et al. [99] described yield as a very complex quantitative trait that is controlled by a network of a “large number of small effect minor genes or QTLs”. For such polygenic traits, with small effect size, increasing the sample size of the study population and densely sampling a population that shows phenotypic diversity should improve the power to detect meaningful associations [51]. However, the relatively small effect size of the markers associated with traits, in this study, where none of the markers explained more than 20% of the phenotypic variation expressed, is not unusual for quantitative traits. Most of the markers studied explained 5 to 11 % of the phenotypic variation expressed. TcSNP 1335, on chromosome 7, explained 17% of variation expressed for seed number.

Significant associations found in this study between certain traits, such as fruit anthocyanin intensity, shape and seed length to width ratio, seed number, and loci on different chromosomes (Table 6), may be explained by the fact that a large part of trait variance was explained by several marker-trait associations, as described by Semagn et al. [98]. There were 26 SNPs significantly associated with fruit shape, two on chromosome 1, seven on chromosome 2, nine on chromosome 3, three on chromosome 5, two on chromosome 7 and three on chromosome 8, based on the results of MLM using 737 unfiltered SNPs. It seems justifiable to hypothesize that minor genes as well as major genes affect fruit shape.

The presence of markers significantly associated with different traits, in the same genomic region, was also observed in this study. The traits were seed number and orbicular shape on chromosome 7 (SNP 390) and seed number and seed length, also on chromosome 7 (SNP 1335) (Tables 6 and 7). These associations may indicate co-localization of the respective markers with a gene or gene block with pleiotropic effect [100] or may represent the phenomenon of linked genes, each one coded separately for a specific trait, as described by Araújo et al. [11]. In the case of seed number and seed length, indicators of seed size, this putative linkage is noteworthy since pod index (a measure of yield potential) is derived from seed number and mass. The relevant associated putative genes may have adaptive influence due to linkage mediated by selective forces, as explained by Yeaman [101]. The likelihood of such a phenomenon being observed during this study was feasible due to the inclusion of at least 48 cultivated accessions, including 28 Imperial College Selections (ICS) (S1 Table). The latter reportedly evolved over a period of more than two hundred years in Trinidad and Tobago and were selected based on large seed size and seed number and favourable yield, among other selection criteria [70]. It must be noted that pleiotropic markers may facilitate simultaneous selection of the multiple traits with which they are significantly associated and thus gene pyramiding.

### Putative candidate genes for yield-related traits

The storage compounds of cacao seeds are starch, lipids (fats) and storage proteins [102]. Bucheli et al. [103] investigated the variation of sugars, carboxylic acids, purine alkaloids, fatty acids, and endoproteinase activity during maturation of cacao seeds. Aspartic endoproteinase activity was observed to increase rapidly during seed expansion and a major change in the fatty acid composition occurred in the young embryo. Mustiga et al. [104] detected a major QTL explaining 24% of the relative level of palmitic acid on the distal end of chromosome 4, located close to the *Thecc1EG017405* gene. The latter is an orthologue and isoform of the stearoyl-acyl carrier protein (ACP) desaturase (SAD) gene that is involved in fatty acid biosynthesis.

Cacao seeds also contain a vicilin-like globulin, a seed storage protein [105]. It is noteworthy that TcSNP 555 on chromosome 4 was co-localised with vicilin in this study.

There are three acyltransferases and a phosphohydrolase involved in the bioassembly of plant storage lipids, *viz*., glycerol-3-phosphate acyltransferase (GPAT), lyso-phosphatidic acid acyltransferase (LPAT), diacylglycerol acyltransferase (DGAT) and phosphatidate phosphohydrolase (PAPase) [90]. Fritz et al. [106] purified glycerol-3-phosphate acyltransferase from the post-microsomal supernatant of cocoa seeds.

Triacylglycerols (TAGs) are the major storage lipids in several plants, and serve as energy reserves in seeds that are later used for germination and seedling development [107, 108]. The terminal step in TAG formation in plants involves the catalytic action of diacylglycerol acyltransferase (DGAT) in the presence of acyl-CoA [107]. Developing seeds in *Brassica napus* have been reported to produce Diaylglycerol (DAG) during the active phase of oil accumulation [109].

The proteins encoded by candidate genes, which were co-localized with SNPs found to be significantly associated with yield-related traits, during this study, are presented in Tables 6 and 7. An association (just below the stringent significance level with Bonferroni correction) observed between seed length and TcSNP 953 on chromosome 4, at a position of 2,822,152 bp, 3.7 Kb upstream of a gene that encodes diacylglycerol acyltransferase (Acyltransferase-like protein At3g26840%2C chloroplastic), is among the most noteworthy of this study and warrants further investigation. Another putative candidate gene, unravelled during this study, was Tc01v2_g022850, which encodes Bifunctional inhibitor/lipid-transfer protein/seed storage 2S albumin superfamily protein involved in Glycerolipid metabolism (chromosome 1, 2.5 Mb downstream of SNP 785 (31,785,158)) (Table 7). The results presented in Table 7 justify further investigation into the putatively functional roles of genes on chromosomes 1, 3, 4, 5, 7 and 9 in determining seed size and mass, components of yield potential in cacao.

### Consensus MTAs involving yield-related traits

Significant (stable) associations between yield-related traits, seed length and seed length to width ratio, seed number and seed mass and pod index, and SNPs were found on chromosomes 1, 3, 4, 5 and 7 in this study (Table 7). Marcano et al. [50] found 8 significant associations between SSR markers and yield-related traits. These included associations with fresh seed mass on chromosomes **1**, 2, **5**, 6, 9 and 10, MTAs involving dried seed mass (100 seeds) on chromosomes 2, **4**, 9 and 10 as well as one marker associated with seed number per fruit on chromosome **5**. In their mapping study, dos Santos et al. [69] identified several QTLs flanked by the markers *Tcm004s00289192*, *Tcm004s00615809,* and *Tcm004s01127580* on chromosome **4**, which were associated with pod index, dried individual seed mass, number of fruits harvested and number of healthy fruits harvested. They also identified a significant association between the marker, *Tcm002s23708704,* and pod index on chromosome 2. The study of dos Santos et al. [69] unravelled 13 candidate genes linked to yield (dried seed mass, pod number), on chromosomes 4 and 2. Nine of these genes are annotated as transmembrane transporters, specializing in sugar transport, two genes are involved in carbohydrate metabolism, one gene is involved in lipid metabolism, and one gene is involved in glucose metabolism [69]. Motilal et al. [87] identified three SNPs (TcSNP368, 697, 1370) on chromosomes **1** and 9 that were significantly associated with seed number. Clément et al. [20] found two QTLs for yield in the clone, POUND 12, located close to a QTL for yield, identified in IMC 78, on chromosome **4**.

Previous studies thus commonly observed loci on chromosomes **1, 2, 4, and 9** to be associated with seed mass and dimensions in cacao [50, 69]. The findings of this study suggest that yield-related traits are associated with loci on chromosomes 1, 3, 4, 5, 6, 7, 8 and 9 (Table 7), putatively linked to functional genes. However, not all of these MTAs were highly significant. Two SNPs, TcSNP 344 and 953, were associated with seed length on chromosome 4 (Tables 6 and 7). TcSNP 953 is located at the top of chromosome 4 (Fig 6) and thus the MTA, though below the stringent Bonferroni level of significance, may be considered validated since it was observed in a common region where yield-related MTAs were located in the studies by dos Santos Fernandes et al. [69] and Clément et al. [20].

**Fig 6.**
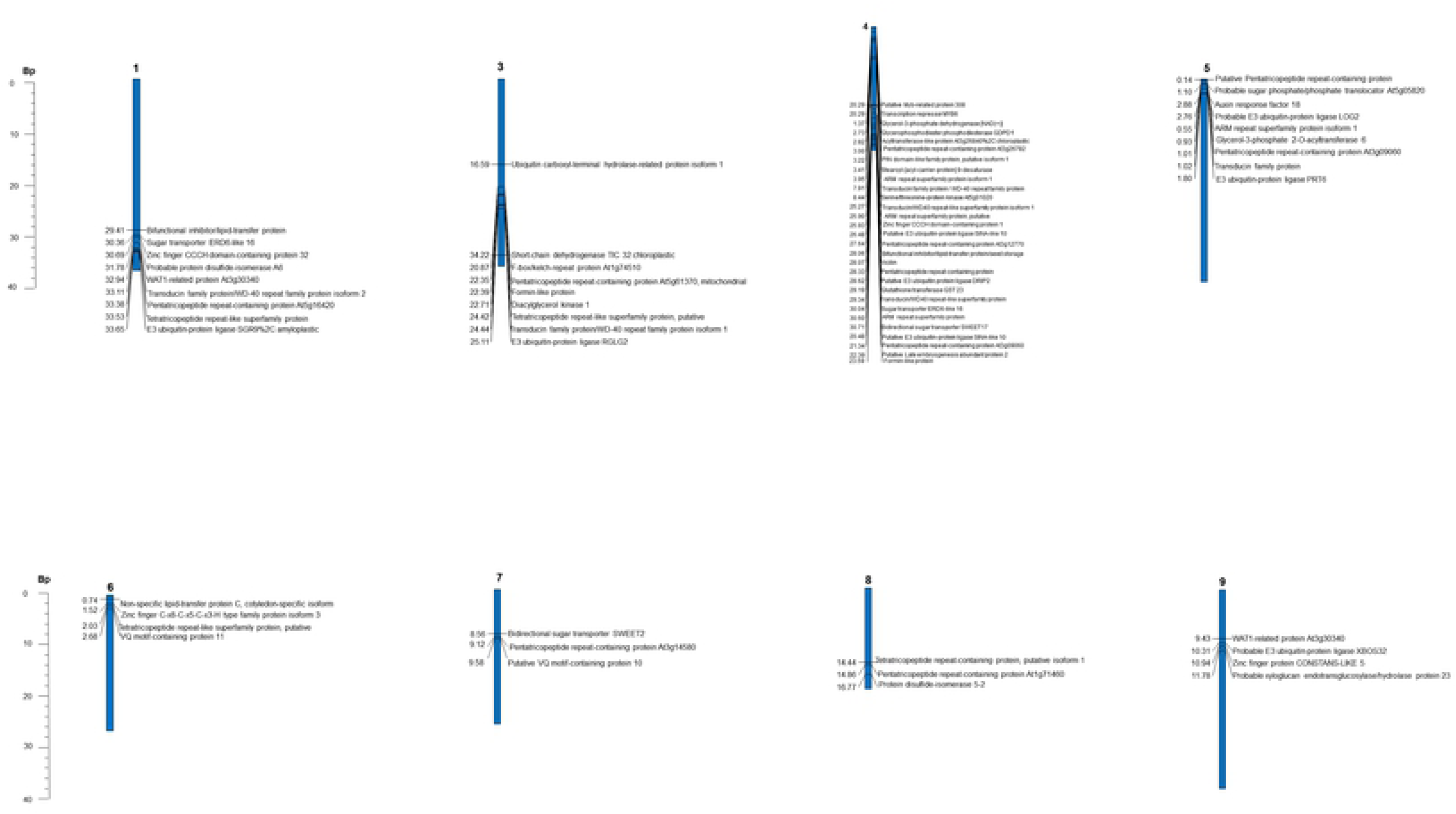
Genetic linkage map of *T. cacao* L. showing candidate genes co-localised with SNP markers associated with yield-related and other traits. Legend: Gene loci and proteins are shown on the right and genetic distances (Mb) are shown on the left. No candidate genes were identified on chromosomes 2 and 10

A highly significant MTA, observed in this study, involved seed number and TcSNP 785 on chromosome 1. Motilal et al. [87] also reported such an MTA on chromosome **1**. Marcano et al. [50] identified MTAs on chromosome 1 involving fruit number, fresh seed mass per fruit, and seed length, width and thickness.

Marcano et al. [50] found that the ‘Criollo allele’ was favourable in 68% of the seed-marker associations studied, and inferred that the ‘Forastero allele’ may sometimes be favourable for these traits. Criollo, Forastero and Trinitario are widely recognised classes of cacao in the trade. The accessions in this study that displayed favourable yield potential were mainly Trinitario (cultivated germplasm) [36] and those with Criollo ancestry. This was due mainly to their large seed sizes (Table 9). This supports the deduction of Doebley et al. [110] that “cultivated species generally have larger fruits or seeds compared to their wild ancestors, indicating that fruit and seed size are major agronomic traits that have been selected in crops during their domestication.” However, several Upper Amazon Forastero types, in this study, also had favourable (low) pod index due to their large seed numbers (Table 9, S7A Fig and S7B Fig).

**Table 9.**
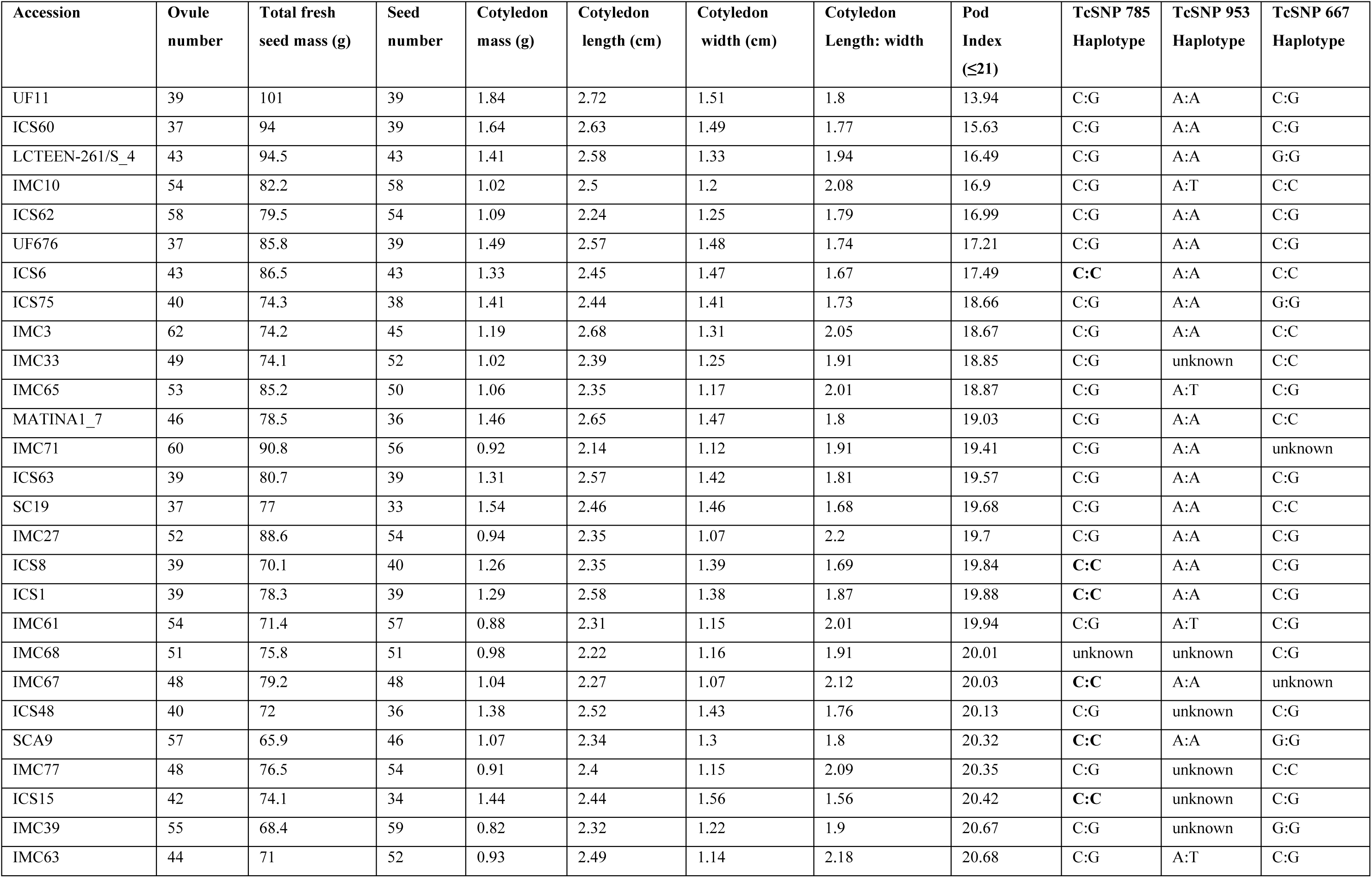
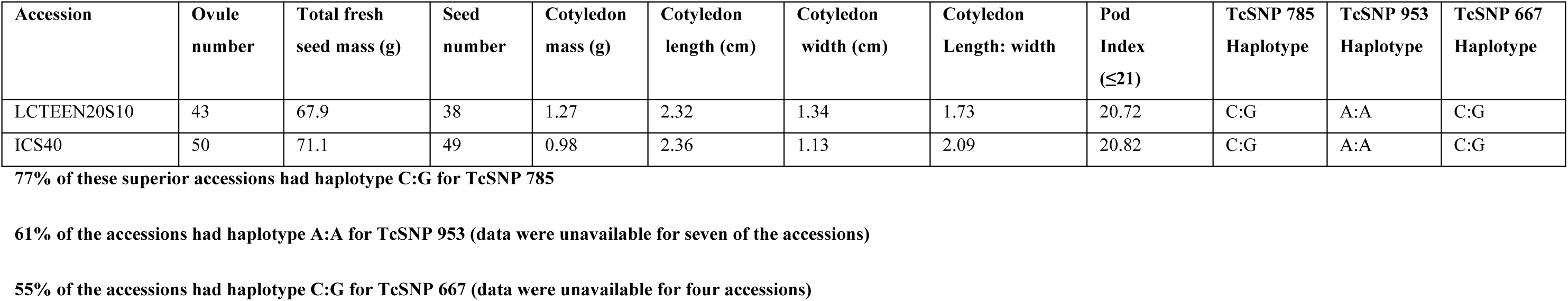
Superior accessions in terms of yield-related traits and associated haplotypes (allelic variants) for SNP markers of interest.

| Accession | Ovule number | Total fresh seed mass (g) | Seed number | Cotyledon mass (g) | Cotyledon length (cm) | Cotyledon width (cm) | Cotyledon Length: width | Pod Index (≤21) | TcSNP 785 Haplotype | TcSNP 953 Haplotype | TcSNP 667 Haplotype |
| --- | --- | --- | --- | --- | --- | --- | --- | --- | --- | --- | --- |
| UF11 | 39 | 101 | 39 | 1.84 | 2.72 | 1.51 | 1.8 | 13.94 | C:G | A:A | C:G |
| ICS60 | 37 | 94 | 39 | 1.64 | 2.63 | 1.49 | 1.77 | 15.63 | C:G | A:A | C:G |
| LCTEEN-261/S_4 | 43 | 94.5 | 43 | 1.41 | 2.58 | 1.33 | 1.94 | 16.49 | C:G | A:A | G:G |
| IMC10 | 54 | 82.2 | 58 | 1.02 | 2.5 | 1.2 | 2.08 | 16.9 | C:G | A:T | C:C |
| ICS62 | 58 | 79.5 | 54 | 1.09 | 2.24 | 1.25 | 1.79 | 16.99 | C:G | A:A | C:G |
| UF676 | 37 | 85.8 | 39 | 1.49 | 2.57 | 1.48 | 1.74 | 17.21 | C:G | A:A | C:G |
| ICS6 | 43 | 86.5 | 43 | 1.33 | 2.45 | 1.47 | 1.67 | 17.49 | C:C | A:A | C:C |
| ICS75 | 40 | 74.3 | 38 | 1.41 | 2.44 | 1.41 | 1.73 | 18.66 | C:G | A:A | G:G |
| IMC3 | 62 | 74.2 | 45 | 1.19 | 2.68 | 1.31 | 2.05 | 18.67 | C:G | A:A | C:C |
| IMC33 | 49 | 74.1 | 52 | 1.02 | 2.39 | 1.25 | 1.91 | 18.85 | C:G | unknown | C:C |
| IMC65 | 53 | 85.2 | 50 | 1.06 | 2.35 | 1.17 | 2.01 | 18.87 | C:G | A:T | C:G |
| MATINA1_7 | 46 | 78.5 | 36 | 1.46 | 2.65 | 1.47 | 1.8 | 19.03 | C:G | A:A | C:C |
| IMC71 | 60 | 90.8 | 56 | 0.92 | 2.14 | 1.12 | 1.91 | 19.41 | C:G | A:A | unknown |
| ICS63 | 39 | 80.7 | 39 | 1.31 | 2.57 | 1.42 | 1.81 | 19.57 | C:G | A:A | C:G |
| SC19 | 37 | 77 | 33 | 1.54 | 2.46 | 1.46 | 1.68 | 19.68 | C:G | A:A | C:C |
| IMC27 | 52 | 88.6 | 54 | 0.94 | 2.35 | 1.07 | 2.2 | 19.7 | C:G | A:A | C:G |
| ICS8 | 39 | 70.1 | 40 | 1.26 | 2.35 | 1.39 | 1.69 | 19.84 | C:C | A:A | C:G |
| ICS1 | 39 | 78.3 | 39 | 1.29 | 2.58 | 1.38 | 1.87 | 19.88 | C:C | A:A | C:G |
| IMC61 | 54 | 71.4 | 57 | 0.88 | 2.31 | 1.15 | 2.01 | 19.94 | C:G | A:T | C:G |
| IMC68 | 51 | 75.8 | 51 | 0.98 | 2.22 | 1.16 | 1.91 | 20.01 | unknown | unknown | C:G |
| IMC67 | 48 | 79.2 | 48 | 1.04 | 2.27 | 1.07 | 2.12 | 20.03 | C:C | A:A | unknown |
| ICS48 | 40 | 72 | 36 | 1.38 | 2.52 | 1.43 | 1.76 | 20.13 | C:G | unknown | C:G |
| SCA9 | 57 | 65.9 | 46 | 1.07 | 2.34 | 1.3 | 1.8 | 20.32 | C:C | A:A | G:G |
| IMC77 | 48 | 76.5 | 54 | 0.91 | 2.4 | 1.15 | 2.09 | 20.35 | C:G | unknown | C:C |
| ICS15 | 42 | 74.1 | 34 | 1.44 | 2.44 | 1.56 | 1.56 | 20.42 | C:C | unknown | C:G |
| IMC39 | 55 | 68.4 | 59 | 0.82 | 2.32 | 1.22 | 1.9 | 20.67 | C:G | unknown | G:G |
| IMC63 | 44 | 71 | 52 | 0.93 | 2.49 | 1.14 | 2.18 | 20.68 | C:G | A:T | C:G |

| Accession | Ovule number | Total fresh seed mass (g) | Seed number | Cotyledon mass (g) | Cotyledon length (cm) | Cotyledon width (cm) | Cotyledon Length: width | Pod Index ($\leq 21$) | TcSNP 785 Haplotype | TcSNP 953 Haplotype | TcSNP 667 Haplotype |
| --- | --- | --- | --- | --- | --- | --- | --- | --- | --- | --- | --- |
| LCTEEN20S10 | 43 | 67.9 | 38 | 1.27 | 2.32 | 1.34 | 1.73 | 20.72 | C:G | A:A | C:G |
| ICS40 | 50 | 71.1 | 49 | 0.98 | 2.36 | 1.13 | 2.09 | 20.82 | C:G | A:A | C:G |

- 1 77% of these superior accessions had haplotype C:G for TcSNP 785 - 2 61% of the accessions had haplotype A:A for TcSNP 953 (data were unavailable for seven of the accessions) - 3 55% of the accessions had haplotype C:G for TcSNP 667 (data were unavailable for four accessions) - 4

Based on the findings of this study, elucidation and selection of genotypes associated with large seed size in *T. cacao* L. may thus be facilitated by using TcSNP953 and TcSNP344 (Table 6 and 7). The results of a preliminary evaluation to detect genotypes associated with favourable yield potential (pod index) are presented in Table 9 and involve TcSNP 785, TcSNP 953 and TcSNP 667. Further investigation with training populations of *T. cacao* L. in genomic selection studies, as described by Bhat et al. [111], are recommended.

Studies are also recommended to further investigate functional genomics associated with yield-related traits in cacao such as was done by dos Santos et al. [69]. Such studies, in wheat, have revealed transcription factors, which can affect seed number, genes involved in metabolism or signalling of growth regulators, genes determining cell division and proliferation related to seed size, and floral regulators that regulate inflorescence architecture and seed number. Genes involved in carbohydrate metabolism, affecting plant architecture and grain yield such as trehalose phosphate synthase (TPS) and trehalose phosphate phosphatase (TPP) genes have also been identified [112].

Recommended follow-up studies would entail expression analysis, involving transcriptomics. The DGAT gene, Tag1, from *Arabidopsis* was shown to encode an acyl-CoA-dependent DGAT [90]. Jako et al. [90] demonstrated that seed-specific over-expression of the Diacylglycerol Acyltransferase (DGAT) cDNA in wild-type *Arabidopsis* “enhances oil deposition and average seed mass, which are correlated with DGAT transcript levels”, and that DGAT has an important role in regulating the quantity of seed triacylglycerols (TAGs), the sink size in developing seeds and thus seed size. They also demonstrated that “over-expression of the acyl-CoA-dependent DGAT in a seed-specific manner in wild-type *Arabidopsis* plants results in increased oil deposition and average seed mass.”

Ohto et al. [113] found that the gene APETALA2, which is a member of a large family of transcription factors, influences embryo, endosperm, and seed coat development and determines seed size in *Arabidopsis*. Kroj et al. [114] found the transcription factor, ABI3, to be implicated in seed maturation and in the expression of genes coding for seed storage proteins in *Arabidopsis*.

Seed size has also been shown to be directly determined by carbohydrate import into seeds, in maize and rice, and involves SWEET-mediated hexose transport [115]. SWEET genes regulate the transport, distribution and storage of carbohydrates in plants, and are involved in many important physiological processes, including phloem loading, reproductive development, disease-resistance, stress response, and host-pathogen interaction. In this study, SWEET17 was localized on chromosome 4 upstream of TcSNP555 (Table 7), which was associated with pod index (*P* ≤ 4.29 × 10^-4^). dos Santos et al. [69] identified a genomic region with copy-number variations of SWEET genes, also on chromosome 4, in their cacao QTL mapping study. In this study, SWEET 2 was also localized downstream of TcSNP1335, which was significantly associated (*P*≤1.15 ×10^-14^) with seed number as well as with log seed length (*P*≤ 6.75 × 10^-05^) on chromosome 7 (Table 6).

### Anthocyanin pigmentation

Marcano et al. [50] identified three regions associated with pigmentation on different organs in cacao. They considered the possible co-localization of markers related to pigmentation of structures especially in a small region of the chromosome 4, in which they found the SSR marker, mTcCIR115, to be located. This sector includes the major locus identified by Crouzillat et al. [12] as responsible for controlling ‘seed colour’ in the Catongo x POUND 12 backcross progeny.

Stack et al. [49] reported that cacao ‘fruit colour’ is considered to be controlled by a single gene localized within “a narrow genetic region with a strong phenotypic effect”. In this study, fruit ridge anthocyanin concentration (fruit colour) was significantly associated with TcSNP 401 on chromosome 4, located at 20,485,872, about 199 Kb upstream of the gene *Tc00_t058610* (mRNA), which encodes a Putative MYB-related protein 308 and 198 Kb upstream of the Transcription repressor MYB 6 (Table 7) (https://cocoa-genome-hub.southgreen.fr/). TcSNP 401 on chromosome 4accounted for 7.1% of the phenotypic variation in fruit colour observed (*P* ≤ 4.78 × 10^-06^) in this study (Table 6). Motamayor et al. [89] detected four SNP variants on chromosome 4, between 20,878,891 and 20,879,148 bp within a MYB transcription factor gene (TcMYB113), which were inferred to encode ‘fruit colour differences’ between cacao varieties.

Another significant association (*P* ≤ 6.79 × 10^-05^) for fruit surface anthocyanin intensity was found in this study when *TASSEL* MLM analysis was performed using 737 unfiltered SNPs. It involved TcSNP 1203, on chromosome 3 (located at 563,101), which accounted for 5.6 % of the phenotypic variation in fruit surface anthocyanin concentration in this cohort of germplasm.

MYB proteins are involved in regulatory networks controlling metabolism, including the synthesis of anthocyanins, responsible for the red pigmentation in cacao [116, 117]. Liu et al. [118] observed that overexpression of the Tc-MYBPA gene elicited increased expression of several genes encoding the major structural enzymes of the proanthocyanidin and anthocyanidin pathway in cacao.

There were also significant associations between filament anthocyanin concentration and SNPs on chromosomes 3 and 8 (TcSNPs 1183 and 1441, respectively) in this study (Table 6). All of the MTAs involving anthocyanin concentration in this study suggest multi-gene control of anthocyanin intensity in the mature fruit epidermis and flower filaments of cacao. Similarly, Marcano et al. [50] concluded that “biosynthesis of anthocyanins, involving several responsible enzymes, may produce a complex genetic system rather than one defined by a single gene.” Furthermore, the differential expression of pigmentation in the seeds and fruits of cacao, observed at the ICGT, as evidenced in the negative correlations in Table 3B, and as stated by Bartley [119], may be explained by the association of pigmentation of seeds and fruit pigmentation with several genomic regions. MTAs involving anthocyanin intensity warrant further investigation to determine their value for genomic selection due to the significance of this trait in differentiating among certain genotypes of interest, such as CCN-51 [89, 117], the nutraceutical value of anthocyanin and its putative role in cacao disease resistance [118].

## Future prospects

The results of this study appear to support the observation of Rockman [120] that most complex traits (such as those related to yield in cacao) are controlled by several (putatively interacting) loci with small effects. Some phenotypic traits are controlled by a small number of loci with large effects (as is often the case for traits under biotic selection) [51] while others may have more complex genetic architectures. The latter may be controlled by many rare variants, each having a large effect on the phenotype or conversely, many common variants with only small effects on the phenotypes, as described by Lee et al. [121]. The causative variants may be clustered in one or a small number of genes or across many genes. The data presented in Table 7 [on gene ontology] provide evidence of polygenic control of yield-related traits in cacao. For such traits, it may be more effective to predict the performance of genotypes by using multiple molecular markers [122]. Multilocus mixed linear models (MMLMs) may be considered for future studies in cacao when complex traits are investigated because these incorporate multiple markers simultaneously as covariates in a step-wise MLM [123].

The results from this GWAS were complementary to some from previous QTL, admixture and other association mapping studies in cacao. Markers identifying novel QTL in this study should be validated in the future. The use of a substantially larger body of reliable SNPs and an even larger subset of diverse cacao germplasm than that used in this study should be useful to further unravel useful MTAs in cacao.

It is also recommended that further research be undertaken to establish whether the trait associations revealed in this study and putatively linked specific gene variants have a functional role via gene expression and protein synthesis in *T. cacao* L.. Such studies were conducted by Bailey et al. [124], and Pokou et al. [125] for disease traits, Lanaud et al. [7] for self-compatibility and dos Santos et al. [69] for yield components, and by Chai et al. [126] in maize for seed oil content.

Rebbeck et al. [127] stated that “the lack of reproducibility of many association studies might reflect the number of studies that involve genetic variants with no functional significance.” It is noteworthy that several MTAs detected by this study were also found in previous ones on cacao and functional roles are expected.

Once the functional roles of putative genes co-localized with markers with significant associations to traits of interest are elucidated, the effects of relationships of these putative genes with geography and local adaptation must be established, as recommended by McKown et al. [128]. Micheli et al. [129] have reported on functional genomics in cacao focusing on genes expressed under specific physiological conditions. Consistency in QTL effects over different genetic backgrounds must also be established. Individuals with favourable marker genotypes and haplotypes may then be used as parental types for enhancement or breeding programmes in targeted environments.

Despite the fact that yield is a complex trait, our results on potential genomic selection (GS) for yield traits are very promising, given the high predictive values obtained for these traits, generally superior to 0.5. The slow progress hitherto realized in cocoa breeding [6] may be improved by the advancement made towards genomic prediction and selection in *T. cacao* L. [130, 35, 131].

Genomic selection predicts genotypic performance using genome-wide marker data [132, 111]. There are prospects for GS or marker-assisted selection (MAS) [133] in cacao, whereby numerous (tens of thousands) genetic markers covering the whole genome may be employed so that all QTL are in linkage disequilibrium with at least one marker. GS-MAS has been found useful for complex traits controlled by many QTL and with low effect and low heritabilities (*h*^2^) [134] once the markers with the most significant associations with traits are close to functional genes [46]. Accurate prediction of plant phenotype from genotype through GS-MAS should be facilitated by the utilization of wild cacao germplasm representing different genepools as sources of favourable alleles for traits of interest. It is noteworthy that Romero Navarro et al. [130] used the results of GWAS and genomic prediction to identify associated markers and develop predictive models for frosty pod rot and black pod diseases, as well as yield traits in a population of improved clones. Similarly, McElroy et al. [131] predicted resistance to *Moniliophthora* spp. diseases in three related populations of cacao using a 15K single nucleotide polymorphism (SNP) microarray for GWAS and genomic selection. They concluded that the “GS framework holds substantial promise in accelerating disease-resistance in cacao.” The results of this study on yield traits substantiate the value of this molecular breeding method to improve cocoa yield.

## Conclusion

In this study, carefully collated phenotyping data on traits of economic interest in cacao, such as yield potential, and SNP genotyping data, generated via transcriptome sequencing, were subjected to GWAS. A total of 421 cacao accessions were used for the GWAS. Thirty-one of these accessions represent promising material for breeding in terms of yield potential (Table 9). The goal was to use a large germplasm collection to decipher the genetic bases of yield traits and identify putative candidate genes linked to important phenotypic traits, as well as to simulate a genomic selection approach to evaluate its utility for cocoa breeding. By taking into account population structure and false discovery rates, genomic regions were found significantly associated with yield-related traits, fruit length, filament and fruit anthocyanin intensity and fruit shape.

The rather limited number of significant (stable) and robust associations (MTAs) detected in this study may be due to the prevalence of small effect size and rare genetic variants, which are not easily detected by GWAS. Genetic variants associated with complex traits, such as yield and disease resistance, are expected to have such small effects on function.

The results presented herein indicated oligogenic and polygenic control of yield-related traits in cacao. The stringently significant marker-trait associations related to yield, found in this study, were indications of the presence of quantitative trait loci on chromosomes such as 1, 3, 4, 5 and 7. They were validated by interval mapping analyses that found some corresponding QTL positions in other studies on chromosomes 1 and 4 [10, 20; 50; 87; 69]. Chromosome 4 putatively contains a QTL cluster associated with yield-related traits. This study may be unique in identifying putatively useful candidate genes, responsible for encoding yield-related traits via proteins involved in seed length and seed number determination, on chromosome 7 (Tables 6 and 7). Further studies for estimation of the functional effects of these putative candidate genes should be pursued.

A combination of genetics and functional genomics will facilitate understanding of gene function and gene interrelationships in cacao, as stated by Allegre et al. [33]. Fine mapping studies, such as that done for self-compatibility in cacao [7] and in cotton [135], will be useful especially since some putative candidate genes associated with seed development, seed lipid accumulation, metabolism and development and plant stress responses (including to drought and to cadmium) have been identified in this research and other studies in cacao. Genomic selection could be efficiently used to facilitate early selection of superior genotypes, using data from training and selected populations [136, 131]. The identification of yield-related traits with good predictive value, in the test population of this study, such as seed mass, number, length, width and length to width ratio as well as ovule number, could further facilitate genomic selection for yield potential in cacao. The performance of the non-phenotyped individuals at the Trinidad genebank (ICGT) could also thus be predicted if they are genotyped. This would be particularly useful for the enhanced genotypes (GEBP progeny) described by Bekele et al. [38]. Identification of parents possessing high predictive values and favourable alleles prior to crossing should prove beneficial for more rapid development of enhanced cacao progenies.

## Author contributions

Conceived and designed the experiments: CL and FLB

Performed the experiments: FLB, GGB, XA, MA, OF, MB

Analyzed the data: FLB, CL, IB

Wrote the manuscript: FLB, CL, XA, DS

All authors reviewed and approved the final manuscript

## Acknowledgements

The Director of CRC, Prof. Pathmanathan Umaharan, is gratefully acknowledged for endorsing this collaborative research. Useful discussions with Drs. Didier Clément, Christian Cilas and Martijn Ten Hoppen, CIRAD, France, Dr. Michelle End and Mr. R.A. (Tony) Lass, UK and Tricianna Maharaj, Trinidad, are deeply appreciated. J. Bhola, Dr. W. Mollineau, V. Badall, A. Richardson-Drakes, N. Persad, S. Samnarine, C. Jagroop, T. Jugmohan, E. Solozano and other individuals are gratefully recognized for technical assistance in phenotyping at various times during the period of study.

Financial support from the Government of Trinidad and Tobago, the Cocoa Research Association, UK and CIRAD, France that facilitated collation of the phenotypic and genotypic data, respectively, is gratefully acknowledged. However, the study design, conduct of this research and preparation of the manuscript were not influenced by the funding agencies.

## Supporting information

S1 Table. Background information on the *T. cacao* L. accessions used in the analyses.

S2A Table. Phenotypic data for 346 cacao accessions fully phenotyped and used to generate descriptive statistics.

S2B Table. Genotype data used in GWAS.

S3 Table. Results of tests of normality performed on the natural log transformed fruit and seed quantitative traits.

S4 Table. Coefficients of membership for clusters of accessions based on STRUCTURE analysis.

S5 Table. Distances over which linkage disequilibrium decayed to 50 percent over chromosomes 1, 4, 5, 7 and 9.

S6 Data. Summary of significantly positive marker-trait associations.

S7A Fig. Summary Report for Pod index in wild cacao.

S7B Fig. Summary Report for Pod index in cultivated cacao.

